# Gut feelings begin in childhood: how the gut metagenome links to early environment, caregiving, and behavior

**DOI:** 10.1101/568717

**Authors:** Jessica E Flannery, Keaton Stagaman, Adam R Burns, Roxana J Hickey, Leslie E Roos, Ryan J Giuliano, Philip A Fisher, Thomas J Sharpton

## Abstract

Psychosocial environments impact normative behavioral development in children, increasing the risk of problem behaviors and psychiatric disorders across the lifespan. Converging evidence demonstrates early normative development is affected by the gut microbiome, which itself can be altered by early psychosocial environments. Nevertheless, these relationships are poorly understood in childhood, particularly beyond peri- and postnatal microbial colonization. To determine the gut microbiome’s role in the associations between childhood adversity and behavioral development, we conducted a metagenomic investigation among cross-sectional sample of early school-aged children with a range of adverse experiences and caregiver stressors and relationships. Our results indicate that the taxonomic and functional composition of the gut microbiome links to behavioral dysregulation during a critical period of child development. Furthermore, our analysis reveals that both socioeconomic risk exposure and child behaviors associate with the relative abundances of specific taxa (e.g., *Bacteroides* and *Bifidobacterium* species) as well as functional modules encoded in their genomes (e.g., monoamine metabolism) that have been linked to cognition and health. We also identified heretofore novel linkages between gut microbiota, their functions, and behavior. These findings hold important translational implications for developmental psychology and microbiome sciences alike, as they suggest that caregiver behavior might mitigate the impact of socioeconomic risk on the microbiome and modify the relationship between subclinical symptoms of behavioral dysregulation and the gut microbiome in early school-aged children.

## INTRODUCTION

Childhood is a formative period of behavioral development that can influence the trajectory of psychiatric disorders and problem behaviors across the lifespan (1). Research has recently clarified the profound impact that a child’s economic, social, and caregiving environment plays in determining such outcomes (2, 3). For example, exposure to environmental risks early in life, such as growing up under low socioeconomic status (e.g., low income to needs ratio) or experiencing high family disruption and turmoil, can increase the risk for children to develop psychiatric disorders and associated problem behaviors (4). Caregivers, however, are one of the most proximal influences on and predictors of child wellbeing, and can modify how socioeconomic risk environments impact the child’s neurobiological and behavioral development (5). Across species, caregivers serve to protect their offsprings’ developing biology from environmental stressors and modify childhood behavioral response to adverse economic and social environments (3). Responsive and predictable caregiver behaviors are linked to improved child outcomes (6). Conversely, negative caregiver behaviors, such as perceived parental stress or disrupted parent-child relationships, can leave children more vulnerable to biological perturbations and behavioral dysregulation problems (7). Identifying early risk factors or correlates of childhood behavioral dysregulation is particularly important given that childhood is a time when mental health symptoms begin to emerge.

Ongoing research is focusing on understanding the underlying mechanisms by which adverse environments and caregiving behaviors (both positive and negative) influence a child’s behavioral development. Such research demonstrates that these factors can alter the developmental trajectory of central, autonomic and peripheral nervous systems function (8). These efforts have helped to influence not only the design of subsequent interventions (9), but also policy and practice (10).

Similarly, emerging research indicates that the gut microbiome may play a critical role in determining how a child’s environment ultimately impacts both their neurobiological function and mental health outcomes (11). The gut microbiome (hereafter “microbiome”) is the community of microbes that reside within the gastrointestinal tract and may be a key, yet relatively under-studied driver of neurobiological and behavioral development. Animal experiments demonstrate that the microbiome communicates with the central nervous system to influence social, explorative, and affective behavior through several pathways, including neuroendocrine and immune system coordination, vagal nerve stimulation, and neurotransmitter metabolism [see (12) for review of mechanisms]. These successional dynamics of the microbiome’s colonization are increasingly understood to interact with and shape the trajectory of neurobiological development (13). However, limited research has investigated the microbiome’s relationship to behavioral dysregulation and to key environments influences such as socioeconomic risk and caregiver behaviors during childhood (14–16).

Although we understand substantially less about the microbiome’s relationship with behavioral dysregulation early in life in humans, recent work links the composition of the microbiome to infant and toddler behaviors, such as surgency/extroversion, fear (15), and cognitive development (16). In addition, preliminary evidence from human studies of autism spectrum disorder suggests that the microbiome continues to play an active role in behavioral development following the first few years of initial gut colonization (17). It remains unclear if the microbiome associates with other forms of behavioral dysregulation and if it links to the onset of psychiatric disorders and problem behaviors. Understanding the link between the gut microbiome and subclinical behavioral dysregulation is particularly important given that normative behavior and behavioral disruptions develop throughout childhood, and that this period of development offers opportunities to intervene and treat disorders as they emerge.

Extensive evidence points to the microbiome’s sensitivity to psychosocial environments (18). Socioeconomic risk as well as caregiving behaviors during the initial programming of the microbiome can lead to long term changes in the microbiome and symptoms of behavioral dysregulation. For example, rodent pups that experienced an early life stressor of low resources, a model designed to mimic low socioeconomic status (SES), exhibited altered microbial compositions, increased intestinal permeability, and increased anxiety-like behaviors in adulthood relative to controls (19). Similarly, human adults from lower SES backgrounds exhibited lower microbial diversity (20).

Ample evidence suggests that caregiver behaviors influence the development of the microbiome and may modify how adverse environments impact the microbiome and subsequent childhood outcomes. In both humans and primates, prenatal physiological stress and a negative mother-infant relationship appear to reduce the level of Bifidobacteria and Lactobacilli in the infant’s microbiome (21, 22). Similarly, rodent pups exposed to repeated, prolonged maternal separation experience altered gut microbial profiles and increased intestinal permeability following social stressors in adulthood (23). The role of socioeconomic risk and caregiver behaviors on the developing microbiome remains notably understudied, and it is unclear if these relationships remain beyond the first few years of life.

Based on this prior research, we undertook an investigation of the microbiome’s link to socioeconomic risk, caregiving behaviors (both positive and negative), and child behavioral dysregulation. The goal of this study was to determine if and how the microbiome relates to environmental factors and behavioral dysregulation symptoms in early school-age children (4-7 years old; See Supplemental Table 1 for all sample metadata). Most studies of the microbiome’s relationship with behavioral dysregulation leverage 16S rRNA gene sequencing to determine how the taxonomic composition of the microbiome relates to environmental factors or behavioral problems and/or mental health outcomes in other populations (16). While this technique offers powerful insight into the kinds of taxa that constitute the microbiome, it offers no direct information about the functional mechanisms that they may utilize to respond to environmental factors or physiology.

In contrast, the approach employed in the present study, a technique known as shotgun metagenomics (24), alternatively involves simultaneously sequencing the genomes of taxa that compose the microbiome. In so doing, it offers insight not only into who resides in the gut, but also clarifies which functional pathways are encoded in their genomes. We generated shotgun metagenomic data from a cohort of children and determined how both the microbial taxa and the specific genetic functions they encode associate with subclinical child behavioral dysregulation symptoms (hereafter “behavioral dysregulation”), socioeconomic risk, and caregiver behaviors. We first tested if concurrent socioeconomic status influenced the child microbiome and whether self-reported parental behaviors statistically interacted with this association to explain additional variance. In addition, we examined how the child microbiome is associated with parent-reported child internalizing and externalizing behaviors, and whether caregiver behavior statistically moderated this association. Finally, we investigated if there were specific microbial taxa and metabolic pathways associated with different metrics of socioeconomic risk and child behavioral dysregulation. To our knowledge, this is the first study to assess the linkage between the microbiome, a child’s environment, and behavioral dysregulation symptoms during the 4-7 year old age range of formative behavioral and biological development. In so doing, this study reveals that exogenous factors including parental behavior impact the gut microbiome during the first few years of life, and that the microbiome indicates behavioral dysregulation, even at subclinical thresholds.

## RESULTS

In order to measure the microbiome, we collected stool from 40 children from a mid-size city in the Pacific Northwest that were already participating in a larger study (25). Parents of the children filled out questionnaires regarding five covariate categories: socioeconomic risk, behavioral dysregulation, caregiver behavior, demography, and gut-related history (i.e., factors known to influence microbiome composition such as antibiotic use). DNA was extracted from the fecal samples, sequencing libraries were prepared, and shotgun metagenomic sequencing was conducted according to standard protocols (see Methods). Unique metagenomic sequences were assigned, if possible, to the bacterial species level which resulted in 213 unique taxon assignments after quality control. Using these assignments, we estimated the taxonomic composition of the microbiome. Sequences were also assigned to molecular functional groups using the Kyoto Encyclopedia of Genes and Genomes (KEGG) database. These assignments are referred to as KEGG orthologs (KOs), and represent individual functions within larger genomic modules, which are components of functional pathways. The sequence set was assigned to 13,183 unique KOs, after quality control. Using these taxonomic and functional assignments, we constructed community tables (matrices of taxon or KO relative abundances by sample) to test associations between the microbiome and our covariates of interest in a statistically rigorous manner (see Methods and Supplemental Methods for specific details regarding participants, sample collection, molecular methods, and sequence analysis).

Because the questionnaires filled out by parents encompassed more potential covariates (52) than microbiome samples (*n* = 40), we began our analysis by selecting the covariates within each covariate category that explained a statistically significant amount of variance in the microbiome composition between samples (see Methods). This covariate selection process returned an identical set of ten covariates for both taxonomic and functional composition (See Table 1). In order to test our hypotheses that socioeconomic risk, behavioral dysregulation, and caregiver behavior covariates significantly associate with the composition of the microbiome, we utilized a constrained correspondence analysis (CCA) to create ordinations. This method is particularly appropriate for our study design because it accounts for the variance in the microbiome explained by factors that prior research indicates may have a strong effect on the composition of the microbiome, but which are not the direct focus of this research (i.e., demography and gut-related history). We then ran a permutational analysis of variance (PERMANOVA) on the remaining, unexplained variance to test the significance of the relationships between covariates and the composition of the microbiome. Selected covariates within each category (e.g., demography, gut-related history, child dysregulation behaviors, socioeconomic risk, and caregiver behavior) were determined by the *envfit* model. For each set of covariates, we tested their association with both the taxonomic (species) and functional (KO) composition of the microbiome.

**Table 1.** The set of covariates selected by *envfit* analysis for both taxonomic- and functional-based microbiome composition. All metrics are reported via questionnaire by the parent. PSI, Parenting Stress Index; LEC, Life Events Checklist; CBQ, Children’s Behavior Questionnaire; CBCL Child Behavior Checklist

| Covariate Category | Covariate | Description |
| --- | --- | --- |
| Parental Stress | PSI Parent-Child Dysfunction | Severity of dysfunctional interactions between parent and child |
| Socioeconomic Risk | LEC Poverty Related Events | Number of self-reported stressful life events associated to poverty |
|  | LEC Turmoil | Number of self-reported stressful life events associated with family turmoil |
|  | LEC Total | Total number of self-reported stressful life events |
|  | Income to Needs | Ratio of yearly household income to federal income needs per number of people supported by that income |
| Child Behavioral Symptoms | CBQ Impulsivity | Speed of response initiation |
|  | CBQ Inhibitory Control | The capacity to plan and to suppress inappropriate approach responses under instructions or in novel or uncertain situations |
|  | CBCL Depressive Problems | A subscale of the CBCL containing items associated with depression |
| Demography | Child Race/Ethnicity | Parent report of child's racial and ethnic background |
| Gut-related History | Geographic Locations | Lived in a different state or country >3 months |

### Microbiome Composition, Socioeconomic Risk, and Caregiver Behavior

We first examined whether metrics of socioeconomic risk and caregiver behavior significantly explain the observed variance in overall microbiome diversity and composition. In addition, we investigated whether these associations manifested at the level of the taxonomic identities of the microbiome constituents or the functional potential of the metagenome. We started by testing the associations between the taxonomic composition of the microbiome and the selected socioeconomic risk and caregiver behavior covariates. To maximize scientific rigor, we constructed a CCA model, which is based on a Euclidian distance, that first accounted for the selected demography (child ethnicity) and the selected gut-linked covariates (previously shown to influence gut [25–28]; geographic location) by determining the amount of variance explained. These two demography and gut history covariates accounted for 19.9% of the total variance in taxonomic composition. The socioeconomic risk and caregiver behavior covariates that remained in the best model (according to Akaike Information Criterion) explained a further 18.5% of the variance, leaving 61.6% of the variance unexplained. A PERMANOVA test on this CCA model revealed a single significant association. Specifically, the taxonomic composition of the gut microbiome taxonomic was associated with the selected socioeconomic risk covariate (number of family turmoil associated life events; F = 1.61, *p* = 0.0094; Fig 1A, Supplemental Table 2). Additionally, the selection of the best model included an interaction between Poverty Events (number of poverty-associated life events) and Parent-Child Dysfunction (parenting stress index subscale), but after controlling for gut-history and demography covariates, it was not significant (F = 1.39, *p* = 0.073).

**Figure 1.**
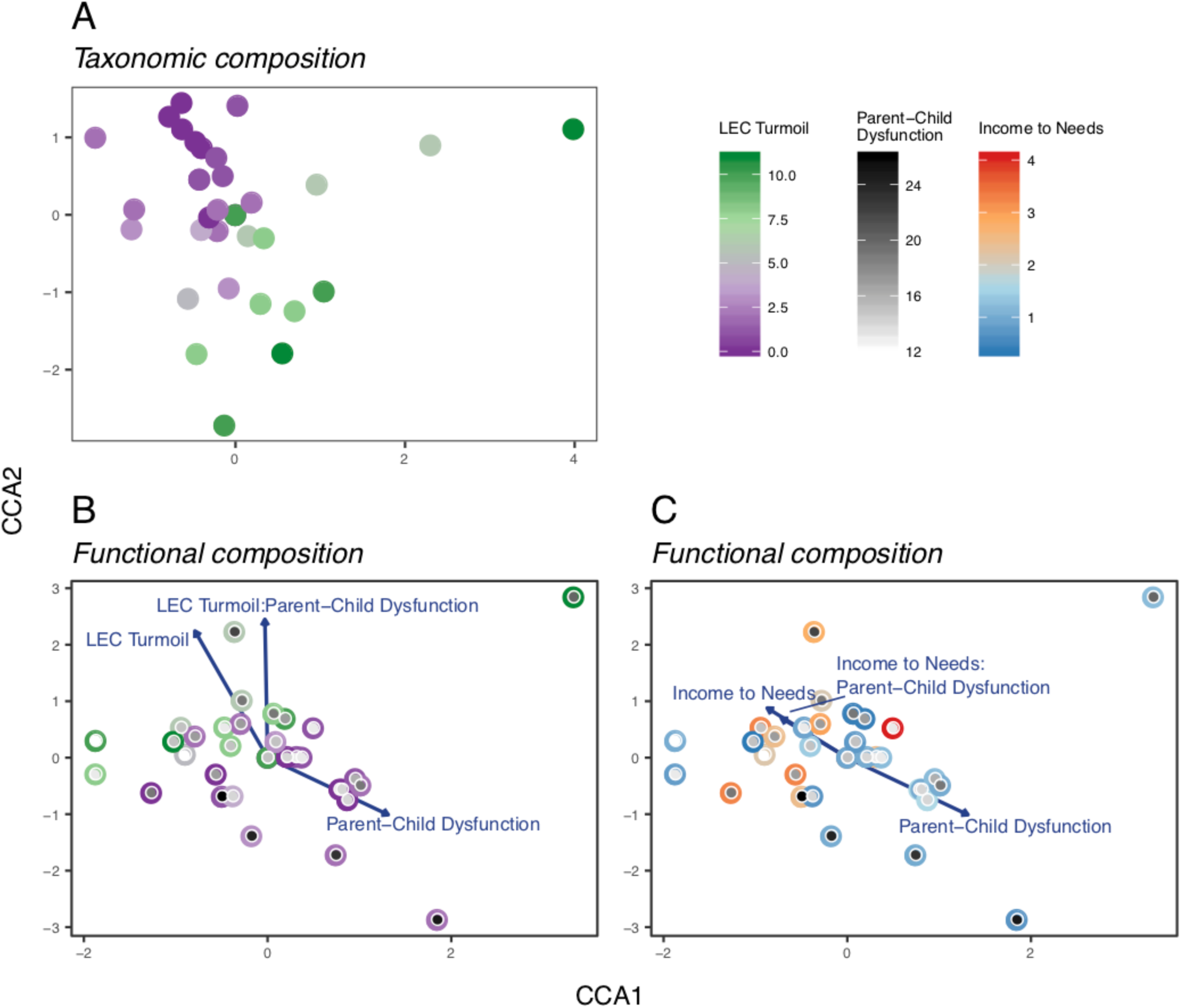
Constrained correspondence analysis (CCA) ordinations for taxonomic and functional composition of the microbiome and socioeconomic risk and caregiver behavior covariates. Only covariates that have significant main effects or are part of a significant interaction are depicted in each ordination. Significance was assessed by PERMANOVA (α = 0.05), see Supp. Tables 2 & 3 for statistical results. **(A)** Ordination of taxonomic (species-level) composition. Each point represents a sample in ordination space and is colored by LEC Turmoil Event score. **(B)** Ordination of functional (KO-level) composition. Each point represents a sample and consists of two parts: the color of the outer circle corresponds to the sample’s LEC Turmoil Event score; the inner circle is shaded from white to black indicating the sample’s Parent-Child Dysfunction score. **(C)** Ordination of functional (KO-level) composition, sample locations are identical to panel B. In this panel the outer circle of the point is colored according of the sample’s Income to Needs score. The inner circle is shaded identically to panel B.

As noted previously, the metagenomic (as opposed to amplicon-based) methodology we employed made it possible to test the associations between socioeconomic risk, caregiver behavior, and the *functional composition* of the microbiome. As in the prior analyses, we set the two demography and gut-related history covariates as conditional variables, which explained 25.7% of the total variance in functional composition. The socioeconomic risk and caregiver behavior covariates that remained in the best model accounted for 28.5% of the total variance in functional composition, while 45.8% remained unexplained. A PERMANOVA test on this model found that the caregiver covariate Parent-Child Dysfunction significantly interacted with both Turmoil Events (F = 2.51, *p* = 0.0097; Fig 1C) and Income to Needs Ratio (measure of poverty; F = 1.86, *p* = 0.041; Fig 1D, Supplemental Table 3). These results provide evidence that, in terms of the microbiome’s functional potential, caregiver behavior can moderate the associations between socioeconomic risk covariates and the microbiome.

### Microbiome Composition, Behavioral dysregulation, and Caregiver Behavior

In order to address our second question, whether metrics of behavioral dysregulation and caregiver behavior significantly explain the observed variance in overall microbiome diversity and composition, we applied the same analysis pipeline as above, substituting the selected child behavioral dysregulation symptom covariates for the socioeconomic risk covariates. The analysis of the taxonomic composition of the microbiome revealed no significant associations (Supplemental Table 4).

For the functional composition of the microbiome, we again found that the caregiver behavior covariate Parent-Child Dysfunction significantly interacted with two child behavioral dysregulation symptom covariates: depression (Depressive Problems; F = 2.69, *p* = 0.0079; Fig. 2A) and ability to inhibit impulses (Inhibitory Control; F = 2.18, *p* = 0.038; Fig. 2B, Supplemental Table 5). Again, these results provide evidence that the microbiome is associated with certain behavioral dysregulation, and that caregiver behavior may moderate these associations. For this particular study sample, however, the evidence suggests that it is the composition of functional groups within the microbiome, more so than the taxonomic composition of the microbiome, which correlates with behavioral dysregulation and caregiver behavior.

**Figure 2.**
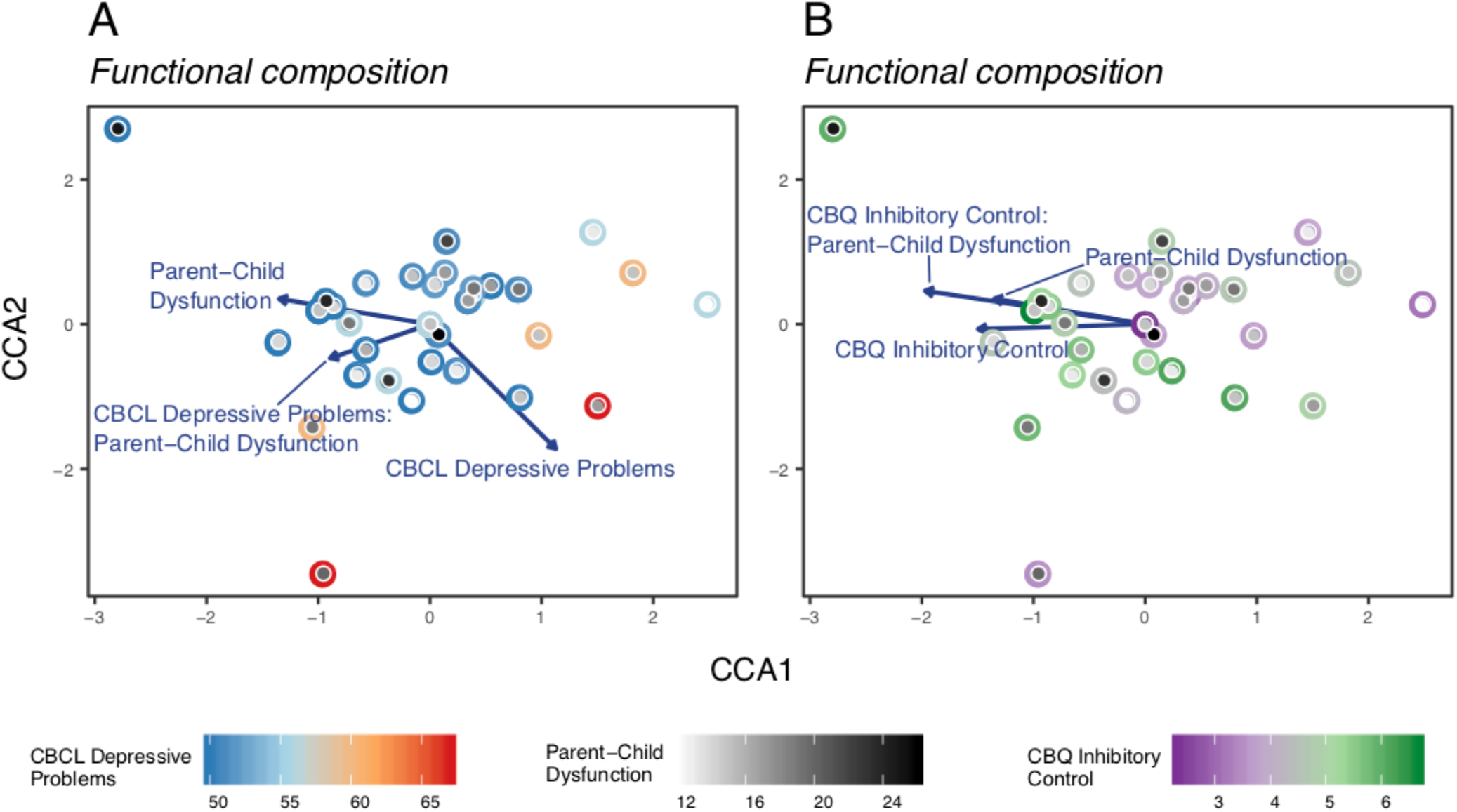
CCA ordinations for functional composition of the microbiome, behavioral dysregulation, and caregiver behavior covariates. Only covariates that have significant main effects or are part of a significant interaction are depicted in each ordination. Significance was assessed by PERMANOVA (α = 0.05), see Supp. Tables 4 & 5 for statistical results. **(A)** Ordination of functional (KO-level) composition. Each point represents a sample and consists of two parts: the color of the outer circle corresponds to the sample’s Depressive Problems score; the inner circle is shaded from white to black indicating the sample’s Parent-Child Dysfunction score. **(B)** Ordination of functional (KO-level) composition, sample locations are identical to panel A. In this panel the outer circle of the point is colored according of the sample’s Inhibitory Control score. The inner circle is shaded identically to panel A.

### Individual Taxa, KOs and Socioeconomic Risk, Child Behavioral Dysregulation Symptom Covariates

The above analyses assessed covariates of the overall composition and diversity of the gut microbiome. To obtain a finer resolution on the interactions between the gut microbiome, socioeconomic risk, and behavioral dysregulation, we employed a pairwise compound Poisson generalized linear models (CPGLM) to regress a specific taxon or KO relative abundance in the gut against each socioeconomic risk or behavioral dysregulation covariate. A comprehensive set of results of the pairwise relationships that maintained significance after false discovery rate (FDR) correction can be found in Supplemental Tables 6 & 7. Briefly, we found 67 significant pairwise relationships between covariates and taxa identified at the species level (48 for behavioral dysregulation, 19 for socioeconomic risk covariates; Fig. 3). For these taxon-covariate relationships, we found numerous associations involving butyrate-producing bacteria, and specific taxa of particular interest, including *Bacteroides fragilis* and *B. thetaiotaomicron*, which have demonstrated anti-inflammatory effects in mice and humans (30). We found significant relationships between 7 socioeconomic risk and 13 child behavioral dysregulation symptom covariates and 695 functions defined at the KO level. Of these 695 pairwise results, 94 KOs were grouped within defined metabolic modules (Fig. 4). Consistent with prior studies, for the KO-covariate relationships we found numerous associations involving monoamine metabolism (including tryptophan, tyrosine, glutamate, and leucine) and inter-microbe antagonism (type VI secretion systems and lipopeptide antibiotics).

**Figure 3.**
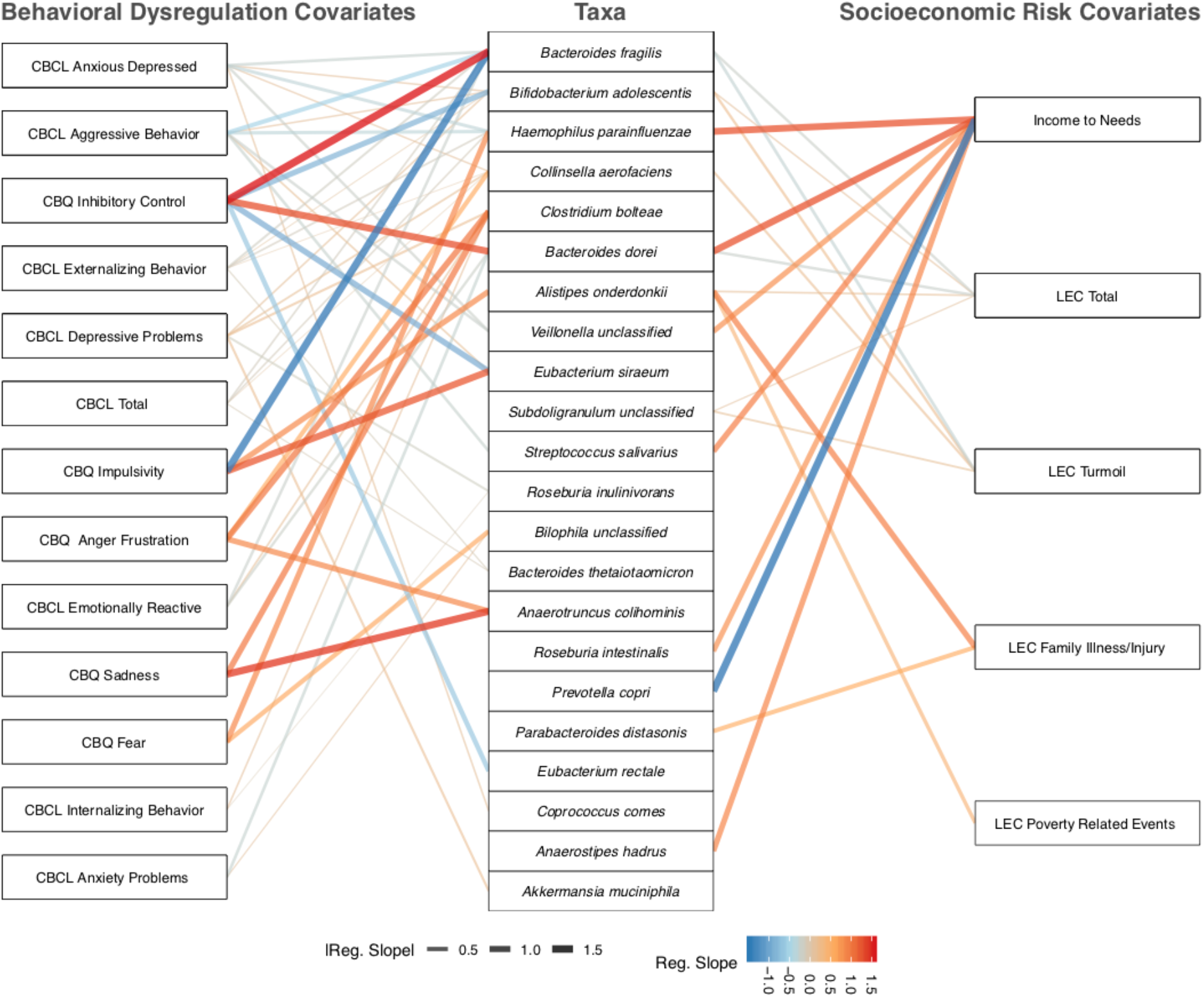
A network representing statistically significant pairwise associations, according to generalized linear models, between individual taxa and behavioral dysregulation or socioeconomic risk covariates. The left column shows individual behavioral dysregulation. The central column shows individual taxa identified to the species level. The right column shows individual socioeconomic risk covariates. Lines are only drawn between a covariate and a taxon if there is a significant relationship. The color of the line represents whether the association between the covariate and taxon is negative (blue) or positive (red). The width and intensity of the line color represents the slope of the regression line that describes the association (steeper regression lines are wider and brighter).

**Figure 4.**
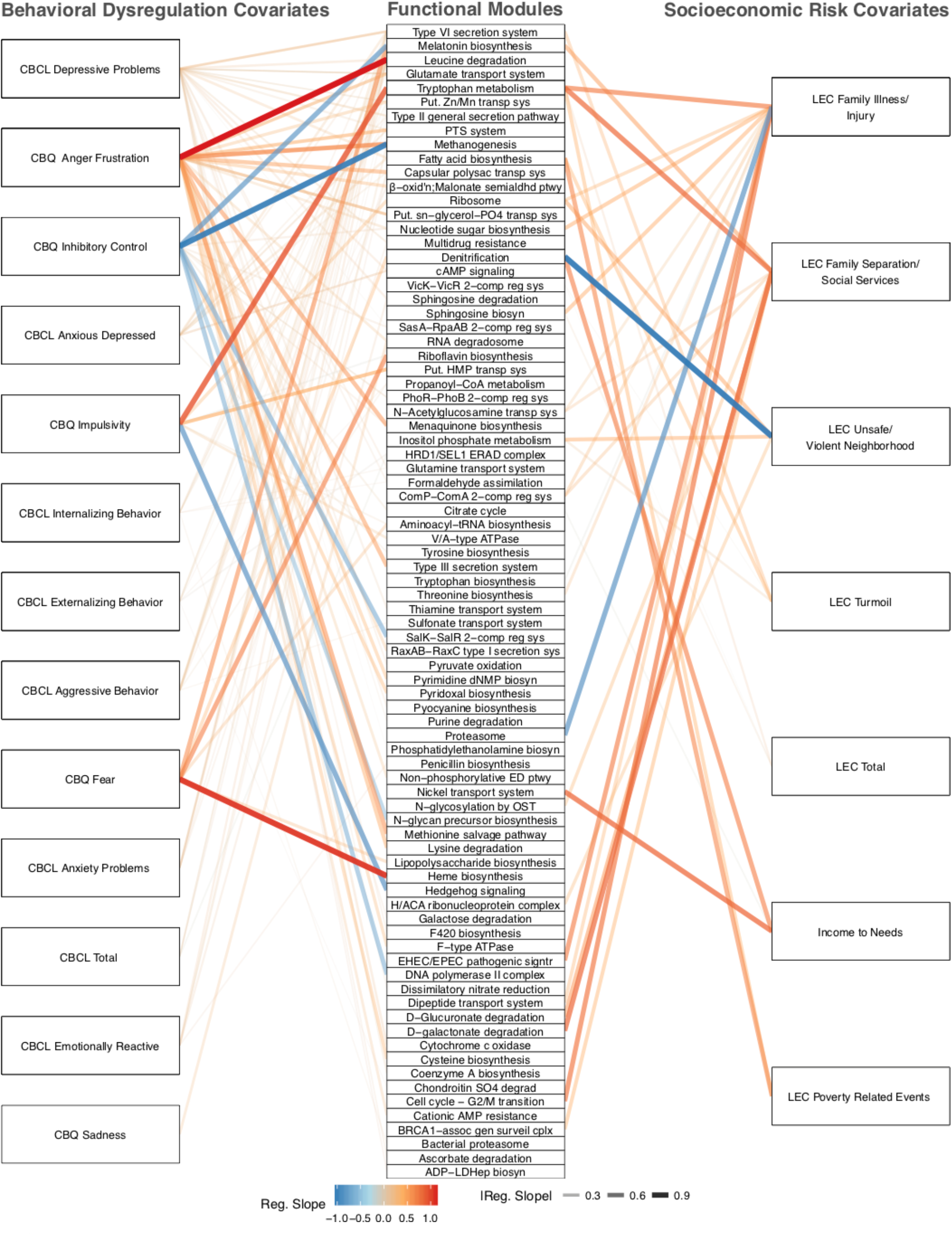
A network representing statistically significant pairwise associations, according to generalized linear models, between individual KOs (grouped into modules) and behavioral dysregulation or socioeconomic risk covariates. The left column shows individual behavioral dysregulation. The central column shows functional groups assigned at the KEGG module level. The right column shows individual socioeconomic risk covariates. Lines are only drawn between a covariate and a module if there is a significant relationship. The color of the line represents whether the association between the covariate and module is negative (blue) or positive (red). The width and intensity of the line color represents the slope of the regression line that describes the association (steeper regression lines are wider and brighter).

## DISCUSSION

The present study provides novel insights into the relationship between the gut microbiome and both the psychosocial environment and behavioral dysregulation in a cross-sectional sample of early school-aged children (Figure 5). Furthermore, this is the first study to assess if caregiving behaviors (i.e., perceived parental stress) can statistically modify the child’s gut microbiome’s association with their level socioeconomic risk exposure and behavioral dysregulation. As such, if replicated, the work provides a potentially new avenue of research into the mechanisms of behavioral intervention in future research. These results provide supportive evidence that the psychosocial environment continues to shape not only the taxonomic composition, but also the functional potential of the microbiome beyond initial gut microbial colonization that occurs in the perinatal period. Notably, the behavioral dysregulation symptoms measured in this study occurred at thresholds not necessarily indicative of psychiatric disorders of childhood. That these relationships were observed at subclinical levels of behavioral dysregulation symptoms suggests that the microbiome may play a role in the emergence of dysregulated behavior (i.e., providing early associative relationships prior to reaching clinical thresholds), rather than simply being present in clinical populations.

**Figure 5.**
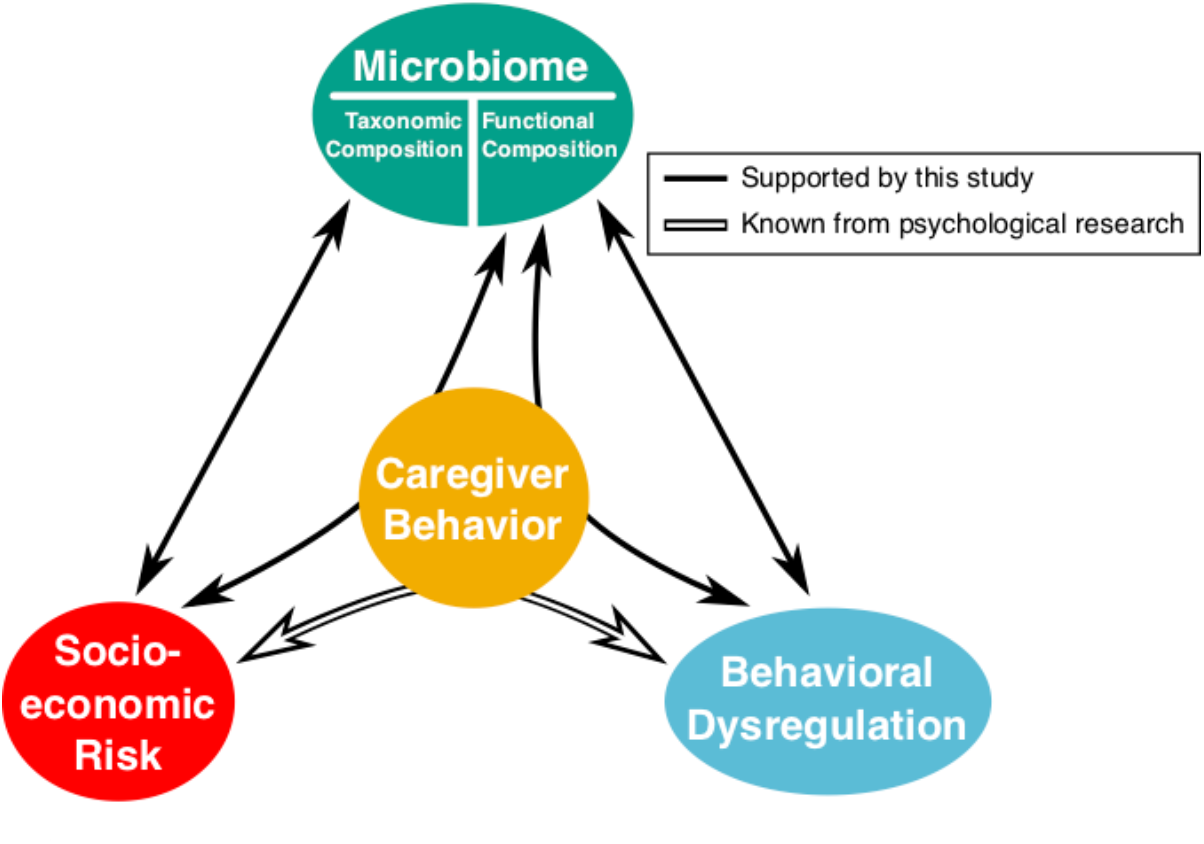
This figure illustrates the results of our hypothesis testing using ordination-based analyses. White solid arrows indicate relationships supported by evidence from prior psychological research. The black arrows represent relationship between the covariate categories and composition (taxonomic or functional) of the gut microbiome, as determined by our ordination- and PERMANOVA-based analysis (see Supp. Tables 2-5). Straight arrows represent significant main effects between the microbiome and a covariate category (e.g. between Socioeconomic Risk and taxonomic composition of the microbiome). Arrows that curve through Caregiver Behavior indicate that there is a significant interaction between Caregiver Behavior and the other covariate category (e.g. our analysis revealed two significant interactions between Socioeconomic Risk and Caregiver Behavior in their association with the functional composition of the microbiome).

Importantly, this study also documented that the quality of the caregiver-child relationship may moderate the microbiome’s association with both socioeconomic risk and behavioral dysregulation. Because our study relied on correlational methods, it is possible that is in fact the socioeconomic risk and the behavioral dysregulation symptoms that are moderating the association between the microbiome and caregiver behavior. However, given that none of the statistical tests support a significant main effect of caregiver behavior, but there is at least one significant association between both socioeconomic risk and behavioral dysregulation symptoms and the microbiome, it is plausible to conclude that the caregiver-child relationship may moderate the microbiome’s association with the other covariate groups rather than the other way around. That said, future work should seek to disentangle these relationships.

These findings have important implications for developmental psychological and developmental microbiome sciences alike, suggesting that the microbiome may be a pathway by which caregiver behavior may mitigate the impact of socioeconomic risk and influence early school-aged child outcomes. Caregivers may influence the microbiome in childhood through reducing or exacerbating their child’s experience of psychosocial stress. For example, increased parental stress that results from reduced economic or social support may in turn increase the child’s stress. Conversely, supportive parenting can reduce a child’s physiological and perceived stress, which may protect the microbiome from perturbations related to the physiology of stress. Furthermore, the metrics of caregiver behavior assessed in this study may be correlated with other symptoms of behavioral dysregulation, such as the home environment and diet of the child that may alter the composition and diversity of the gut microbiome (31, 32). Alteration of the child microbiome through the home environment and diet may in turn influence the level of physiological stress response in the child.

While we detected a significant association between parent-reported family turmoil and the taxonomic composition of the microbiome, we cannot conclude that caregiver behavior moderates this association, as there was no significant interaction between these variables. Due to our sample size, we cannot rule out that this study may have been underpowered to detect this relationship. However, as shown in Figures 1B & C (Supplemental Table 3), in terms of the functional composition of the microbiome, there were significant interactions between parent-reported parent-child dysfunction and two metrics of socioeconomic risk: income-to-needs ratio and family turmoil. Therefore, we found supportive evidence that the dysfunctionality of the parent-child relationship moderates the nature of the relationship between both economic and social forms of adversity and the functional potential of the gut microbiome of a child. This underscores the potential for caregivers to mitigate the impact of socioeconomic risk exposure on the developing gut microbiome. One mechanism by which socioeconomic risk may influence the microbiome is by exposing the child to different environmental microbes. For example, for modernized urban populations, there is evidence that greater socioeconomic status affords people the ability to travel away from human-dominated environments and gain exposure to microbe associated with the natural environment. Such differences in microbial exposure in early development associate with different profiles of immune function (33). Furthermore, adverse postnatal environments that are often comorbid with socioeconomic risk, such as frequent antibiotic use or toxin exposure, associate with altered microbial composition and intestinal permeability (34, 35). Future work should build upon these findings to test if the microbiome may serve as a mechanism by which economic and social adversity (socioeconomic risk) influences behavioral dysregulation.

When we tested whether the relationship between the gut microbiome and behavioral dysregulation depended on the parent-child relationship, our analyses only found significant interactions for the functional potential of the microbiome (Fig. 2A & B; Supplemental Table 5). In this case, the nature of the relationships between the functional microbiome and two behavioral dysregulation -- depressive problems and inhibitory control -- were modified by the quality of the parent-child relationship. This is consistent with prior literature that indicates behavioral dysregulation in childhood spans internalizing and externalizing dimensions (36). Again, the lack of any significant behavioral dysregulation for the taxonomic microbiome may indicate either that this study is underpowered at the taxonomic level or that these relationships are more dependent on the metabolic capabilities of the whole microbiome rather than attributes associated with specific taxa. In either case, it suggests that intervening to improve the parent-child relationship may influence the functional potential of the microbiome more strongly than its taxonomic composition. One proposed way that humans mothers may help regulate the gut-brain-microbiota axis is through skin-to-skin contact, particularly with high-risk infants (i.e., preterm; [31]. Future work should seek to tease apart the mechanisms by which parenting behaviors may influence the microbiome in later periods of development.

In addition to assessing the relationship between the selected covariates and the microbiome as a whole, we sought to understand such relationships at a finer scale of resolution. Therefore, we conducted pairwise comparisons between covariate scores and the abundances of each taxon and KO to determine whether there were specific relationships between socioeconomic risk or behavioral dysregulation and specific taxa or functions found to be important in previous studies (Figs. 3 & 4, Supplemental Tables 6 & 7). The taxon that associated with the greatest number of socioeconomic risk and behavioral dysregulation covariates was *Bacteroides fragilis*. Interestingly, *B. fragilis* associated with reduced levels of aggression, anxiety, emotional reactivity, externalizing behavior, impulsivity, and an increase in inhibitory control (i.e., better mental health). It was also associated with lower reported incidents of family turmoil. *B. fragilis* has been shown, in mice, to modulate the mammalian immune system and protect against pathogen-induced inflammation, specifically through the production of polysaccharide A (30, 37). *B. thetaiotaomicron* has also been shown to have anti-inflammatory effects in the mammalian intestine, and it too associates with decreases in externalizing behavior, as well as the overall score for negative behavioral dysregulation (38). Recent psychological research has provided strong evidence for a relationship between chronic intestinal inflammation and depression/anxiety (39, 40). The inflammation-depression relationship, therefore, is a likely mechanism linking decreases in negative behaviors and the abundance of these anti-inflammatory *Bacteroides* species.

Of the taxa that significantly associated with significant socioeconomic risk or behavioral dysregulation, three taxa belong to species containing known butyrate producers. The production of butyrate from plant-derived polysaccharides by the gut microbiome is understood to be an important mechanism through which high-fiber diets promote beneficial health effects. There are, however, only certain taxa that have the ability to produce butyrate (41). Surprisingly, two of the butyrate-producing species in our samples, *Coprococcus comes* and *Eubacterium rectale,* positively associated with elevated anxious-depression and reduced inhibitory control, respectively. This observation defies our expectation given that prior work points to butyrate’s important role in maintaining gut health and behavior dysregulation (42). It is possible that these taxa carry other functions that overwhelm the effects of their butyrate production on symptoms of behavioral dysregulation, or that overall butyrate production is reduced even though the relative abundances of these two taxa are high in certain microbiomes, or that butyrate production links to adverse behaviors under some contexts. On the other hand, the third butyrate-producing taxon in our samples, *Roseburia inulinivorans,* associated with a decrease in depressive problems and internalizing behavior. This is consistent with prior literature that suggests that increases in butyrate production improve overall mental health (42). Future work should seek to disentangle butryate’s specific role in mediating behavioral dysregulation and how its production by different taxa or in conjunction with different diets impacts this role.

In addition to the significant associations between the selected covariates and specific taxa, our analyses also linked these covariates to specific microbiome functions at the module and KO levels (Fig. 4; Supplemental Table 7). In particular, we found significant associations between a number of covariates and pathways involved in the bacterial Type VI secretion system (T6SS). Research into the psychology of depression has unveiled a possible link between depression/anxiety and chronic low-grade inflammation in the gut, suggesting a role for the microbiome in contributing to such disorders (39, 40). One possible mechanism for generating inflammation is dysbiosis caused by invading pathogens. For example, both *Vibrio* and *Salmonella* species can use the T6SS to attack commensal bacterial species and establish in the vertebrate gut (43, 44). Furthermore, T6SS have been shown to directly generate intestinal inflammation in a mouse model (45). We found that the abundances of three KOs assigned to a T6SS module, as well as a few KOs with possible T6SS homology, significantly associated with the increase in scores for aggression, anxiety, anxious-depression, depression, internalizing behavior, and number of turmoil-related life events. To determine if T6SS-associated KOs correlated with the abundances of any known T6SS-carrying taxa, we ran a similar CPGLM regression analysis as before (Supplemental Figure 1). We compared the abundances of all taxa and the five T6SS-assigned KOs. We found a number of taxa belonging to genera that we would expect, from prior investigations of their genomes, to carry T6SS such as *Bacteroides, Parabacteroides, and Escherichia* (46, 47). However, we also found significant associations with taxa assigned to genera with no documented cases, to our knowledge, of T6SS production, such as *Collinsella* and *Alistipes*. Future studies will be needed to elucidate whether type VI secretion systems have direct or indirect effects on the gut-brain axis and which taxa in the gut carry these systems.

In addition to T6SS, other mechanisms of inter-microbial competition also significantly associated with behavioral dysregulation. These associations included KOs assigned to functional groups involved in the synthesis of putative lipopeptide antibiotics (e.g. fengycin–an anti-fungal–and arthrofactin). These antibiotic-assigned KOs associated with the same set of behavioral dysregulation as the T6SS KOs, with the addition of emotional reactivity, decreased inhibitory control, and externalizing rather than internalizing behavior. Synthesis of lipopeptide antibiotics also significantly associated with adversity, such as family separation and poverty. The increase in these putative functions may indicate an increase in inter-microbial antagonism. This could possibly be due to invasion by pathogens, which could lead to intestinal inflammation that underlies their relationship with behaviors such as depression and anxiety.

Intriguingly, we also found relationships between both socioeconomic risk and behavioral dysregulation and microbial functional groups that have been implicated in modifying behaviors or cognitive function. For example, we discovered associations between these covariates and various KOs and modules associated with metabolism of monoamines that are often used as, or are common precursors to, neurotransmitters and neurohormones. We also found 8 covariates (7 behavioral, 1 socioeconomic risk) positively associated with modules involved in biosynthesis of melatonin from metabolism of tryptophan. Tryptophan is an essential amino acid, meaning it must be derived from the diet, and therefore the concentrations of available tryptophan can feasibly be altered by microbial metabolism (48). Indeed, many studies have found a relationship between symptoms of depression and anxiety and the availability of peripheral tryptophan (39, 49, 50). As a specific example, it has been shown that germ-free mice have greater plasma concentrations of tryptophan (49, 51), greater concentrations of hippocampal serotonin levels, and a lower kynurenine to tryptophan ratio (a common marker of tryptophan degradation; (49). Furthermore, germ-free mice were shown to have reduced levels of anxiety, as compared to conventional mice. Their anxiety, along with their kynurenine to tryptophan ratio, normalized after colonization with a conventional microbiome, presumably due to the introduction of taxa capable of metabolizing tryptophan and making it unavailable to the host (49). We found that the abundances of two KOs associated with degradation of tryptophan correlate with increases in behaviors including aggression, anxiety, anxious-depression, and impulsivity, as well as increases in exposure to adverse life events involving family separation, illness, and poverty. Moreover, we observed a KO involved in tryptophan biosynthesis that correlated with a decrease in life events related to family illness or injury.

Additionally, we detected significant associations between covariates and the metabolism of other notable monoamines such as glutamate (52–54), leucine (53, 55), and glutamine (56). Glutamate is the most abundant excitatory neurotransmitter in the vertebrate central nervous system as well as the most abundant amino acid in their diets (57). While dietary glutamate has not been linked to any neuropathology, the excitatory effects of glutamate have been linked to neurodegenerative disorders such as motor neuron disease (MND) or amyotrophic lateral sclerosis (ALS), Huntington’s disease, Parkinson’s disease and Alzheimer’s disease (57). Another monoamine, leucine can relatively easily pass through the blood brain barrier, where astrocytes convert it into glutamate (58, 59). Glutamine is also a precursor to glutamate, but is also directly involved in the maintenance of a healthy gut and its response to injury (60). Therefore, it is possible that the effect of the microbiome on the abundance of these monoamines may play a role in influencing the gut-brain axis.

Notably, these findings provide the foundation for future studies to replicate with larger samples and to assess longitudinal changes to better tease apart causal relationships. Additionally, this study offers a fundamental step toward translating animal models to sensitive periods of human development, and it provides a proof of concept to determine if the microbiome is linked to behavioral dysregulation and socioeconomic risk. Importantly, diet could be an important factor that confounds the relationships between the gut microbiome and socioeconomic risk or parent behavior. Properly interrogating the role of diet would require meticulously monitored diets, which was beyond the scope of the current study. Future work should build upon these findings to specifically interrogate the impacts of diet. If diet proved to be a mechanism driving these relationships, it could provide a targeted direction to include within psychosocial intervention designs.

## CONCLUSION

We tested associations between socioeconomic risk, child behavioral dysregulation, and the microbiome in terms of both taxonomic and functional composition in a cross-sectional sample 4-7 years old. In doing so, we discovered that not only are there significant associations between metrics of socioeconomic risk and behavioral dysregulation with the microbiome, but that the quality of caregiver behavior statistically moderated these relationships. Furthermore, we uncovered associations between individual taxa (e.g., *B. fragilis*) and functional groups (e.g. monoamine metabolism) within the microbiome and metrics of socioeconomic risk and behavioral dysregulation. These taxa and functional groups potentially, if replicated, represent mechanisms through which the microbiome associates with socioeconomic risk and behavioral dysregulation and possibly even targets for future intervention studies to investigate to improve children’s mental health outcomes.

The results of this study suggest that, when examining the trajectory of child psychological development, we need to consider biology, physiology, psychosocial environment, and the microbiome. All of these can have mutual effects, indicating that the way in which one factor impacts the psychological development of a child may change depending on the nature of one or more of the other relationships. Future studies, utilizing both human and animal models, should seek to tease apart specific behavioral links with the microbiome and extend this design to a wider range of behavioral symptomatology and socioeconomic risk.

## MATERIALS AND METHODS

### Sample Collection

Parents were instructed to collect a small stool sample from their child using a clean plastic collection device and OMNIgene-Gut collection tube (DNA Genotek, Ottawa, ON, Canada). Collection tubes were packaged and mailed at ambient temperature to the University of Oregon (Eugene, OR), where they transferred to −80°C upon receipt. See Supplemental Methods for greater detail, including measures of diet and health.

### Questionnaires

*Socioeconomic Risk* were indexed using metrics of socioeconomic status and the Life Events Checklist (LEC; (61)). The Life Events Checklist was used to provide an index adverse home environment exposure. This provides a total score, and subscales to identify specific components of adverse life events. Subscales included poverty, turmoil, family illness, neighborhood violence, family separation, and an overall total score. Household poverty was indexed by incomes-need-ratio. See Supplemental Methods for range, mean and SD of subscales.

*Behavioral dysregulation* were indexed using two previously validated parent-report measures: the Child Behavior Questionnaire (CBQ; (62)) and the Child Behavior Checklist (CBCL; (63)). Given childhood is a period in which behavioral dysregulation symptoms shares common risk factors and less differentiation across both internalizing and externalizing dimensions of disorders than typically discussed in adult samples, we included both internalizing (e.g., depression, anxiety) and externalizing (e.g., inhibitory control, aggression) symptoms in our analyses. Subscales of interested included anxiety problems, depression, emotional reactivity, anxious depressed, internalizing total, aggressive behavior, externalizing total, overall total score, and inhibitory control. See Supplemental Methods for range, mean and SD of subscales.

*Caregiver Behavior* was indexed via parent-report Parenting Stress Index (PSI; (64)) Interpersonal Mindfulness in Parenting (IEM-P; (65)), and the Five Factor Mindfulness Questionnaire (FFMQ; (66)). These questionnaires provided a range of perceived parental stress and wellbeing, both in general and within the parent-child relationship. See Supplemental Methods for range, mean and SD of subscales.

### DNA extraction and sequencing

DNA was extracted from 250 µl aliquots of the OMNIgene-Gut samples using the MoBio PowerLyzer PowerSoil kit (Qiagen, Hilden, Germany) with the following protocol modifications: following the addition of solution C1, a 1-minute bead-beating step was performed on a Mini-BeadBeater-96 (BioSpec Products, Bartlesville, OK, USA), followed by a 10-minute incubation at 65°C; in the final step DNA was eluted in two stages for a combined total of 100 µl.

### Metagenomic analyses

Raw metagenome sequences were prepared for analysis using the shotcleaner workflow (67), which follows the Human Microbiome Project Consortium data processing guidelines (68). All raw sequences can be accessed through the NCBI at BioProject PRJNA496479, and the code for all analyses can be accessed at https://github.com/kstagaman/flannery_stagaman_analysis. Briefly, low quality sequences are trimmed or removed, sequences matching to the human genome are discarded, and identical sequences are collapsed into a single read. As additional quality control, we removed 3 of 40 fecal samples due to poor sequencing coverage (coverage range of removed samples: 19,013 to 23,743; coverage range of remaining samples: 3,499,106 to 15,776,004). These high-quality sequences were then run through shotmap (67) to quantify KEGG Orthology (KO) group relative abundance and metaphlan2 to quantify taxon relative abundance (69). All resulting data and the sample metadata (Supplemental Table 1) were analyzed in R (70).

We applied a data reduction technique to minimize the number of covariates considered in our subsequent analyses. This process is important to reduce the potential for model overfitting given the large number of covariates relative to the number of samples measured in our study. Using the *ordinate* function from the phyloseq package (71), we generated a PCoA ordination based on the Bray-Curtis dissimilarities for both the functional (KO) and taxonomic communities (Supplemental Figure 2). Briefly, we applied the *envfit* function (72) to Bray-Curtis dissimilarity-based PCoAs of microbiome taxonomy (species level) or functional capacity and identified covariates that explained a significant amount of variation across individuals (Supplemental Figures 3 & 4). Despite being analyzed independently, an identical set of 10 significant covariates best explained the taxonomic and functional variation among individuals. This finding is unsurprising given the strong correlation between taxonomic and functional beta-diversity (Procrustes *r* =∼ 0.84, *p* < 0.0001; Supplemental Table 8). The significant covariates used in our successive analyses are defined in Table 1. See Supplemental Methods for additional details.

We utilized a constrained correspondence analysis (CCA; *cca* function; (72)) to determine the variance in microbiome composition (functional and taxonomic) that covariates within the socioeconomic risk, child behavioral dysregulation, and caregiver behavior categories explained. The CCA method is useful in this case because it allows us to first account for the variance in microbiome composition explained by demographic and gut-related covariates, which might otherwise confound our analysis, before assessing the variance explained by the covariates of interest for this study. We assessed the significance of associations between the selected covariates and the microbiome using a permutational ANOVA (PERMANOVA) analysis (*anova.cca* function; (72)) on the resulting CCA ordination.

To determine if the *envfit*-selected caregiver behavior covariate Parent-Child Dysfunction interacted with either the socioeconomic risk or child behavioral dysregulation covariates, we first built a CCA models (one for socioeconomic risk, one for child behavioral dysregulation) with all possible covariate interactions. However, this builds large models that reduce our chance of finding real, significant association due to the number of terms. Therefore, before running a PERMANOVA test, we subjected each CCA object to model selection based on the Akaike Information Criterion (AIC) by stepwise addition or subtraction of terms (*ordistep* function; (72)). The model selected by this method was then analyzed using PERMANOVA to determine if there were significant associations between covariate interactions and the microbiome. All of these computational methods are available as supplemental data.

The above methods analyze the relationships between the covariates of interest and the overall composition of the microbiome (in terms of taxonomy and functional potential), but they may miss important relationships between covariates and individual taxa or microbial functions. To determine if such relationships exist in this data set we conducted pairwise regressions between the abundance of each taxa or KO and each socioeconomic risk and child behavioral dysregulation covariate. We included in each regression model the same demographic and gut-related terms to account for their variance as well. The regression method used was a compound Poisson generalized linear model (CPGLM; (73)), which uses a distribution that has a point mass over zero, allowing it to better handle the sparseness of functional and taxonomic community data (74). After all pairwise regressions, we adjusted the p-values using the False Discovery Rate (FDR) with a cutoff of *q* = 0.05. We then removed any pairs where the taxon or KO was absent from half of the samples or more and presented the results in Supplemental Tables 6 and 7.

## ACKNOWLEDGEMENTS

This work was supported by a Hemera Foundation grant awarded to P.A.F., a National Science Foundation Graduate Research Fellowship [2015172132] to J.E.F.; its contents are solely the responsibility of the authors and do not necessarily represent the official views of the NIH. Hickey was supported by a postdoctoral fellowship funded by a grant to the BioBE Center from the Alfred P. Sloan Foundation. Stagaman was supported by the NIEHS Integrated Regional Training Program in Environmental Health Sciences grant (PI Robert L Tanguay, T32-ES007060-38).

We would like to thank Jessica Green and Clarisse Betancourt for project assistance, including DNA extraction and subject running. We would also like to thank Natalia Pachote and Adrienne Luba for assistance in project coordination and subject running.

## SUPPLEMENTAL METHODS

### Sample Collection

A subsample of families from a larger study conducted in the Stress Neurobiology and Prevention laboratory were asked to participate in a follow-up study to collect a child gut microbial sample via at home stool collection. Parents were instructed to wait to collect sample at least 2-4 weeks following antibiotic use or illness; no current stool irregularities, no anticipated stressors, and during a week with a typical diet. Recruitment was pre-determined to be complete once we reached 40 completed samples. Forty-five families consented to be in the study; five families did not complete the stool sample; one sample was determined to be lost in the mail, one child remained within the window of recent antibiotic use and illness through the duration of the study, one child changed their mind about participating, and two families continued to express interest in completing the sample but did not return a sample. Two experimenters went to the family’s home. Parents provided consented and children provided assent. A visual depiction of the study (coloring book) was used to ensure child understood the study. During the home visit, parents filled out questionnaires and parents were instructed to collect a stool sample from their child a week after the visit using Genotek OmiGene kits (DNA Genotek, Ottawa, ON, Canada). This procedure allowed families to mail the sample in after collection without sample degradation. This was important to reach a broad range of socioeconomic backgrounds and to eliminate variability in post-collection procedures across the sample. The experimenter provided a collection demonstration with a toilet seat and playdough for parent to collect the sample from their child a week after the visit. In the week prior to collection, parents were asked to fill out a daily diary of basic food categories the child ate at breakfast, lunch, and dinner. Notably, parent’s knowledge of child’s daily diet was variable depending on child’s enrollment in subsidized lunch at school and mother’s work schedule. Families were compensated for their time at the home visit and again after receiving the stool sample.

### Important Runtime Parameters

shotcleaner.pl

> Output format [-of]: fastq
>
> Bowtie database name [-n]: all_GRCh38.p7

shotmap.pl

> Shotmap database [-d]: KEGG_021515_1M
>
> Class score [--class-score]: 34
>
> [--ags-method]: none

### Analysis in R

#### Data processing

Functional and taxonomic community tables were built using relative abundances and associated with the sample metadata using the package phyloseq. All participants’ (mothers and children) ages were calculated *in days* from their date of birth to the date of the second session, when stool samples were collected.

We had both the forward and reverse reads for each sequence. We conducted a Procrustes analysis on PCoA ordinations based on the Bray-Curtis dissimilarities to determine if there was a significant correlation between the two read sets for both the functional and taxonomic reads. For both functional and taxonomic reads, the correlation coefficients between the forward and reverse read based ordinations was greater than 0.99 and statistically significant (*p* = 0.0001). We therefore continued with the remainder of the analyses using only the forward reads.

#### Covariate reduction

Within each covariate category (ESA, child behavior, parenting, demography, and gut-related history), we used the *envfit* function from the vegan package to determine which covariates (e.g., ESA covariates include LEC Poverty Related Events and LEC Turmoil; child behavior covariates include CBQ Impusivity and CBQ Inhibitory Control) explained a significant proportion of microbiome diversity along any of the first four PCoA axes (same PCoA ordination generated in the Data Processing section above; Supplemental Figures 3 & 4).

The code used to conduct all analyses can be found at https://github.com/kstagaman/flannery_stagaman_analysis.

All metagenome data can be found at https://www.ncbi.nlm.nih.gov/sra/?term=PRJNA496479 https://www.ncbi.nlm.nih.gov/Traces/study/?acc=PRJNA496479

**Supplemental Figure 1.**
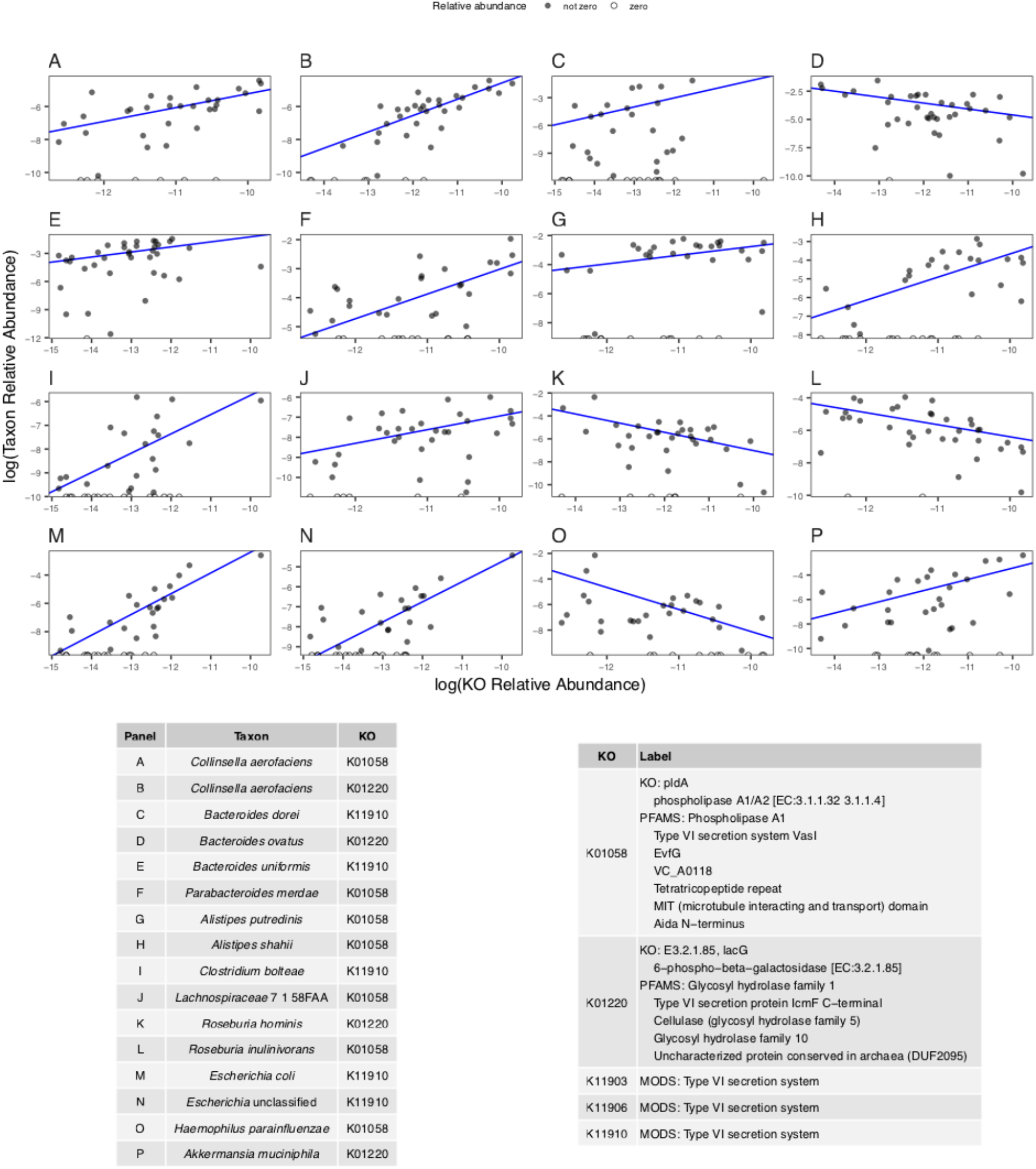
Each panel (A-P) is a scatter plot of the relative abundance of a single KO (x-axis, log-transformed) and the relative abundance of an individual taxon (y-axis, log transformed). The blue lines represent the CPGLM regression line as fit to the data. Filled circles represent taxon abundances that were greater than zero before log transformation, and open circles represent taxon abundances that were zero before log transformation. The table on the left details the KO-taxon pair used in each panel, and the table on the right gives the descriptive name of each KO identification number.

**Supplemental Figure 2.**
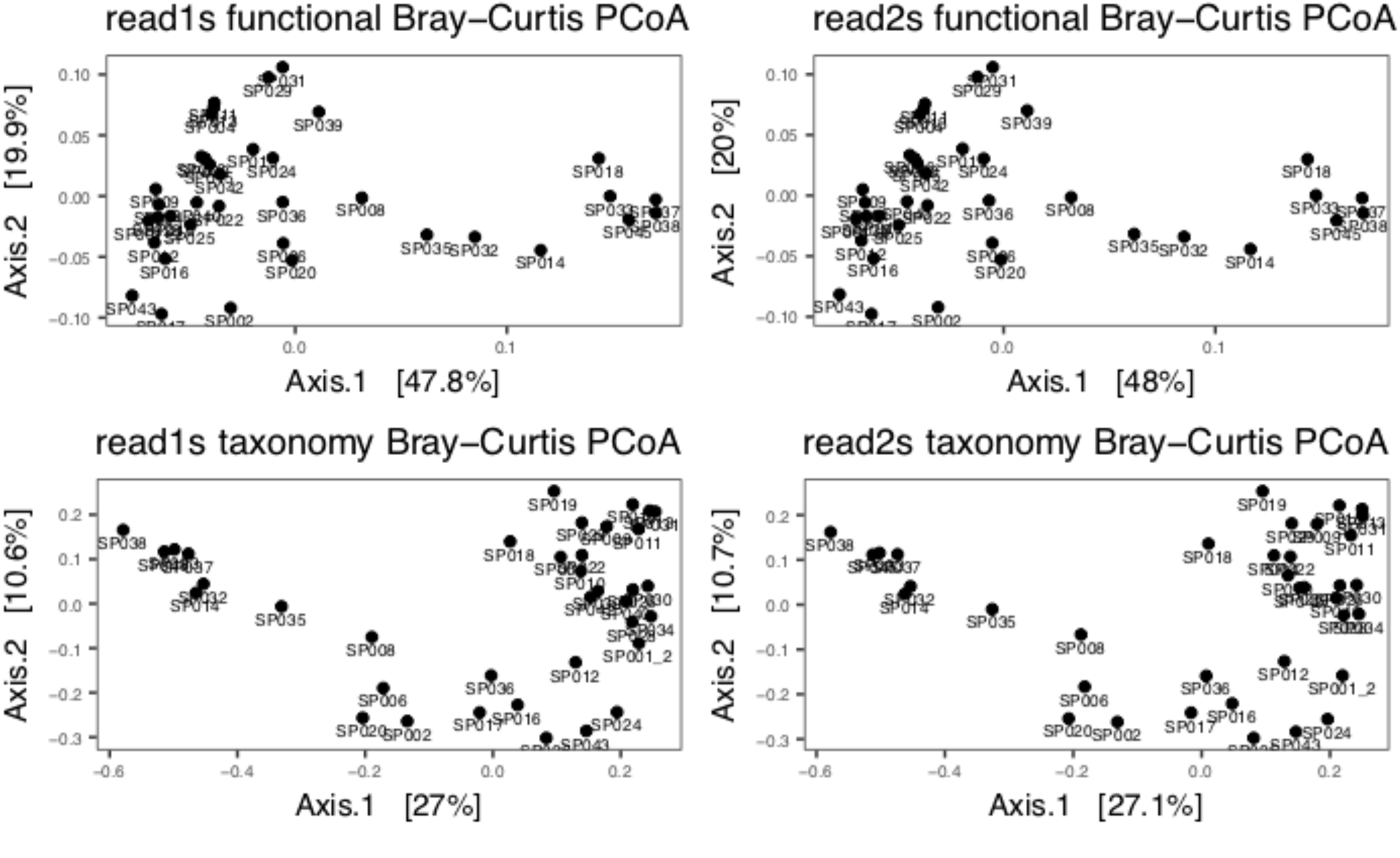
Principal Coordinate Analysis ordinations for the metagenomic data. The top two panels were created using the KO-annotated sequences for the read1 (left) and read2 (right) data. The bottom two panels were created using the taxon-annotated sequences for the read1 (left) and read2 (right) data. The percentages in brackets along each axis represent the total variance explained by that axis. All distances were measured using the Bray-Curtis dissimilarity.

**Supplemental Figure 3.**
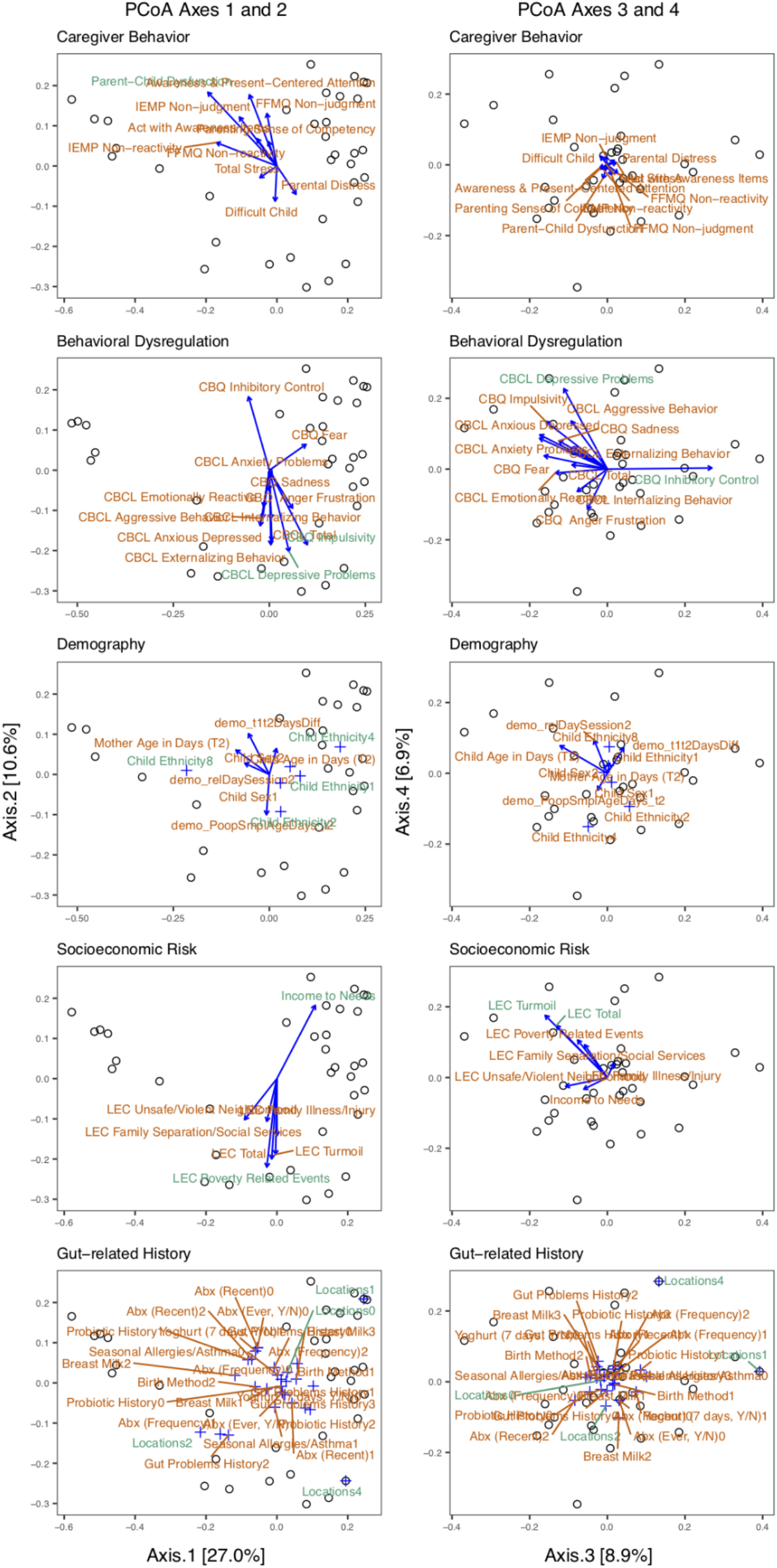
The results of the envfit analysis for each category of covariates (each row of panels corresponds to an analysis within a single category) on the taxonomy-based PCoA ordinations. The panels on the left show the first and second axes of each ordination and the panels on the right show the third and fourth axes of each ordination.

**Supplemental Figure 4.**
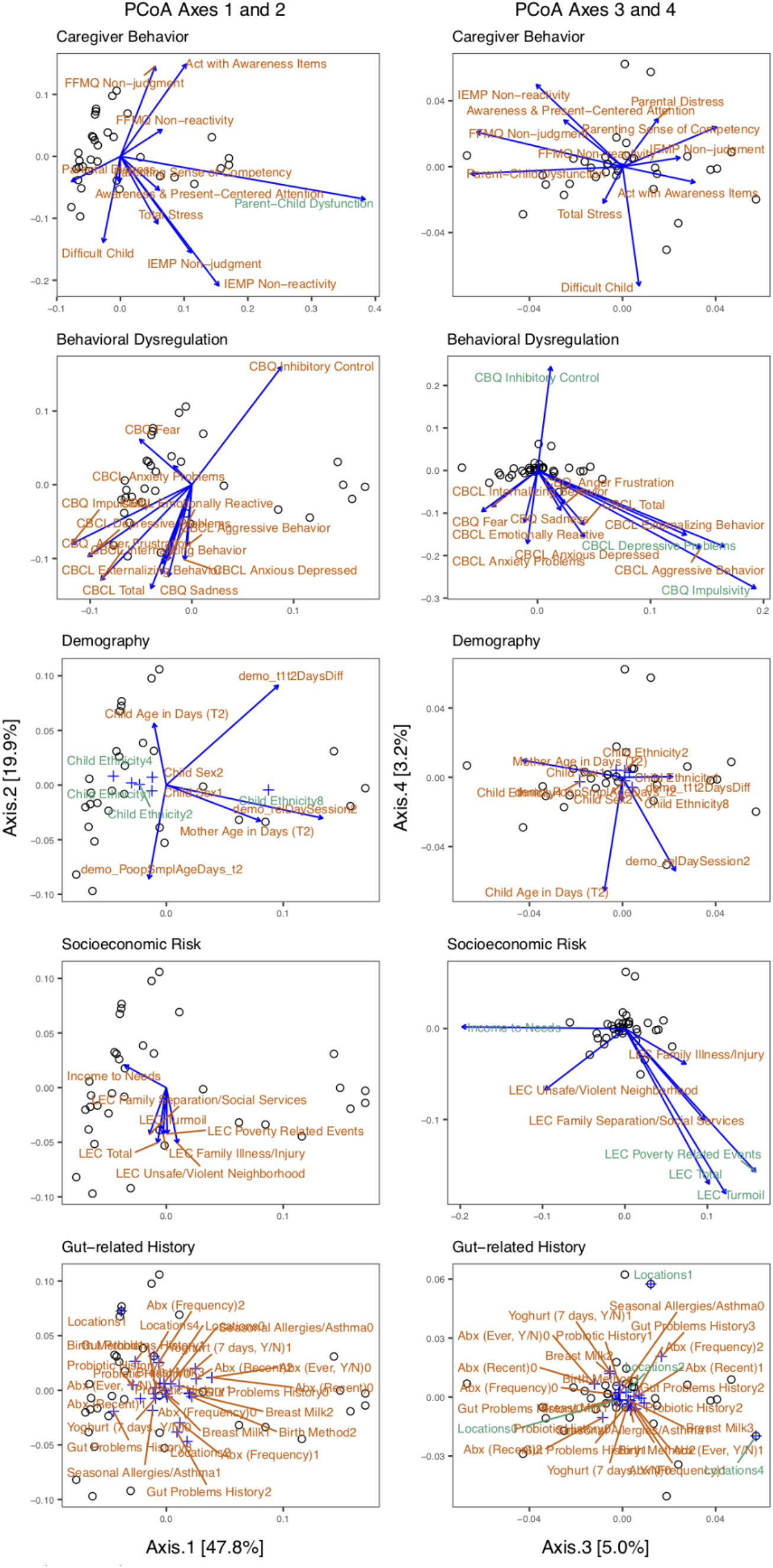
The results of the *envfit* analysis for each category of covariates (each row of panels corresponds to an analysis within a single category) on the functional group-based PCoA ordinations. The panels on the left show the first and second axes of each ordination and the panels on the right show the third and fourth axes of each ordination.

Supplemental Table 1 is a dataset provided as a separate file.

**Supplemental Table 2.** Results of PERMANOVA analysis on AIC-selected covariates within Socioeconomic Risk and Caregiver Behavior (including interactions) and their relationship with the taxonomic-based composition of the microbiome.

|  | Df | ChiSquare | F | Pr(>F) |  |
| --- | --- | --- | --- | --- | --- |
| LEC Turmoil | 1 | 0.216 | 1.611 | 0.009 | * |
| LEC Poverty Related Events | 1 | 0.136 | 1.01 | 0.458 |  |
| Parent-Child Dysfunction | 1 | 0.131 | 0.973 | 0.495 |  |
| LEC Turmoil:Parent-Child Dysfunction | 1 | 0.177 | 1.319 | 0.126 |  |
| LEC Poverty Related Events :Parent-Child Dysfunction | 1 | 0.186 | 1.386 | 0.073 |  |
| Residual | 21 | 2.822 |  |  |  |

**Supplemental Table 3.** Results of PERMANOVA analysis on AIC-selected covariates within Socioeconomic Risk and Caregiver Behavior (including interactions) and their relationship with the functional group-based composition of the microbiome.

|  | Df | ChiSquare | F | Pr(>F) |  |
| --- | --- | --- | --- | --- | --- |
| LEC Poverty Related Events | 1 | 0.003 | 1.222 | 0.226 |  |
| LEC Turmoil | 1 | 0.004 | 1.433 | 0.139 |  |
| Income to Needs | 1 | 0.004 | 1.483 | 0.113 |  |
| Parent-Child Dysfunction | 1 | 0.005 | 1.763 | 0.069 |  |
| LEC Poverty Related Events :Parent-Child Dysfunction | 1 | 0.004 | 1.574 | 0.108 |  |
| LEC Turmoil:Parent-Child Dysfunction | 1 | 0.007 | 2.506 | 0.01 | * |
| Income to Needs:Parent-Child Dysfunction | 1 | 0.005 | 1.859 | 0.041 | * |
| Residual | 19 | 0.054 |  |  |  |

**Supplemental Table 4.** Results of PERMANOVA analysis on AIC-selected covariates within Behavioral dysregulation and Caregiver Behavior (including interactions) and their relationship with the taxonomic-based composition of the microbiome.

|  | Df | ChiSquare | F | Pr(>F) |
| --- | --- | --- | --- | --- |
| CBCL Depressive Problems | 1 | 0.206 | 1.485 | 0.071 |
| Parent-Child Dysfunction | 1 | 0.135 | 0.973 | 0.489 |
| Residual | 24 | 3.328 |  |  |

**Supplemental Table 5.**
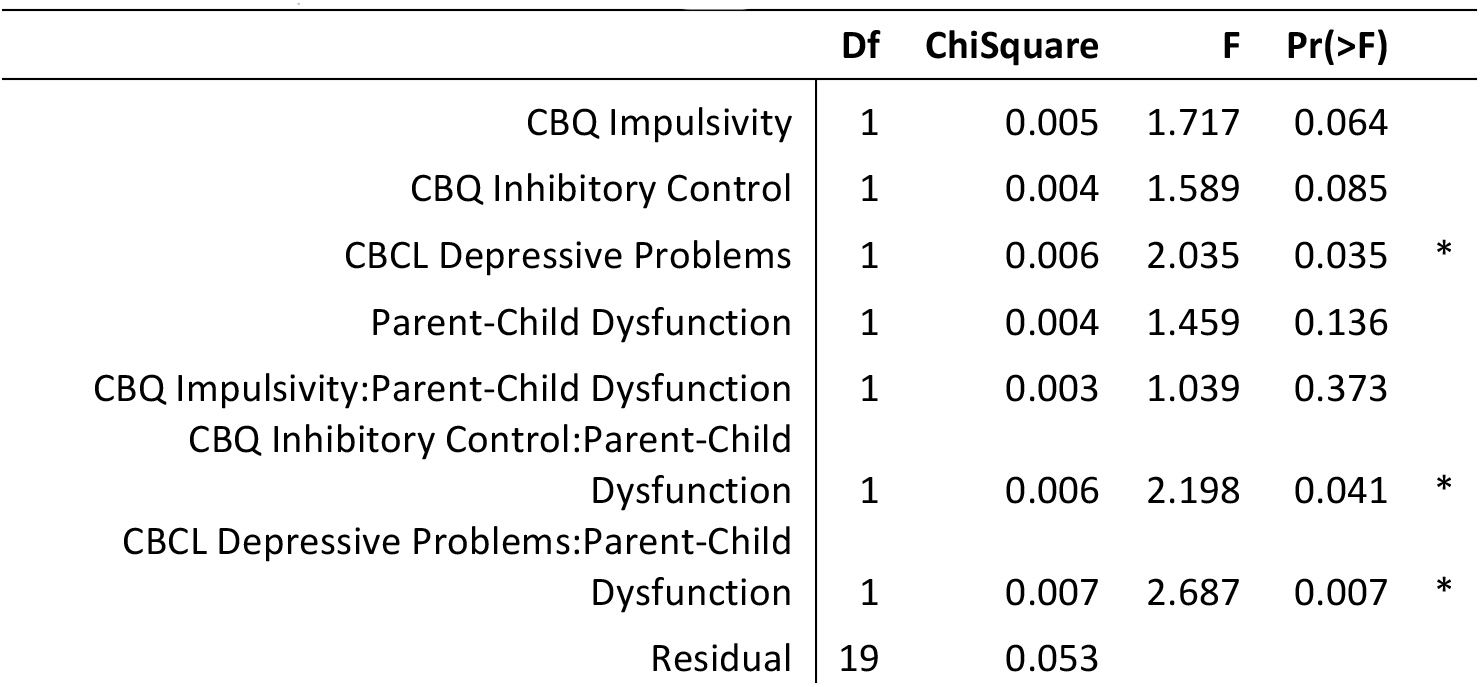
Results of PERMANOVA analysis on AIC-selected covariates within Behavioral dysregulation and Caregiver Behavior (including interactions) and their relationship with the functional group-based composition of the microbiome.

|  | Df | ChiSquare | F | Pr(>F) |  |
| --- | --- | --- | --- | --- | --- |
| CBQ Impulsivity | 1 | 0.005 | 1.717 | 0.064 |  |
| CBQ Inhibitory Control | 1 | 0.004 | 1.589 | 0.085 |  |
| CBCL Depressive Problems | 1 | 0.006 | 2.035 | 0.035 | * |
| Parent-Child Dysfunction | 1 | 0.004 | 1.459 | 0.136 |  |
| CBQ Impulsivity:Parent-Child Dysfunction | 1 | 0.003 | 1.039 | 0.373 |  |
| CBQ Inhibitory Control:Parent-Child Dysfunction | 1 | 0.006 | 2.198 | 0.041 | * |
| CBCL Depressive Problems:Parent-Child Dysfunction | 1 | 0.007 | 2.687 | 0.007 | * |
| Residual | 19 | 0.053 |  |  |  |

Supplemental Table 6 is a dataset provided as a separate file.

Supplemental Table 7 is a dataset provided as a separate file.

**Supplemental Table 8.** R-squared values and p-values from Procrustes analyses comparing the ordinations based on taxonomic or functional group composition, and between read1 and read2 sequencing data.

| Read comp. | Type comp. | R<sup>2</sup> | p-value |
| --- | --- | --- | --- |
| 1 vs. 2 | KOs vs. KOs | 0.9997 | 1.00E-04 |
|  | Taxa vs. Taxa | 0.9973 | 1.00E-04 |
| 1 vs. 1 | KOs vs. Taxa | 0.8375 | 1.00E-04 |
| 2 vs. 2 |  | 0.8364 | 1.00E-04 |

