## Supplementary material for "Gut feelings begin in childhood: how the gut metagenome links to early environment, caregiving, and behavior": Supp Table 6

| Variable | Relationship | Kingdom | Phylum | Class | Order | Family | Genus | Species | sTrain | | References |
|---|---|---|---|---|---|---|---|---|---|---|---|
| CBCL Aggressive Behavior (T1)<br>CBCL Anxious Depressed (T1)<br>CBCL Emotionally Reactive (T1)<br>CBCL Externalizing Behavior (T1)<br>CBCL Total (T1)<br>CBQ Impulsivity (T1)<br>CBQ Inhibitory Control (T1)<br>LEC Total<br>LEC Turmoil | -<br>-<br>-<br>-<br>-<br>-<br>+<br>-<br>- | Bacteria | Bacteroidetes | Bacteroidia | Bacteroidales | Bacteroidaceae | Bacteroides | Bacteroides_fragilis | Bacteroides_fragilis_unclassified | Plot | 1. J. Ochoa-Repáraz et al., A polysaccharide from the human commensal Bacteroides fragilis protects against CNS demyelinating disease. Mucosal Immunol. 3, 487–495 (2010).<br>2. E. Y. Hsiao et al., Microbiota Modulate Behavioral and Physiological Abnormalities Associated with Neurodevelopmental Disorders. Cell. 155, 1451–1463 (2013).<br>3. [review] T. R. Sampson, S. K. Mazmanian, Control of brain development, function, and behavior by the microbiome. Cell Host Microbe. 17, 565–576 (2015).<br>4. [review] J. F. Cryan, T. G. Dinan, More than a Gut Feeling: the Microbiota Regulates Neurodevelopment and Behavior. Neuropsychopharmacology. 40, 241–242 (2015). |
| CBCL Aggressive Behavior (T1)<br>CBCL Anxious Depressed (T1)<br>CBCL Depressive Problems (T1)<br>CBCL Externalizing Behavior (T1)<br>CBQ Inhibitory Control (T1)<br>LEC Total<br>LEC Turmoil | +<br>+<br>+<br>+<br>-<br>+<br>+ | Bacteria | Actinobacteria | Actinobacteria | Bifidobacteriales | Bifidobacteriaceae | Bifidobacterium | Bifidobacterium_adolescentis | Bifidobacterium_adolescentis_unclassified | Plot | 1. A. Clark, N. Mach, Exercise-induced stress behavior, gut-microbiota-brain axis and diet: a systematic review for athletes. J. Int. Soc. Sports Nutr. 13, 43 (2016).<br>2. D. Gevers et al., The Treatment-Naive Microbiome in New-Onset Crohn's Disease. Cell Host Microbe. 15, 382–392 (2014). |
| CBCL Anxious Depressed (T1)<br>CBCL Depressive Problems (T1)<br>CBCL Externalizing Behavior (T1)<br>CBCL Total (T1)<br>CBQ Anger Frustration (T1)<br>LEC Turmoil | +<br>+<br>+<br>+<br>+<br>+ | Bacteria | Actinobacteria | Actinobacteria | Coriobacteriales | Coriobacteriaceae | Collinsella | Collinsella_aerofaciens | GCF_000169035 | Plot | 1. M. De Angelis et al., Fecal Microbiota and Metabolome of Children with Autism and Pervasive Developmental Disorder Not Otherwise Specified. PLoS One. 8, e76993 (2013).<br>2. A. L. Richards et al., Gut microbiota composition impacts host gene expression by changing chromatin accessibility, 1–18 (2018).<br>3. L. Bonfrate, J. Tack, Microbiota in health and irritable bowel syndrome: current knowledge, perspectives and therapeutic options. Scand. J. Gastroenterol. 48, 995–1009 (2013).<br>4. J. Jalanka-Tuovinen et al., Faecal microbiota composition and host–microbe cross-talk following gastroenteritis and in postinfectious irritable bowel syndrome. Gut. 63, 1737–1745 (2014). |
| CBCL Aggressive Behavior (T1)<br>CBCL Anxious Depressed (T1)<br>CBCL Externalizing Behavior (T1)<br>CBCL Total (T1)<br>CBQ Fear (T1)<br>Income to Needs | -<br>-<br>-<br>-<br>-<br>+ | Bacteria | Proteobacteria | Gammaproteobacteria | Pasteurellales | Pasteurellaceae | Haemophilus | Haemophilus_parainfluenzae | Haemophilus_parainfluenzae_unclassified | Plot | 1. D. Gevers et al., The treatment-naive microbiome in new-onset Crohn's disease. Cell Host Microbe. 15, 382–392 (2014).<br>2. Y. Qiao et al., Alterations of oral microbiota distinguish children with autism spectrum disorders from healthy controls. Sci. Rep. 8, 1–12 (2018).<br>3. P. Wurm et al., Antibiotic-associated apoptotic enterocolitis in the absence of a defined pathogen: The role of intestinal microbiota depletion. Crit. Care Med. 45, e600–e606 (2017). |
| CBCL Anxiety Problems (T1)<br>CBCL Emotionally Reactive (T1)<br>CBQ Inhibitory Control (T1)<br>Income to Needs<br>LEC Total | -<br>-<br>+<br>+<br>- | Bacteria | Bacteroidetes | Bacteroidia | Bacteroidales | Bacteroidaceae | Bacteroides | Bacteroides_dorei | Bacteroides_dorei_unclassified | Plot | 1. A. Endo, A. Prtty, M. Kalliomki, E. Isolauri, S. Salminen, Long-term monitoring of the human intestinal microbiota from the 2nd week to 13 years of age. Anaerobe. 28, 149–156 (2014).<br>2. E. B. Hollister et al., Structure and function of the healthy pre-adolescent pediatric gut microbiome. Microbiome. 3, 36 (2015).<br>3. T. Vatanen et al., Variation in Microbiome LPS Immunogenicity Contributes to Autoimmunity in Humans. Cell. 165, 842–853 (2016). |
| CBCL Depressive Problems (T1)<br>CBCL Emotionally Reactive (T1)<br>CBCL Internalizing Behavior (T1)<br>CBQ Anger Frustration (T1)<br>CBQ Sadness (T1) | +<br>+<br>+<br>+<br>+ | Bacteria | Firmicutes | Clostridia | Clostridiales | Clostridiaceae | Clostridium | Clostridium_bolteae | Clostridium_bolteae_unclassified | Plot | **1. [review] M. C. Toh, E. Allen-Vercoe, The human gut microbiota with reference to autism spectrum disorder: considering the whole as more than a sum of its parts. Microb. Ecol. Heal. Dis. 26 (2015), doi:10.3402/mehd.v26.26309.**<br>**2. [review] H. T. Ding, Y. Taur, J. T. Walkup, Gut Microbiota and Autism: Key Concepts and Findings. J. Autism Dev. Disord. 47, 480–489 (2017).**<br>**3. [review] J. M. Kinross, A. W. Darzi, J. K. Nicholson, Gut microbiome-host interactions in health and disease. Genome Med. 3, 14 (2011).**<br>**4. [review] P. Louis, Does the Human Gut Microbiota Contribute to the Etiology of Autism Spectrum Disorders? Dig. Dis. Sci. 57, 1987–1989 (2012).** |
| CBQ Impulsivity (T1)<br>LEC Family Illness/Injury<br>LEC Poverty Related Events<br>LEC Total | +<br>+<br>+<br>+ | Bacteria | Bacteroidetes | Bacteroidia | Bacteroidales | Rikenellaceae | Alistipes | Alistipes_onderdonkii | GCF_000374505 | Plot | 1. A. Evrensel, M. E. Ceylan, Gut-Microbiota-Brain Axis and Depression. Understanding Depression (Springer Singapore, Singapore, 2018; http://linkinghub.elsevier.com/retrieve/pii/B9781856178099100179), pp. 197–207.<br>2. J. V. Vlainić, J. Šuran, T. Vlainić, A. L. Vukorep, Probiotics as an Adjuvant Therapy in Major Depressive Disorder. Curr. Neuropharmacol. 14, 952–958 (2016).<br>3. Y. Kang, Y. Cai, Gut microbiota and depression. Rev. Med. Microbiol. 28, 56–62 (2017).<br>4. [review] T. G. Dinan, J. F. Cryan, Microbes, Immunity and Behavior: Psychoneuroimmunology Meets the Microbiome. Neuropsychopharmacology. 42, 178–192 (2017). |
| CBCL Aggressive Behavior (T1)<br>CBQ Impulsivity (T1)<br>CBQ Inhibitory Control (T1) | +<br>+<br>- | Bacteria | Firmicutes | Clostridia | Clostridiales | Eubacteriaceae | Eubacterium | Eubacterium_siraeum | Eubacterium_siraeum_unclassified | Plot | 1. H. Mkaouar et al., Siropins, novel serine protease inhibitors from gut microbiota acting on human proteases involved in inflammatory bowel diseases. Microb. Cell Fact. 15, 1–13 (2016).<br>2. M. De Angelis et al., Fecal Microbiota and Metabolome of Children with Autism and Pervasive Developmental Disorder Not Otherwise Specified. PLoS One. 8, e76993 (2013). |
| CBCL Aggressive Behavior (T1)<br>CBCL Anxious Depressed (T1)<br>Income to Needs | -<br>-<br>+ | Bacteria | Firmicutes | Negativicutes | Selenomonadales | Veillonellaceae | Veillonella | Veillonella_unclassified | t_NA | Plot | |
| CBQ Anger Frustration (T1)<br>CBQ Sadness (T1) | +<br>+ | Bacteria | Firmicutes | Clostridia | Clostridiales | Ruminococcaceae | Anaerotruncus | Anaerotruncus_colihominis | GCF_000154565 | Plot | |
| CBCL Externalizing Behavior (T1)<br>CBCL Total (T1) | -<br>- | Bacteria | Bacteroidetes | Bacteroidia | Bacteroidales | Bacteroidaceae | Bacteroides | Bacteroides_thetaiotaomicron | Bacteroides_thetaiotaomicron_unclassified | Plot | |
| CBCL Anxiety Problems (T1)<br>CBQ Fear (T1) | +<br>+ | Bacteria | Proteobacteria | Deltaproteobacteria | Desulfovibrionales | Desulfovibrionaceae | Bilophila | Bilophila_unclassified | t_NA | Plot | |
| CBCL Depressive Problems (T1)<br>CBCL Internalizing Behavior (T1) | -<br>- | Bacteria | Firmicutes | Clostridia | Clostridiales | Lachnospiraceae | Roseburia | Roseburia_inulinivorans | GCF_000174195 | Plot | |
| CBCL Aggressive Behavior (T1)<br>Income to Needs | -<br>+ | Bacteria | Firmicutes | Bacilli | Lactobacillales | Streptococcaceae | Streptococcus | Streptococcus_salivarius | Streptococcus_salivarius_unclassified | Plot | |
| LEC Total<br>LEC Turmoil | +<br>+ | Bacteria | Firmicutes | Clostridia | Clostridiales | Ruminococcaceae | Subdoligranulum | Subdoligranulum_unclassified | t_NA | Plot | |
| CBCL Depressive Problems (T1) | + | Bacteria | Verrucomicrobia | Verrucomicrobiae | Verrucomicrobiales | Verrucomicrobiaceae | Akkermansia | Akkermansia_muciniphila | GCF_000020225 | Plot | 1. M. C. Toh, E. Allen-Vercoe, The human gut microbiota with reference to autism spectrum disorder: considering the whole as more than a sum of its parts. Microb. Ecol. Heal. Dis. 26 (2015), doi:10.3402/mehd.v26.26309. |
| Income to Needs | + | Bacteria | Firmicutes | Clostridia | Clostridiales | Lachnospiraceae | Anaerostipes | Anaerostipes_hadrus | GCF_000332875 | Plot | |
| CBCL Anxious Depressed (T1) | + | Bacteria | Firmicutes | Clostridia | Clostridiales | Lachnospiraceae | Coprococcus | Coprococcus_comes | GCF_000155875 | Plot | |
| CBQ Inhibitory Control (T1) | - | Bacteria | Firmicutes | Clostridia | Clostridiales | Eubacteriaceae | Eubacterium | Eubacterium_rectale | Eubacterium_rectale_unclassified | Plot | |
| LEC Family Illness/Injury | + | Bacteria | Bacteroidetes | Bacteroidia | Bacteroidales | Porphyromonadaceae | Parabacteroides | Parabacteroides_distasonis | Parabacteroides_distasonis_unclassified | Plot | |
| Income to Needs | - | Bacteria | Bacteroidetes | Bacteroidia | Bacteroidales | Prevotellaceae | Prevotella | Prevotella_copri | GCF_000157935 | Plot | 1. A. J. Obregon-Tito et al., Subsistence strategies in traditional societies distinguish gut microbiomes. Nat. Commun. 6, 6505 (2015).<br>2. L. Prideaux et al., Impact of Ethnicity, Geography, and Disease on the Microbiota in Health and Inflammatory Bowel Disease. Inflamm. Bowel Dis. 19, 2906–2918 (2013).<br>3. T. Yatsunenko et al., Human gut microbiome viewed across age and geography. Nature. 457, 222–227 (2012).<br>4. J. F. Ruiz-calderon et al., Walls talk: Microbial biogeography of homes spanning urbanization. Sci. Adv. 2, e1501061–e1501061 (2016).<br>5. J. C. Clemente, L. K. Ursell, L. W. Parfrey, R. Knight, The Impact of the Gut Microbiota on Human Health: An Integrative View. Cell. 148, 1258–1270 (2012).<br>6. S. L. Schnorr et al., Gut microbiome of the Hadza hunter-gatherers. Nat. Commun. 5, 3654 (2014).<br>7. C. De Filippo et al., Impact of diet in shaping gut microbiota revealed by a comparative study in children from Europe and rural Africa. Proc. Natl. Acad. Sci. 107, 14691–14696 (2010).<br>8. A. Gomez et al., Gut Microbiome of Coexisting BaAka Pygmies and Bantu Reflects Gradients of Traditional Subsistence Patterns. Cell Rep. 14, 2142–2153 (2016).<br>9. K. Stagaman et al., Market Integration Predicts Human Gut Microbiome Attributes across a Gradient of Economic Development. mSystems. 3, e00122-17 (2018). |
| Income to Needs | + | Bacteria | Firmicutes | Clostridia | Clostridiales | Lachnospiraceae | Roseburia | Roseburia_intestinalis | Roseburia_intestinalis_unclassified | Plot | 1. T. G. Dinan, J. F. Cryan, Microbes, Immunity and Behavior: Psychoneuroimmunology Meets the Microbiome. Neuropsychopharmacology. 42, 178–192 (2017). |

| Notes |
| --- |
| 4. Major depressives had increased levels of Enterobacteriaceae and Alistipes but reduced levels of Faecalibacterium. |
| conjugated linoleic acid (CLA) is the most extensively studied with multiple putative health benefits including an antiatherosclerotic effect, an antiobesogenic impact, and probable anticarcinogenic properties<br>Roseburia spp. has been identified among the most active producers in the gut (Devillard et al, 2007)<br>Unlike polyunsaturated fatty acids, so far no clear mental health benefits from CLA have been identified. |

Supplemental Table 6. Every significant association between an individual covariate and taxon (assign

| Variable | Relationship | Kingdom | Phylum | Class |
|---|---|---|---|---|
| CBCL Aggressive Behavior (T1) | - | Bacteria | Bacteroidetes | Bacteroidia |
| CBCL Anxious Depressed (T1) | - | Bacteria | Bacteroidetes | Bacteroidia |
| CBCL Emotionally Reactive (T1) | - | Bacteria | Bacteroidetes | Bacteroidia |
| CBCL Externalizing Behavior (T1) | - | Bacteria | Bacteroidetes | Bacteroidia |
| CBCL Total (T1) | - | Bacteria | Bacteroidetes | Bacteroidia |
| CBQ Impulsivity (T1) | - | Bacteria | Bacteroidetes | Bacteroidia |
| CBQ Inhibitory Control (T1) | + | Bacteria | Bacteroidetes | Bacteroidia |
| LEC Total | - | Bacteria | Bacteroidetes | Bacteroidia |
| LEC Turmoil | - | Bacteria | Bacteroidetes | Bacteroidia |
| CBCL Aggressive Behavior (T1) | + | Bacteria | Actinobacteria | Actinobacteria |
| CBCL Anxious Depressed (T1) | + | Bacteria | Actinobacteria | Actinobacteria |
| CBCL Depressive Problems (T1) | + | Bacteria | Actinobacteria | Actinobacteria |
| CBCL Externalizing Behavior (T1) | + | Bacteria | Actinobacteria | Actinobacteria |
| CBQ Inhibitory Control (T1) | - | Bacteria | Actinobacteria | Actinobacteria |
| LEC Total | + | Bacteria | Actinobacteria | Actinobacteria |
| LEC Turmoil | + | Bacteria | Actinobacteria | Actinobacteria |
| CBCL Anxious Depressed (T1) | + | Bacteria | Actinobacteria | Actinobacteria |
| CBCL Depressive Problems (T1) | + | Bacteria | Actinobacteria | Actinobacteria |
| CBCL Externalizing Behavior (T1) | + | Bacteria | Actinobacteria | Actinobacteria |
| CBCL Total (T1) | + | Bacteria | Actinobacteria | Actinobacteria |
| CBQ Anger Frustration (T1) | + | Bacteria | Actinobacteria | Actinobacteria |
| LEC Turmoil | + | Bacteria | Actinobacteria | Actinobacteria |
| CBCL Aggressive Behavior (T1) | - | Bacteria | Proteobacteria | Gammaproteobacteria |
| CBCL Anxious Depressed (T1) | - | Bacteria | Proteobacteria | Gammaproteobacteria |
| CBCL Externalizing Behavior (T1) | - | Bacteria | Proteobacteria | Gammaproteobacteria |
| CBCL Total (T1) | - | Bacteria | Proteobacteria | Gammaproteobacteria |
| CBQ Fear (T1) | - | Bacteria | Proteobacteria | Gammaproteobacteria |
| Income to Needs | + | Bacteria | Proteobacteria | Gammaproteobacteria |
| CBCL Anxiety Problems (T1) | - | Bacteria | Bacteroidetes | Bacteroidia |
| CBCL Emotionally Reactive (T1) | - | Bacteria | Bacteroidetes | Bacteroidia |
| CBQ Inhibitory Control (T1) | + | Bacteria | Bacteroidetes | Bacteroidia |

| | | | | |
|---|---|---|---|---|
| Income to Needs | + | Bacteria | Bacteroidetes | Bacteroidia |
| LEC Total | - | Bacteria | Bacteroidetes | Bacteroidia |
| CBCL Depressive Problems (T1) | + | Bacteria | Firmicutes | Clostridia |
| CBCL Emotionally Reactive (T1) | + | Bacteria | Firmicutes | Clostridia |
| CBCL Internalizing Behavior (T1) | + | Bacteria | Firmicutes | Clostridia |
| CBQ Anger Frustration (T1) | + | Bacteria | Firmicutes | Clostridia |
| CBQ Sadness (T1) | + | Bacteria | Firmicutes | Clostridia |
| CBQ Impulsivity (T1) | + | Bacteria | Bacteroidetes | Bacteroidia |
| LEC Family Illness/Injury | + | Bacteria | Bacteroidetes | Bacteroidia |
| LEC Poverty Related Events | + | Bacteria | Bacteroidetes | Bacteroidia |
| LEC Total | + | Bacteria | Bacteroidetes | Bacteroidia |
| CBCL Aggressive Behavior (T1) | + | Bacteria | Firmicutes | Clostridia |
| CBQ Impulsivity (T1) | + | Bacteria | Firmicutes | Clostridia |
| CBQ Inhibitory Control (T1) | - | Bacteria | Firmicutes | Clostridia |
| CBCL Aggressive Behavior (T1) | - | Bacteria | Firmicutes | Negativicutes |
| CBCL Anxious Depressed (T1) | - | Bacteria | Firmicutes | Negativicutes |
| Income to Needs | + | Bacteria | Firmicutes | Negativicutes |
| CBQ Anger Frustration (T1) | + | Bacteria | Firmicutes | Clostridia |
| CBQ Sadness (T1) | + | Bacteria | Firmicutes | Clostridia |
| CBCL Externalizing Behavior (T1) | - | Bacteria | Bacteroidetes | Bacteroidia |
| CBCL Total (T1) | - | Bacteria | Bacteroidetes | Bacteroidia |
| CBCL Anxiety Problems (T1) | + | Bacteria | Proteobacteria | Deltaproteobacteria |
| CBQ Fear (T1) | + | Bacteria | Proteobacteria | Deltaproteobacteria |
| CBCL Depressive Problems (T1) | - | Bacteria | Firmicutes | Clostridia |
| CBCL Internalizing Behavior (T1) | - | Bacteria | Firmicutes | Clostridia |
| CBCL Aggressive Behavior (T1) | - | Bacteria | Firmicutes | Bacilli |
| Income to Needs | + | Bacteria | Firmicutes | Bacilli |
| LEC Total | + | Bacteria | Firmicutes | Clostridia |
| LEC Turmoil | + | Bacteria | Firmicutes | Clostridia |
| CBCL Depressive Problems (T1) | + | Bacteria | Verrucomicrobia | Verrucomicrobiae |
| Income to Needs | + | Bacteria | Firmicutes | Clostridia |
| CBCL Anxious Depressed (T1) | + | Bacteria | Firmicutes | Clostridia |

| | | | | |
|---|---|---|---|---|
| CBQ Inhibitory Control (T1) | - | Bacteria | Firmicutes | Clostridia |
| LEC Family Illness/Injury | + | Bacteria | Bacteroidetes | Bacteroidia |
| Income to Needs | - | Bacteria | Bacteroidetes | Bacteroidia |
| Income to Needs | + | Bacteria | Firmicutes | Clostridia |

xon (assigned to the species level) as determined by CPGLM regression analysis.

| Order | Family | Genus | Species |
|---|---|---|---|
| Bacteroidales | Bacteroidaceae | Bacteroides | Bacteroides_fragilis |
| Bifidobacteriales | Bifidobacteriaceae | Bifidobacterium | Bifidobacterium_adolescentis |
| Coriobacteriales | Coriobacteriaceae | Collinsella | Collinsella_aerofaciens |
| Pasteurellales | Pasteurellaceae | Haemophilus | Haemophilus_parainfluenzae |
| Bacteroidales | Bacteroidaceae | Bacteroides | Bacteroides_dorei |

| | | | |
|---|---|---|---|
| Clostridiales | Clostridiaceae | Clostridium | Clostridium_bolteae |
| Bacteroidales | Rikenellaceae | Alistipes | Alistipes_onderdonkii |
| Clostridiales | Eubacteriaceae | Eubacterium | Eubacterium_siraeum |
| Selenomonadales | Veillonellaceae | Veillonella | Veillonella_unclassified |
| Clostridiales | Ruminococcaceae | Anaerotruncus | Anaerotruncus_colihominis |
| Bacteroidales | Bacteroidaceae | Bacteroides | Bacteroides_thetaiotaomicron |
| Desulfovibrionales | Desulfovibrionaceae | Bilophila | Bilophila_unclassified |
| Clostridiales | Lachnospiraceae | Roseburia | Roseburia_inulinivorans |
| Lactobacillales | Streptococcaceae | Streptococcus | Streptococcus_salivarius |
| Clostridiales | Ruminococcaceae | Subdoligranulum | Subdoligranulum_unclassified |
| Verrucomicrobiales | Verrucomicrobiaceae | Akkermansia | Akkermansia_muciniphila |
| Clostridiales | Lachnospiraceae | Anaerostipes | Anaerostipes_hadrus |
| Clostridiales | Lachnospiraceae | Coprococcus | Coprococcus_comes |
