## Supplementary material for "Gut feelings begin in childhood: how the gut metagenome links to early environment, caregiving, and behavior": Supp Table 7

Supplemental Table 7. Every significant association between an individual covariate and functional group (assigned to the KO level) as determined by CPGLM regression analysis.

| Variable | Unit | Relationshi | Label | See also |
|---|---|---|---|---|
| CBCL Aggressive Behavior | K05914 | + | KO: E1.13.12.7; photinus-luciferin 4-monooxygenase (ATP-hydrolysing) [EC:1.13.12.7] :: PFAMS: AMP-binding enzyme; Condensation domain; Methyltransferase domain; Thioesterase domain; AMP-binding enzyme C-terminal domain | K15665 K15658 K15660 K16130 K12239 K16120 K16129 K16416 K16124 |
| CBCL Emotionally Reactive | K05914 | + | | |
| CBCL Externalizing Behavior | K05914 | + | | |
| CBCL Total | K05914 | + | | |
| CBQ Inhibitory Control | K05914 | - | | |
| amily Separation/Social Services | K05914 | + | | |
| LEC Poverty Related Events | K05914 | + | | |
| LEC Total | K05914 | + | | |
| LEC Turmoil | K05914 | + | | |
| CBCL Aggressive Behavior | K15665 | + | KO: K15665, ppsB, fenD; fengycin family lipopeptide synthetase B :: PFAMS: Condensation domain; AMP-binding enzyme; Phosphopantetheine attachment site; AMP-binding enzyme C-terminal domain; Transferase family | K05914 K15658 K15660 K16130 K12239 K16120 K16129 K16416 K16124 |
| CBCL Anxious Depressed | K15665 | + | | |
| CBCL Emotionally Reactive | K15665 | + | | |
| CBCL Externalizing Behavior | K15665 | + | | |
| CBCL Total | K15665 | + | | |
| CBQ Inhibitory Control | K15665 | - | | |
| LEC Poverty Related Events | K15665 | + | | |
| LEC Total | K15665 | + | | |
| LEC Turmoil | K15665 | + | | |
| CBCL Aggressive Behavior | K11903 | + | MODS: Type VI secretion system | K01220 K01058 |
| CBCL Anxiety Problems | K11903 | + | | |
| CBCL Anxious Depressed | K11903 | + | | |
| CBCL Depressive Problems | K11903 | + | | |
| LEC Turmoil | K11903 | + | | |
| CBCL Anxiety Problems | K11906 | + | | |
| CBCL Depressive Problems | K11906 | + | | |
| CBCL Depressive Problems | K11910 | + | | |
| CBCL Internalizing Behavior | K11910 | + | | |
| CBCL Aggressive Behavior | K05607 | + | MODS: Leucine degradation, leucine => acetoacetate + acetyl-CoA | K00249 |
| CBCL Anxiety Problems | K05607 | + | | |
| CBCL Depressive Problems | K05607 | + | | |
| CBCL Emotionally Reactive | K05607 | + | | |
| CBCL Externalizing Behavior | K05607 | + | | |
| CBCL Internalizing Behavior | K05607 | + | | |
| CBCL Total | K05607 | + | | |
| CBQ Anger Frustration | K05607 | + | | |
| CBCL Aggressive Behavior | K01593 | + | MODS: Melatonin biosynthesis, tryptophan => serotonin => melatonin; Catecholamine biosynthesis, tyrosine => dopamine => noradrenaline => adrenaline | K16314 K04092 K01556 K01658 |
| CBCL Anxiety Problems | K01593 | + | | |
| CBCL Anxious Depressed | K01593 | + | | |
| CBCL Externalizing Behavior | K01593 | + | | |
| CBCL Total | K01593 | + | | |
| CBQ Fear | K01593 | + | | |
| CBQ Inhibitory Control | K01593 | - | | |
| amily Separation/Social Services | K01593 | + | | |
| CBCL Aggressive Behavior | K15658 | + | KO: ofaA, arfA; arthrofactin-type cyclic lipopeptide synthetase A :: PFAMS: AMP-binding enzyme; Condensation domain; Phosphopantetheine attachment site; AMP-binding enzyme C-terminal domain; Periplasmic sensor domain | K15660 K15665 K05914 K15658 K16130 K12239 K16120 K16129 K16416 K16124 |
| CBCL Anxiety Problems | K15658 | + | | |
| CBCL Emotionally Reactive | K15658 | + | | |
| CBCL Externalizing Behavior | K15658 | + | | |
| CBCL Total | K15658 | + | | |
| CBQ Anger Frustration | K15658 | + | | |
| LEC Turmoil | K15658 | + | | |
| CBCL Depressive Problems | K10005 | + | MODS: Glutamate transport system | K05612 K00467 K00228 |
| CBQ Anger Frustration | K10005 | + | | |
| CBCL Depressive Problems | K10006 | + | | |
| CBCL Depressive Problems | K10007 | + | | |
| CBCL Internalizing Behavior | K10007 | + | | |
| CBCL Depressive Problems | K10008 | + | | |
| CBQ Anger Frustration | K10008 | + | | |
| CBCL Anxious Depressed | K01220 | + | KO: E3.2.1.85, lacG; 6-phospho-beta-galactosidase [EC:3.2.1.85] :: PFAMS: Glycosyl hydrolase family 1; Type VI secretion protein IcmF C-terminal; Cellulase (glycosyl hydrolase family 5); Glycosyl hydrolase family 10; Uncharacterized protein conserved in archaea (DUF2095) | K11903 K11906 K11910 K01224 K05991 K01058 |
| CBCL Depressive Problems | K01220 | + | | |
| CBCL Emotionally Reactive | K01220 | + | | |
| CBCL Externalizing Behavior | K01220 | + | | |
| CBCL Total | K01220 | + | | |
| LEC Turmoil | K01220 | + | | |
| CBCL Anxiety Problems | K16180 | + | KO: pylB; methylornithine synthase [EC:5.4.99.58] :: PFAMS: Radical SAM superfamily; 4Fe-4S single cluster domain; Biotin and Thiamin Synthesis associated domain; Pterin binding enzyme; Histidine biosynthesis protein | |
| CBCL Emotionally Reactive | K16180 | + | | |
| amily Separation/Social Services | K16180 | + | | |
| LEC Poverty Related Events | K16180 | + | | |
| LEC Total | K16180 | + | | |
| LEC Turmoil | K16180 | + | | |
| CBCL Depressive Problems | K02074 | + | MODS: Putative zinc/manganese transport system | K16846 K14716 K15726 K15727 |
| CBCL Externalizing Behavior | K02074 | + | | |
| CBCL Internalizing Behavior | K02074 | + | | |
| CBCL Total | K02074 | + | | |
| CBQ Anger Frustration | K02074 | + | | |
| CBCL Depressive Problems | K02077 | + | | |
| CBCL Aggressive Behavior | K01556 | + | MODS: Tryptophan metabolism, tryptophan => kynurenine => 2-aminomuconate | K01593 K01658 |
| CBQ Impulsivity | K01556 | + | | |
| LEC Family Illness/Injury | K01556 | + | | |
| amily Separation/Social Services | K01556 | + | | |
| LEC Poverty Related Events | K01556 | + | | |
| LEC Total | K01556 | + | | |
| CBCL Aggressive Behavior | K04768 | + | KO: acuC; acetoin utilization protein AcuC :: PFAMS: Histone deacetylase domain; B12 binding domain; Glycosyl transferase family 1; Putative aminopeptidase; Monogalactosyldiacylglycerol (MGDG) synthase | K15507 |
| CBCL Anxiety Problems | K04768 | + | | |
| CBCL Externalizing Behavior | K04768 | + | | |
| CBCL Total | K04768 | + | | |
| amily Separation/Social Services | K04768 | + | | |
| CBCL Anxiety Problems | K10725 | + | KO: cdc6A; archaeal cell division control protein 6 :: PFAMS: AAA domain; CDC6; C terminal winged helix domain; AAA ATPase domain; ATPase family associated with various cellular activities (AAA); Bacterial TniB protein | |
| CBCL Anxious Depressed | K10725 | + | | |
| CBCL Emotionally Reactive | K10725 | + | | |
| amily Separation/Social Services | K10725 | + | | |
| LEC Total | K10725 | + | | |
| CBCL Aggressive Behavior | K09138 | + | KO: K09138; uncharacterized protein :: PFAMS: Putative heavy-metal chelation; Protein of unknown function (DUF2478); DNA photolyase; NTPase; Poxvirus nucleic acid binding protein VP8/L4R | K14701 |
| CBCL Externalizing Behavior | K09138 | + | | |
| amily Separation/Social Services | K09138 | + | | |
| LEC Poverty Related Events | K09138 | + | | |
| LEC Total | K09138 | + | | |
| CBCL Aggressive Behavior | K02526 | + | KO: kdgT; 2-keto-3-deoxygluconate permease :: PFAMS: 2-keto-3-deoxygluconate permease; Protein of unknown function (DUF2953); Protein of unknown function (DUF3334); Transition state regulatory protein AbrB; Conserved hypothetical protein 698 | K10820 K10975 K10974 K07243 K14708 K14701 |
| CBCL Externalizing Behavior | K02526 | + | | |
| CBCL Total | K02526 | + | | |
| CBQ Anger Frustration | K02526 | + | | |
| amily Separation/Social Services | K02526 | + | | |
| CBCL Anxious Depressed | K01187 | - | KO: malZ; alpha-glucosidase [EC:3.2.1.20] :: PFAMS: Alpha amylase; catalytic domain; Glycosyl hydrolases family 31; Glycoside hydrolase 97; Galactose mutarotase-like; Glycosyl-hydrolase 97 N-terminal | K01224 K01220 K06113 K09963 K09955 K05988 K05991 K01223 K11931 K15531 K15924 |
| CBCL Depressive Problems | K01187 | - | | |
| CBCL Externalizing Behavior | K01187 | - | | |
| CBCL Total | K01187 | - | | |
| CBQ Inhibitory Control | K01187 | + | | |
| CBCL Anxious Depressed | K05363 | + | KO: murM; serine/alanine adding enzyme [EC:2.3.2.10] :: PFAMS: FemAB family; Acetyltransferase (GNAT) domain; Retrotransposon gag protein; Domain of unknown function (DUF1611_N) Rossmann-like domain; Septation ring formation regulator; EzrA | K03980 |
| CBCL Depressive Problems | K05363 | + | | |
| CBCL Emotionally Reactive | K05363 | + | | |
| CBCL Internalizing Behavior | K05363 | + | | |
| CBCL Total | K05363 | + | | |
| CBCL Anxious Depressed | K10793 | + | KO: prdA; D-proline reductase (dithiol) PrdA [EC:1.21.4.1] :: PFAMS: Glycine/sarcosine/betaine reductase component B subunits; SAF domain; Domain of unknown function (DUF4342); MutS domain I; cAMP-dependent protein kinase inhibitor | K10794 K10795 K10670 |
| CBCL Emotionally Reactive | K10793 | + | | |
| CBCL Total | K10793 | + | | |
| CBQ Anger Frustration | K10793 | + | | |

| | | | |
|---|---|---|---|
| CBQ Fear K10793 | + | KO: prdB; D-proline reductase (dithiol) PrdB [EC:1.21.4.1] :: PFAMS: Glycine/sarcosine/betaine reductase selenoprotein B (GRDB); Glycine/sarcosine/betaine reductase component B subunits; 2-hydroxyglutaryl-CoA dehydratase; D-component; Diacylglycerol kinase catalytic domain; NifU-like domain | K10793 K10795 K10670 |
| CBCL Anxious Depressed K10794 | + | | |
| CBCL Emotionally Reactive K10794 | + | | |
| CBCL Internalizing Behavior K10794 | + | | |
| CBCL Total K10794 | + | | |
| CBQ Anger Frustration K10794 | + | | |
| CBCL Depressive Problems K02452 | + | MODS: Type II general secretion pathway | K09764 K02663 K06218 |
| CBCL Anxious Depressed K02454 | + | | |
| CBCL Depressive Problems K02454 | + | | |
| CBCL Internalizing Behavior K02454 | + | | |
| CBCL Total K02454 | + | | |
| CBCL Aggressive Behavior K16173 | + | KO: acd; glutaryl-CoA dehydrogenase (non-decarboxylating) [EC:1.3.99.32] :: PFAMS: Acyl-CoA dehydrogenase; C-terminal domain; Acyl-CoA dehydrogenase; N-terminal domain; Acyl-CoA dehydrogenase; middle domain; 4-hydroxyphenylacetate 3-hydroxylase N terminal; Domain of unknown function (DUF1966) | K07749 |
| amily Separation/Social Services K16173 | + | | |
| LEC Poverty Related Events K16173 | + | | |
| LEC Total K16173 | + | | |
| amily Separation/Social Services K16753 | + | KO: CCDC34; coiled-coil domain-containing protein 34 :: PFAMS: Domain of unknown function (DUF4207); Borrelia P83/100 protein; Nucleoporin protein Ndc1-Nup; Kinase phosphorylation protein; PPIC-type PPIASE domain | |
| LEC Poverty Related Events K16753 | + | | |
| LEC Total K16753 | + | | |
| LEC Turmoil K16753 | + | | |
| CBQ Impulsivity K13535 | + | KO: CLD1; cardiolipin-specific phospholipase [EC:3.1.1.-] :: PFAMS: alpha/beta hydrolase fold; Alpha/beta hydrolase family; Serine aminopeptidase; S33; Putative esterase; Lipid-droplet associated hydrolase | K01114 K01058 |
| LEC Family Illness/Injury K13535 | + | | |
| amily Separation/Social Services K13535 | + | | |
| LEC Total K13535 | + | | |
| amily Separation/Social Services K01224 | - | KO: E3.2.1.89; arabinogalactan endo-1,4-beta-galactosidase [EC:3.2.1.89] :: PFAMS: Glycosyl hydrolase family 53; Cellulase (glycosyl hydrolase family 5); Glycosyl hydrolase catalytic core; Bacterial Ig-like domain (group 4); Ricin-type beta-trefoil lectin domain-like | K11903 K11906 K11910 K01220 K05991 K01058 |
| LEC Poverty Related Events K01224 | - | | |
| LEC Total K01224 | - | | |
| LEC Turmoil K01224 | - | | |
| CBCL Aggressive Behavior K11420 | + | KO: EHMT; euchromatic histone-lysine N-methyltransferase [EC:2.1.1.43] :: PFAMS: SAD/SRA domain; SET domain; Ankyrin repeats (many copies); Ankyrin repeat; Pre-SET motif | K09186 K11428 |
| CBCL Anxiety Problems K11420 | + | | |
| CBCL Anxious Depressed K11420 | + | | |
| CBCL Depressive Problems K11420 | + | | |
| CBCL Emotionally Reactive K12471 | + | KO: EPN; epsin :: PFAMS: ENTH domain; ANTH domain; VHS domain; Ubiquitin interaction motif; Domain of unknown function (DUF1720) | |
| CBCL Externalizing Behavior K12471 | + | | |
| CBCL Total K12471 | + | | |
| C Unsafe/Violent Neighborhood K12471 | + | | |
| CBCL Anxious Depressed K00443 | + | KO: frhG; coenzyme F420 hydrogenase subunit gamma [EC:1.12.98.1] :: PFAMS: 4Fe-4S dicluster domain; 4Fe-4S binding domain; NADH ubiquinone oxidoreductase; 20 Kd subunit; 4Fe-4S double cluster binding domain; 4Fe-4S single cluster domain of Ferredoxin I | |
| CBCL Depressive Problems K00443 | + | | |
| CBCL Internalizing Behavior K00443 | + | | |
| CBCL Total K00443 | + | | |
| CBCL Aggressive Behavior K06294 | + | KO: gerD; spore germination protein D :: PFAMS: Death-like domain of SPT6; Insect allergen related repeat; nitrile-specifier detoxification; Protein of unknown function (DUF445); Primosomal protein DnaI N-terminus; UCH-binding domain | K06296 |
| CBCL Anxiety Problems K06294 | + | | |
| CBCL Emotionally Reactive K06294 | + | | |
| CBQ Anger Frustration K06294 | + | | |
| CBQ Impulsivity K14830 | - | KO: MAK11, PAK1IP1; protein MAK11 :: PFAMS: WD domain; G-beta repeat; Anaphase-promoting complex subunit 4 WD40 domain; Nucleoporin Nup120/160; WD40-like domain; Nup133 N terminal like | |
| CBQ Inhibitory Control K14830 | + | | |
| LEC Total K14830 | - | | |
| C Unsafe/Violent Neighborhood K14830 | - | | |
| CBCL Emotionally Reactive K01820 | + | KO: rhaA; L-rhamnose isomerase / sugar isomerase [EC:5.3.1.14 5.3.1.-] :: PFAMS: Xylose isomerase-like TIM barrel; L-rhamnose isomerase (RhaA); SIS domain; L-fucose isomerase; C-terminal domain; Protein of unknown function (DUF1428) | |
| CBCL Internalizing Behavior K01820 | + | | |
| CBCL Total K01820 | + | | |
| CBQ Anger Frustration K01820 | + | | |
| CBCL Anxiety Problems K06314 | + | KO: rsfA; prespore-specific regulator :: PFAMS: Myb-like DNA-binding domain; Septum formation initiator; Cell division protein ZapB; IncA protein; Autophagy protein Apg6 | |
| CBCL Anxious Depressed K06314 | + | | |
| CBCL Depressive Problems K06314 | + | | |
| LEC Turmoil K06314 | + | | |
| CBCL Aggressive Behavior K16846 | + | KO: suyB; (2R)-sulfolactate sulfo-lyase subunit beta [EC:4.4.1.24] :: PFAMS: D-galactarate dehydratase / Altronate hydrolase; C terminus; Zinc-uptake complex component A periplasmic; Family of unknown function (DUF5363); WHEP-TRS domain; Domain of unknown function (DUF4124) | K02074 K14716 |
| CBCL Emotionally Reactive K16846 | + | | |
| CBCL Externalizing Behavior K16846 | + | | |
| CBQ Anger Frustration K16846 | + | | |
| CBCL Anxiety Problems K10107 | + | MODS: Capsular polysaccharide transport system | |
| CBCL Anxious Depressed K10107 | + | | |
| CBCL Emotionally Reactive K10107 | + | | |
| CBQ Anger Frustration K10107 | + | | |
| CBCL Anxious Depressed K00665 | + | MODS: Fatty acid biosynthesis, elongation; Fatty acid biosynthesis, initiation | K10255 K10680 K09940 |
| CBCL Depressive Problems K00665 | - | | |
| LEC Poverty Related Events K00665 | + | | |
| LEC Turmoil K00665 | + | | |
| CBQ Impulsivity K06113 | - | KO: abnA; arabinan endo-1,5-alpha-L-arabinosidase [EC:3.2.1.99] :: PFAMS: Glycosyl hydrolases family 43; Ricin-type beta-trefoil lectin domain-like; Glycosyl hydrolases family 32 N-terminal domain; Levansucrase/Invertase; Ricin-type beta-trefoil lectin domain | K01224 K01220 K01187 K09963 K09955 K05988 K05991 K01223 K11931 K15531 K15924 |
| CBQ Inhibitory Control K06113 | + | | |
| LEC Total K06113 | - | | |
| CBCL Aggressive Behavior K01483 | + | KO: allA; ureidoglycolate lyase [EC:4.3.2.3] :: PFAMS: Ureidoglycolate lyase; P53 transactivation motif; Domain of unknown function (DUF4867); Protein of unknown function (DUF1357); Ubiquitin carboxyl-terminal hydrolase; family 1 | |
| CBCL Anxious Depressed K01483 | + | | |
| CBCL Depressive Problems K01483 | + | | |
| CBQ Impulsivity K11497 | + | KO: CENPC; centromere protein C :: PFAMS: Mif2/CENP-C like; Kinetochore assembly subunit CENP-C N-terminal; Centromere assembly component CENP-C middle DNMT3B-binding region; Kinetochore CENP-C fungal homologue; Mif2; N-terminal; Cupin domain | |
| amily Separation/Social Services K11497 | + | | |
| LEC Poverty Related Events K11497 | + | | |
| CBCL Emotionally Reactive K10643 | + | KO: CNOT4, NOT4, MOT2; CCR4-NOT transcription complex subunit 4 [EC:2.3.2.27] :: PFAMS: RING/Ubox like zinc-binding domain; Zinc finger; C3HC4 type (RING finger); RNA recognition motif. (a.k.a. RRM; RBD; or RNP domain); Ring finger domain; Prokaryotic RING finger family 4 | |
| CBQ Anger Frustration K10643 | + | | |
| C Unsafe/Violent Neighborhood K10643 | + | | |
| CBCL Anxiety Problems K10820 | + | KO: E3.6.3.17; monosaccharide-transporting ATPase [EC:3.6.3.17] :: PFAMS: Branched-chain amino acid transport system / permease component; Periplasmic binding protein domain; ABC transporter; AAA domain; Domain of unknown function (DUF4432) | K02526 K10975 K10974 K07243 K14708 K14701 |
| CBCL Anxious Depressed K10820 | + | | |
| CBCL Depressive Problems K10820 | + | | |
| CBCL Anxiety Problems K01509 | + | KO: ENTPD2; adenosinetriphosphatase [EC:3.6.1.3] :: PFAMS: GDA1/CD39 (nucleoside phosphatase) family; Dynein heavy chain and region D6 of dynein motor; SNF2 family N-terminal domain; Helicase conserved C-terminal domain; DEAD/DEAH box helicase | |
| CBCL Anxious Depressed K01509 | + | | |
| CBCL Depressive Problems K01509 | + | | |
| CBCL Aggressive Behavior K16364 | + | KO: GLDN; GLDN; gliomedin :: PFAMS: Olfactomedin-like domain; Collagen triple helix repeat (20 copies); Chorismate lyase; RNA 2'-O ribose methyltransferase substrate binding; BNR repeat-containing family member | |
| LEC Family Illness/Injury K16364 | + | | |
| amily Separation/Social Services K16364 | + | | |
| CBCL Aggressive Behavior K08972 | + | KO: K08972; putative membrane protein :: PFAMS: Mycobacterial 4 TMS phage holin; superfamily IV; Protein of unknown function (DUF1700); Transglycosylase associated protein; OST3 / OST6 family; transporter family; Uncharacterized protein conserved in bacteria (DUF2062) | |
| CBCL Depressive Problems K08972 | + | | |
| CBCL Total K08972 | + | | |
| CBCL Depressive Problems K09963 | + | KO: K09963; uncharacterized protein :: PFAMS: Bacterial protein of unknown function (DUF871); Cellulase (glycosyl hydrolase family 5); GxGYxYP putative glycoside hydrolase C-terminal domain; Tetrahydromethanopterin S-methyltransferase subunit B; OstA-like protein | K01224 K01220 K06113 K01187 K09955 K05988 K05991 K01223 K11931 K15531 K15924 |
| CBCL Internalizing Behavior K09963 | + | | |
| CBQ Anger Frustration K09963 | + | | |
| amily Separation/Social Services K06895 | + | KO: lysE, argO; L-lysine exporter family protein LysE/ArgO :: PFAMS: LysE type translocator; Protein of unknown function; DUF485; CAAD domains of cyanobacterial aminoacyl-tRNA synthetase; Integral membrane protein TerC family; MarC family integral membrane protein | |
| LEC Total K06895 | + | | |
| LEC Turmoil K06895 | + | | |
| LEC Poverty Related Events K09037 | + | KO: MAFF_G_K; transcription factor MAFF/G/K :: PFAMS: bZIP Maf transcription factor; bZIP transcription factor; Septum formation initiator; Protein of unknown function (DUF1192); Basic region leucine zipper | |
| LEC Total K09037 | + | | |
| C Unsafe/Violent Neighborhood K09037 | + | | |
| CBCL Aggressive Behavior K05565 | + | KO: mnhA, mrpA; multicomponent Na+:H+ antiporter subunit A :: PFAMS: Proton-conducting membrane transporter; Domain of unknown function (DUF4040); NADH-Ubiquinone oxidoreductase (complex I); chain 5 N-terminus; Domain related to MnhB subunit of Na+/H+ antiporter; Predicted membrane protein (DUF2108) | K05568 K05570 |
| CBCL Emotionally Reactive K05565 | + | | |
| CBQ Anger Frustration K05565 | + | | |
| CBCL Depressive Problems K15660 | + | KO: ofaC, arfC; arthrofactin-type cyclic lipopeptide synthetase C :: PFAMS: AMP-binding enzyme; Condensation domain; Phosphopantetheine attachment site; Thioesterase domain; AMP-binding enzyme C-terminal domain | K15665 K15658 K05914 K16130 K12239 K16120 K16129 K16416 K16124 |
| CBCL Emotionally Reactive K15660 | + | | |
| CBQ Anger Frustration K15660 | + | | |
| LEC Poverty Related Events K01027 | + | KO: OXCT; 3-oxoacid CoA-transferase [EC:2.8.3.5] :: PFAMS: Coenzyme A transferase; Acetyl-CoA hydrolase/transferase C-terminal domain; Malonate decarboxylase; alpha subunit; transporter; CoA-transferase family III; Acetyl-CoA hydrolase/transferase N-terminal domain | K01029 |
| LEC Total K01027 | + | | |
| LEC Turmoil K01027 | + | | |
| CBCL Aggressive Behavior K09251 | + | KO: patA; putrescine aminotransferase [EC:2.6.1.82] :: PFAMS: Aminotransferase class-III; Aminotransferase class I and II; | |

| | | | |
|---|---|---|---|
| CBQ Anger Frustration K09251<br>amily Separation/Social Services K09251 | +<br>+ | KO: patA; putrescine aminotransferase [EC:2.6.1.82] :: PFAMS: Aminotransferase class-III; Aminotransferase class I and II; Aminotransferase class-V; Oxidoreductase family; NAD-binding Rossmann fold; Beta-eliminating lyase | K13252 |
| CBCL Aggressive Behavior K01114<br>CBCL Externalizing Behavior K01114<br>CBQ Anger Frustration K01114 | -<br>-<br>- | KO: plc; phospholipase C [EC:3.1.4.3] :: PFAMS: Phosphoesterase family; Domain of unknown function (DUF756); Zinc dependent phospholipase C; TAT (twin-arginine translocation) pathway signal sequence; Endonuclease/Exonuclease/phosphatase family | K13535 K01058 |
| CBCL Depressive Problems K09020<br>CBCL Internalizing Behavior K09020<br>CBCL Total K09020 | +<br>+<br>+ | KO: rutB; ureidoacrylate peracid hydrolase [EC:3.5.1.110] :: PFAMS: Isochorismatase family; Carbon-nitrogen hydrolase; NYN domain; Protein of unknown function (DUF4232); Domain of unknown function (DUF5059) | |
| CBCL Anxious Depressed K05612<br>CBCL Externalizing Behavior K05612<br>CBCL Total K05612 | +<br>+<br>+ | KO: SLC1A1, EAAT3; solute carrier family 1 (neuronal/epithelial high affinity glutamate transporter), member 1 :: PFAMS: Sodium:dicarboxylate symporter family; Phage holin T7 family; holin superfamily II; Protein of unknown function (DUF3094); Serpentine type 7TM GPCR chemoreceptor Sra; Protein of unknown function (DUF2730) | K10005 K00467 K00228 |
| amily Separation/Social Services K06665<br>LEC Poverty Related Events K06665<br>LEC Total K06665 | +<br>+<br>+ | KO: SSN6, CYC8; general transcriptional corepressor CYC8 :: PFAMS: Tetratricopeptide repeat; TPR repeat; Anaphase-promoting complex; cyclosome; subunit 3; Tetratricopeptide repeat-like domain; ChAPs (Chs5p-Arf1p-binding proteins) | |
| CBCL Aggressive Behavior K08357<br>CBCL Externalizing Behavior K08357<br>amily Separation/Social Services K08357 | +<br>+<br>+ | KO: ttrA; tetrathionate reductase subunit A :: PFAMS: Molybdopterin oxidoreductase; Molydopterin dinucleotide binding domain; Molybdopterin oxidoreductase Fe4S4 domain; TAT (twin-arginine translocation) pathway signal sequence; NADH dehydrogenase [ubiquinone] 1 subunit C1; mitochondrial | K00183 K00397 |
| CBCL Aggressive Behavior K00249<br>CBCL Externalizing Behavior K00249<br>CBQ Anger Frustration K00249 | +<br>+<br>+ | MODS: beta-Oxidation; Malonate semialdehyde pathway, propanoyl-CoA => acetyl-CoA; Leucine degradation, leucine => acetoacetate + acetyl-CoA | K05607 K13929 K13932 K01027 |
| CBCL Anxious Depressed K04345<br>CBCL Depressive Problems K04345<br>CBCL Anxious Depressed K08049 | +<br>+<br>+ | MODS: cAMP signaling | |
| CBQ Anger Frustration K05813<br>CBQ Anger Frustration K05814<br>CBQ Anger Frustration K05815 | +<br>+<br>+ | MODS: Putative sn-glycerol-phosphate transport system | |
| CBCL Depressive Problems K10939<br>amily Separation/Social Services K10939 | +<br>+ | KO: acfD; accessory colonization factor AcfD :: PFAMS: Domain of unknown function (DUF4092); Peptidase M60; enhancin and enhancin-like; N-terminal domain of M60-like peptidases; Procyclic acidic repetitive protein (PARP); PQQ enzyme | |
| CBCL Anxious Depressed K01444<br>CBCL Internalizing Behavior K01444 | +<br>+ | KO: AGA, aspG; N4-(beta-N-acetylglucosaminyl)-L-asparaginase [EC:3.5.1.26] :: PFAMS: Asparaginase; TAT (twin-arginine translocation) pathway signal sequence; RED-like protein N-terminal region; RED-like protein C-terminal region; | |
| CBCL Aggressive Behavior K06254<br>CBCL Externalizing Behavior K06254 | +<br>+ | KO: AGRN; agrin :: PFAMS: Laminin G domain; Kazal-type serine protease inhibitor domain; Agrin NtA domain; Laminin EGF domain; EGF-like domain | |
| CBCL Externalizing Behavior K15504<br>C Unsafe/Violent Neighborhood K15504 | +<br>+ | KO: ANKRD52; serine/threonine-protein phosphatase 6 regulatory ankyrin repeat subunit C :: PFAMS: Ankyrin repeats (many copies); Ankyrin repeat; Ankyrin repeats (3 copies); Glutamine amidotransferase domain; PEGA domain | K16314 |
| CBCL Anxiety Problems K08117<br>C Unsafe/Violent Neighborhood K08117 | +<br>+ | KO: APLP2; amyloid-like protein 2 :: PFAMS: E2 domain of amyloid precursor protein; Amyloid A4 N-terminal heparin-binding; beta-amyloid precursor protein C-terminus; Copper-binding of amyloid precursor; CuBD; Kunitz/Bovine | |
| CBCL Depressive Problems K11653<br>CBQ Anger Frustration K11653 | +<br>+ | KO: ARID1; AT-rich interactive domain-containing protein 1 :: PFAMS: SWI/SNF-like complex subunit BAF250/Osa; ARID/BRIGHT DNA binding domain; Protein of unknown function (DUF1510); DNA polymerase phi; SDA1 | |
| CBQ Impulsivity K14201<br>CBQ Inhibitory Control K14201 | +<br>- | KO: clfA; clumping factor A :: PFAMS: C-terminus of bacterial fibrinogen-binding adhesin; YSIRK type signal peptide; LPXTG cell wall anchor motif; Lamina-associated polypeptide 1C (LAP1C); FAM196 family | K14192 |
| CBQ Anger Frustration K13540<br>CBQ Sadness K13540 | +<br>+ | KO: cobIJ; precorrin-2 C20-methyltransferase / precorrin-3B C17-methyltransferase [EC:2.1.1.130 2.1.1.131] :: PFAMS: Tetrapyrrole (Corrin/Porphyrin) Methylases; Radical SAM superfamily; Lamina-associated polypeptide 2 alpha; Domain of | K16329 |
| CBCL Depressive Problems K01753<br>CBQ Anger Frustration K01753 | +<br>+ | KO: dsdA; D-serine dehydratase [EC:4.3.1.18] :: PFAMS: Pyridoxal-phosphate dependent enzyme; Putative serine dehydratase domain; Alanine racemase; N-terminal domain; NmrA-like family; Domain of unknown function (DUF4123) | |
| CBCL Depressive Problems K00467<br>LEC Turmoil K00467 | +<br>+ | KO: E1.13.12.4; lactate 2-monooxygenase [EC:1.13.12.4] :: PFAMS: FMN-dependent dehydrogenase; Thiazole biosynthesis protein ThiG; IMP dehydrogenase / GMP reductase domain; Conserved region in glutamate synthase; Nitronate | K05612 K10005 K00228 |
| CBQ Anger Frustration K01029<br>amily Separation/Social Services K01029 | +<br>+ | KO: E2.8.3.5B, scoB; 3-oxoacid CoA-transferase subunit B [EC:2.8.3.5] :: PFAMS: Coenzyme A transferase; Acetyl-CoA hydrolase/transferase C-terminal domain; Initiation factor 2 subunit family; DeoR C terminal sensor domain; Citrate lyase; | K01027 |
| LEC Poverty Related Events K14091<br>LEC Total K14091 | +<br>+ | KO: echF; ech hydrogenase subunit F :: PFAMS: 4Fe-4S dicluster domain; 4Fe-4S binding domain; 4Fe-4S double cluster binding domain; 4Fe-4S single cluster domain; Electron transfer flavoprotein-ubiquinone oxidoreductase; 4Fe-4S | K14087 |
| LEC Family Illness/Injury K11462<br>LEC Total K11462 | +<br>+ | KO: EED; polycomb protein EED :: PFAMS: WD domain; G-beta repeat; Anaphase-promoting complex subunit 4 WD40 domain; Nucleoporin Nup120/160; WD40 region of Ge1; enhancer of mRNA-decapping protein; Double-stranded RNA | |
| CBQ Anger Frustration K00124<br>CBQ Fear K00124 | +<br>+ | KO: fdoH; formate dehydrogenase iron-sulfur subunit :: PFAMS: 4Fe-4S dicluster domain; 4Fe-4S binding domain; Formate dehydrogenase N; transmembrane; Respiratory-chain NADH dehydrogenase 51 Kd subunit; 4Fe-4S double cluster binding | |
| CBCL Aggressive Behavior K07749<br>amily Separation/Social Services K07749 | +<br>+ | KO: frc; formyl-CoA transferase [EC:2.8.3.16] :: PFAMS: CoA-transferase family III; DeoC/LacD family aldolase; Putative antitoxin of bacterial toxin-antitoxin system; YdaS/YdaT; LEM domain; T-cell surface glycoprotein CD3 zeta chain | K16173 |
| CBCL Anxious Depressed K08313<br>CBCL Depressive Problems K08313 | +<br>+ | KO: fsaA, mipB; fructose-6-phosphate aldolase 1 [EC:4.1.2.-] :: PFAMS: Transaldolase/Fructose-6-phosphate aldolase; Protein of unknown function (DUF3549); Histidine biosynthesis protein; Thiazole biosynthesis protein ThiG; Domain of | |
| LEC Poverty Related Events K01630<br>LEC Total K01630 | +<br>+ | KO: garL; 2-dehydro-3-deoxyglucarate aldolase [EC:4.1.2.20] :: PFAMS: HpcH/HpaI aldolase/citrate lyase family; Pyruvate kinase; barrel domain; PEP-utilising enzyme; TIM barrel domain; BpuSI N-terminal domain; B12 binding domain | |
| CBCL Internalizing Behavior K03746<br>CBCL Total K03746 | +<br>+ | KO: hns; DNA-binding protein H-NS :: PFAMS: H-NS histone family; Protein of unknown function (DUF3037); PaaX-like protein C-terminal domain; Swi5; HAUS augmin-like complex subunit 3 | |
| LEC Family Illness/Injury K06662<br>C Unsafe/Violent Neighborhood K06662 | +<br>+ | KO: HRAD17, RAD24; cell cycle checkpoint protein :: PFAMS: Rad17 cell cycle checkpoint protein; AAA domain; AAA ATPase domain; ATPase family associated with various cellular activities (AAA); ABC transporter | K06632 |
| CBCL Internalizing Behavior K07461<br>CBCL Total K07461 | +<br>+ | KO: K07461; putative endonuclease :: PFAMS: GIY-YIG catalytic domain; T5orf172 domain; Meiotically up-regulated gene 113; Methyltransferase domain; Arm DNA-binding domain | |
| CBQ Impulsivity K09955<br>CBQ Inhibitory Control K09955 | -<br>+ | KO: K09955; uncharacterized protein :: PFAMS: Beta-L-arabinofuranosidase; GH127; Domain of unknown function; Glycosyl Hydrolase Family 88; Glycosyl hydrolase family 76; Ricin-type beta-trefoil lectin domain-like | K01224 K01220 K06113 K09963 K01187 K05988 K05991 K01223 K11931 K15531 K15924 |
| LEC Poverty Related Events K02528<br>LEC Total K02528 | +<br>+ | KO: ksgA; 16S rRNA (adenine1518-N6/adenine1519-N6)-dimethyltransferase [EC:2.1.1.182] :: PFAMS: Ribosomal RNA adenine dimethylase; Methyltransferase domain; Nodulation protein S (NodS); Methyltransferase small domain; Putative | |
| amily Separation/Social Services K05898<br>LEC Total K05898 | +<br>+ | KO: kstD; 3-oxosteroid 1-dehydrogenase [EC:1.3.99.4] :: PFAMS: FAD binding domain; FAD dependent oxidoreductase; Pyridine nucleotide-disulphide oxidoreductase; HI0933-like protein; NAD(P)-binding Rossmann-like domain | |
| CBCL Anxiety Problems K16130<br>CBCL Emotionally Reactive K16130 | +<br>+ | KO: mcyA; microcystin synthetase protein McyA :: PFAMS: AMP-binding enzyme; Condensation domain; Methyltransferase domain; Phosphopantetheine attachment site; AMP-binding enzyme C-terminal domain | K15665 K15658 K15660 K05914 K12239 K16120 K16129 K16416 K16124 |
| CBQ Anger Frustration K03750<br>CBQ Fear K03750 | +<br>+ | KO: moeA; molybdopterin molybdotransferase [EC:2.10.1.1] :: PFAMS: MoeA N-terminal region (domain I and II); Probable molybdopterin binding domain; MoeA C-terminal region (domain IV); Molybdopterin guanine dinucleotide | |
| CBCL Depressive Problems K03980<br>CBQ Anger Frustration K03980 | +<br>+ | KO: murJ, mviN; putative peptidoglycan lipid II flippase :: PFAMS: MviN-like protein; Polysaccharide biosynthesis C-terminal domain; MatE; Polysaccharide biosynthesis protein; Apolipoprotein O | K05363 K03327 |
| CBCL Total K14552<br>CBQ Sadness K14552 | -<br>- | KO: NAN1, UTP17, WDR75; NET1-associated nuclear protein 1 (U3 small nucleolar RNA-associated protein 17) :: PFAMS: WD domain; G-beta repeat; Anaphase-promoting complex subunit 4 WD40 domain; WD40-like Beta Propeller Repeat; | |
| CBCL Anxious Depressed K12239<br>CBCL Depressive Problems K12239 | +<br>+ | KO: pchE; dihydroaeruginoic acid synthetase :: PFAMS: AMP-binding enzyme; Condensation domain; Phosphopantetheine attachment site; Methyltransferase domain; AMP-binding enzyme C-terminal domain | K15665 K15658 K15660 K16130 K05914 K16120 K16129 K16416 K16124 |
| CBQ Anger Frustration K06875<br>CBQ Sadness K06875 | +<br>+ | KO: PDCD5, TFAR19; programmed cell death protein 5 :: PFAMS: Double-stranded DNA-binding domain; PCI domain; Eukaryotic elongation factor 1 beta central acidic region; Radial spoke protein 3; Domain of unknown function (DUF4615) | |
| CBCL Externalizing Behavior K13923<br>LEC Turmoil K13923 | +<br>+ | KO: pduL; phosphate propanoyltransferase [EC:2.3.1.222] :: PFAMS: Phosphate propanoyltransferase; Molydopterin dinucleotide binding domain; Antidote-toxin recognition MazE; bacterial antitoxin; Gill-associated viral 3C-like peptidase; | |
| CBCL Anxiety Problems K11935<br>CBCL Depressive Problems K11935 | +<br>+ | KO: pgaA; biofilm PGA synthesis protein PgaA :: PFAMS: Tetratricopeptide repeat; TPR repeat; Anaphase-promoting complex; cyclosome; subunit 3; Protein of unknown function (DUF560); Coatomer epsilon subunit | K11931 |
| CBCL Anxious Depressed K11456<br>CBCL Depressive Problems K11456 | +<br>- | KO: PHC1, EDR1; polyhomeotic-like protein 1 :: PFAMS: SAM domain (Sterile alpha motif); Unstructured region on Polyhomeotic-like protein 1 and 2; Sterile alpha motif (SAM)/Pointed domain; CGNR zinc finger; MYM-type Zinc finger | |
| CBCL Anxious Depressed K10795<br>CBCL Total K10795 | +<br>+ | KO: prdD; D-proline reductase (dithiol)-stabilizing protein PrdD :: PFAMS: Glycine/sarcosine/betaine reductase component B subunits | K10793 K10794 K10670 |
| CBCL Depressive Problems K01354<br>CBQ Anger Frustration K01354 | +<br>+ | KO: ptrB; oligopeptidase B [EC:3.4.21.83] :: PFAMS: Prolyl oligopeptidase; N-terminal beta-propeller domain; Prolyl oligopeptidase family; alpha/beta hydrolase fold; Acetyl xylan esterase (AXE1); X-Pro dipeptidyl-peptidase (S15 family) | |
| LEC Poverty Related Events K10839<br>LEC Total K10839 | +<br>+ | KO: RAD23, HR23; UV excision repair protein RAD23 :: PFAMS: UBA/TS-N domain; XPC-binding domain; Ubiquitin family; Ubiquitin-2 like Rad60 SUMO-like; Fungal ubiquitin-associated domain | K03631 K03515 K03660 K01249 |
| CBCL Aggressive Behavior K03631<br>CBCL Externalizing Behavior K03631 | +<br>+ | KO: recN; DNA repair protein RecN (Recombination protein N) :: PFAMS: AAA domain; RecF/RecN/SMC N terminal domain; AAA ATPase domain; AAA domain; putative AbiEii toxin; Type IV TA system; P-loop containing region of AAA | K10839 K03515 K03660 K01249 |
| CBCL Total K03515<br>CBQ Anger Frustration K03515 | -<br>- | KO: REV1; DNA repair protein REV1 [EC:2.7.7.-] :: PFAMS: impB/mucB/samB family; impB/mucB/samB family C-terminal domain; DNA repair protein REV1 C-terminal domain; BRCT domain; a BRCA1 C-terminus domain; Domain of unknown | |
| CBCL Aggressive Behavior K15471<br>CBQ Anger Frustration K15471 | +<br>+ | KO: rhiI; O-methyltransferase [EC:2.1.1.-] :: PFAMS: Methyltransferase domain; ubiE/COQ5 methyltransferase family; Hypothetical methyltransferase; FtsJ-like methyltransferase; Mycolic acid cyclopropane synthetase | K00599 K07442 |
| CBCL Anxious Depressed K01160<br>CBCL Depressive Problems K01160 | +<br>+ | KO: rusA; crossover junction endodeoxyribonuclease RusA [EC:3.1.22.4] :: PFAMS: Endodeoxyribonuclease RusA; Domain of unknown function (DUF3368); SLBB domain; Protein of unknown function (DUF3168); SecA preprotein cross-linking | |
| CBQ Impulsivity K14649<br>amily Separation/Social Services K14649 | +<br>+ | KO: TAF8; transcription initiation factor TFIID subunit 8 :: PFAMS: Transcription factor TFIID complex subunit 8 C-term; Bromodomain associated; Transcription initiation factor IID; 31kD subunit; Histone-fold protein; CENP-S protein | |
| CBCL Depressive Problems K10805<br>CBCL Internalizing Behavior K10805 | +<br>+ | KO: tesB; acyl-CoA thioesterase II [EC:3.1.2.-] :: PFAMS: Thioesterase-like superfamily; Acyl-CoA thioesterase; Thioesterase superfamily; Protein of unknown function (DUF3181); Domain of unknown function (DUF1957) | |

| | | | |
|---|---|---|---|
| CBCL Anxious Depressed K03153 | - | KO: thiO; glycine oxidase [EC:1.4.3.19] :: PFAMS: FAD dependent oxidoreductase; Pyridine nucleotide-disulphide oxidoreductase; FAD binding domain; NAD(P)-binding Rossmann-like domain; HI0933-like protein | |
| CBCL Depressive Problems K03153 | + | | |
| CBCL Aggressive Behavior K15712 | + | KO: TTC3; E3 ubiquitin-protein ligase TTC3 [EC:2.3.2.27] :: PFAMS: Tetratricopeptide repeat; Zinc finger; C3HC4 type (RING finger); Ring finger domain; RING-H2 zinc finger domain; TPR repeat | K10639 K10661 K15710 |
| CBCL Externalizing Behavior K15712 | + | | |
| CBCL Depressive Problems K03623 | + | KO: yhcO; ribonuclease inhibitor :: PFAMS: Barstar (barnase inhibitor); Domain of unknown function (DUF4391); Domain of unknown function (DUF3903); EspG family; Mitochondrial import protein Pam17 | |
| CBQ Anger Frustration K03623 | + |  |  |
| LEC Family Illness/Injury K07680 | + | MODS: ComP-ComA (competence) two-component regulatory system | |
| amily Separation/Social Services K07680 | + | | |
| CBQ Fear K02567 | + | MODS: Denitrification, nitrate => nitrogen; Dissimilatory nitrate reduction, nitrate => ammonia | K04561 K00362 |
| C Unsafe/Violent Neighborhood K02568 | - | | |
| CBCL Depressive Problems K01595 | + | MODS: Formaldehyde assimilation, serine pathway; Dicarboxylate-hydroxybutyrate cycle; C4-dicarboxylic acid cycle, phosphoenolpyruvate carboxykinase type; Reductive citrate cycle (Arnon-Buchanan cycle); C4-dicarboxylic acid cycle, | K00024 |
| CBCL Internalizing Behavior K01595 | + | | |
| CBCL Depressive Problems K10037 | + | MODS: Glutamine transport system | |
| CBCL Internalizing Behavior K10037 | + | | |
| CBCL Externalizing Behavior K11519 | - | MODS: HRD1/SEL1 ERAD complex | |
| CBCL Total K11519 | - |  |  |
| C Unsafe/Violent Neighborhood K00888 | + | MODS: Inositol phosphate metabolism, PI=> PIP2 => Ins(1,4,5)P3 => Ins(1,3,4,5)P4 | |
| LEC Turmoil K05858 | + | | |
| CBQ Inhibitory Control K02548 | + | MODS: Menaquinone biosynthesis, chorismate => menaquinone | K06998 K01658 K04092 |
| CBQ Anger Frustration K02552 | + |  |  |
| CBQ Anger Frustration K14083 | + | MODS: Methanogenesis, methylamine/dimethylamine/trimethylamine => methane | K13788 K04480 |
| CBCL Emotionally Reactive K16176 | + | | |
| CBCL Anxiety Problems K11742 | + | MODS: Multidrug resistance, efflux pump MdtIJ | K03327 K11939 K03577 |
| CBCL Depressive Problems K11742 | + | | |
| LEC Family Illness/Injury K10200 | + | MODS: N-Acetylglucosamine transport system | |
| amily Separation/Social Services K10201 | + | | |
| CBQ Inhibitory Control K07636 | + | MODS: PhoR-PhoB (phosphate starvation response) two-component regulatory system | |
| CBQ Inhibitory Control K07657 | + | | |
| CBCL Aggressive Behavior K01848 | + | MODS: Propanoyl-CoA metabolism, propanoyl-CoA => succinyl-CoA; Hydroxypropionate-hydroxybutylate cycle; 3-Hydroxypropionate bi-cycle | K00249 |
| CBCL Externalizing Behavior K01848 | + | | |
| CBCL Anxious Depressed K10984 | + | MODS: PTS system, galactosamine-specific II component | K02788 K02799 K02810 |
| CBCL Anxious Depressed K10985 | + | | |
| CBQ Impulsivity K15598 | + | MODS: Putative hydroxymethylpyrimidine transport system | |
| CBQ Impulsivity K15599 | + | | |
| CBQ Fear K01497 | + | MODS: Riboflavin biosynthesis, GTP => riboflavin/FMN/FAD | |
| CBQ Impulsivity K11752 | - | | |
| CBQ Impulsivity K02936 | + | MODS: Ribosome, eukaryotes; Ribosome, archaea | K02942 |
| LEC Family Illness/Injury K02936 | + | | |
| CBQ Anger Frustration K08300 | + | MODS: RNA degradosome |  |
| CBQ Sadness K08300 | + | | |
| CBCL Depressive Problems K08479 | - | MODS: SasA-RpaAB (circadian timing mediating) two-component regulatory system | |
| CBQ Inhibitory Control K08479 | + | | |
| LEC Family Illness/Injury K00654 | + | MODS: Sphingosine biosynthesis; Ceramide biosynthesis | K04718 |
| LEC Total K00654 | + | | |
| CBCL Depressive Problems K04718 | + | MODS: Sphingosine degradation | K00654 |
| CBCL Emotionally Reactive K04718 | + | | |
| LEC Total K07652 | - | MODS: VicK-VicR (cell wall metabolism) two-component regulatory system | |
| LEC Turmoil K07652 | - | | |
| CBQ Impulsivity K05593 | + | KO: aadK; aminoglycoside 6-adenylyltransferase [EC:2.7.7.-] :: PFAMS: Streptomycin adenylyltransferase; PrkA serine protein kinase C-terminal domain; Nucleotidyltransferase domain; Protein of unknown function (DUF632); Immunity protein 51 | |
| CBQ Inhibitory Control K01991 | + | KO: ABC-2.OM, wza; polysaccharide biosynthesis/export protein :: PFAMS: Polysaccharide biosynthesis/export protein; SLBB domain; Late transcription unit A protein; Domain of unknown function (DUF5122) beta-propeller; TrkA-C domain | |
| CBQ Inhibitory Control K01993 | + | KO: ABC-2.TX; HlyD family secretion protein :: PFAMS: HlyD family secretion protein; Barrel-sandwich domain of CusB or HlyD membrane-fusion; Biotin-lipoyl like; Biotin-requiring enzyme; HlyD membrane-fusion protein of T1SS | K09964 K05570 |
| C Unsafe/Violent Neighborhood K00709 | - | KO: ABO; histo-blood group ABO system transferase [EC:2.4.1.40 2.4.1.37] :: PFAMS: Glycosyltransferase family 6; Retro-transcribing viruses envelope glycoprotein; Retroviral envelope protein; Domain of unknown function (DUF4898) | |
| CBQ Anger Frustration K00997 | + | KO: acpS; holo-[acyl-carrier protein] synthase [EC:2.7.8.7] :: PFAMS: 4'-phosphopantetheinyl transferase superfamily; Tectiviridae; minor capsid; Putative antitoxin of bacterial toxin-antitoxin system; YdaS/YdaT; Fimbrial; major and minor subunit; Uncharacterized protein family UPF0054 |  |
| CBQ Impulsivity K13530 | - | KO: adaA; AraC family transcriptional regulator, regulatory protein of adaptative response / methylphosphotriester-DNA alkyltransferase methyltransferase [EC:2.1.1.-] :: PFAMS: Metal binding domain of Ada; Helix-turn-helix domain; Bacterial regulatory helix-turn-helix proteins; AraC family; UBX domain; Sigma-70; region 4 | |
| CBCL Depressive Problems K11173 | + | KO: ADHFE1; hydroxyacid-oxoacid transhydrogenase [EC:1.1.99.24] :: PFAMS: Iron-containing alcohol dehydrogenase; Periplasmic binding protein; Family of unknown function (DUF5333); RNA polymerase Rpb3/Rpb11 dimerisation domain; B-box zinc finger | |
| C Unsafe/Violent Neighborhood K10975 | - | KO: allP; allantoin permease :: PFAMS: Permease for cytosine/purines; uracil; thiamine; allantoin; Polycystin cation channel; Spore morphogenesis and germination protein YwcE; ESSS subunit of NADH:ubiquinone oxidoreductase (complex I); IPP transferase | K10820 K10975 K10974 K07243 K14708 K14701 K10973 |
| CBCL Depressive Problems K10973 | + | KO: allR; IclR family transcriptional regulator, negative regulator of allantoin and glyoxylate utilization operons :: PFAMS: Bacterial transcriptional regulator; IclR helix-turn-helix domain; MarR family; Bacterial regulatory protein; arsR family; Winged helix-turn-helix DNA-binding | K10975 K07721 |
| CBQ Anger Frustration K01490 | - | KO: AMPD; AMP deaminase [EC:3.5.4.6] :: PFAMS: Adenosine/AMP deaminase; Amidohydrolase family; Ras family; AAA domain; B-box zinc finger |  |
| CBCL Anxiety Problems K12490 | + | KO: ARAP3; Arf-GAP with Rho-GAP domain, ANK repeat and PH domain-containing protein 3 :: PFAMS: PH domain; RhoGAP domain; Pleckstrin homology domain; Putative GTPase activating protein for Arf; SAM domain (Sterile alpha motif) |  |
| LEC Family Illness/Injury K03487 | + | KO: ascG; LacI family transcriptional regulator, asc operon repressor :: PFAMS: Periplasmic binding protein-like domain; Periplasmic binding proteins and sugar binding domain of LacI family; Bacterial regulatory proteins; lacI family; Periplasmic binding protein domain; Helix-turn-helix domain | |
| amily Separation/Social Services K08343 | + | KO: ATG3; ubiquitin-like-conjugating enzyme ATG3 :: PFAMS: Autophagocytosis associated protein (Atg3); N-terminal domain; Autophagocytosis associated protein; active-site domain; Autophagocytosis associated protein C-terminal; SDA1; NUP50 (Nucleoporin 50 kDa) | |
| amily Separation/Social Services K13777 | + | KO: atuF; geranyl-CoA carboxylase alpha subunit [EC:6.4.1.5] :: PFAMS: Carbamoyl-phosphate synthase L chain; ATP binding domain; Biotin carboxylase; N-terminal domain; Biotin carboxylase C-terminal domain; Biotin-requiring enzyme; ATP-grasp domain | |
| CBQ Fear K06186 | + | KO: bamE, smpA; outer membrane protein assembly factor BamE :: PFAMS: SmpA / OmlA family; Protein of unknown function (DUF3192); Sulfolobus plasmid regulatory protein; Flagellar basal body protein FlaE; PAS domain | |
| amily Separation/Social Services K08298 | + | KO: caiB; L-carnitine CoA-transferase [EC:2.8.3.21] :: PFAMS: CoA-transferase family III; C-terminal domain of CHU protein family; Short C-terminal domain; Domain of unknown function (DUF4339); Dipeptidyl peptidase IV (DPP IV) N-terminal region | |
| CBQ Anger Frustration K01255 | + | KO: CARP, pepA; leucyl aminopeptidase [EC:3.4.11.1] :: PFAMS: Cytosol aminopeptidase family; catalytic domain; Cytosol aminopeptidase family; N-terminal domain; DALR anticodon binding domain; Aminopeptidase I zinc metalloprotease (M18); The Golgi pH Regulator (GPHR) Family N-terminal |  |
| LEC Turmoil K00816 | + | KO: CCBL; kynurenine---oxoglutarate transaminase / cysteine-S-conjugate beta-lyase / glutamine---phenylpyruvate transaminase [EC:2.6.1.7 4.4.1.13 2.6.1.64] :: PFAMS: Aminotransferase class I and II; Cys/Met metabolism PLP-dependent enzyme; DegT/DnrJ/EryC1/StrS aminotransferase family; Aminotransferase class-V; Beta-eliminating lyase | |
| C Unsafe/Violent Neighborhood K10639 | + | KO: CCNB1IP1, HEI10; E3 ubiquitin-protein ligase CCNP1IP1 [EC:2.3.2.27] :: PFAMS: Zinc finger; C3HC4 type (RING finger); zinc-RING finger domain; Prokaryotic RING finger family 4; Ring finger domain; IncA protein | K15712 K10661 K15710 |
| CBQ Anger Frustration K10049 | - | KO: CEBPG; CCAAT/enhancer binding protein (C/EBP), gamma :: PFAMS: Basic region leucine zipper; bZIP transcription factor; IncA protein; Haemolysin XhlA; bZIP Maf transcription factor |  |
| LEC Family Illness/Injury K16458 | + | KO: CEP104; centrosomal protein CEP104 :: PFAMS: UvrB/uvrC motif; Tetratricopeptide repeat; Drought induced 19 protein (Di19); zinc-binding; CLASP N terminal; C2H2-type zinc finger | K16457 |
| CBQ Impulsivity K16457 | + | KO: CEP76; centrosomal protein CEP76 :: PFAMS: CEP76 C2 domain; Transglutaminase-like superfamily; C2 domain; CC2D2A N-terminal C2 domain; Transglutaminase-like domain | K16458 |
| CBQ Fear K10750 | - | KO: CHAF1A; chromatin assembly factor 1 subunit A :: PFAMS: CAF1 complex subunit p150; region binding to CAF1-p60 at C-term; CAF1 complex subunit p150; region binding to PCNA; Chromatin assembly factor 1 complex p150 subunit; N-terminal; Chromatin assembly factor 1 subunit A; SprA-related family | |
| CBCL Depressive Problems K07227 | + | KO: chuX; heme iron utilization protein :: PFAMS: Haem utilisation ChuX/HutX; Haemin-degrading HemS.ChuX domain; LysM domain; Protein of unknown function (DUF1481); Transcriptional regulator DELLA protein N terminal | |
| CBCL Aggressive Behavior K13688 | + | KO: chvB, cgs, ndvB; cyclic beta-1,2-glucan synthetase [EC:2.4.1.-] :: PFAMS: Glycosyltransferase family 36; Glycosyl hydrolase 36 superfamily; catalytic domain; Putative glucoamylase; Protein of unknown function (DUF3131); Glucodextranase; domain N | |
| CBQ Impulsivity K14192 | + | KO: clfB; clumping factor B :: PFAMS: C-terminus of bacterial fibrinogen-binding adhesin; YSIRK type signal peptide; LPXTG cell wall anchor motif; TonB N-terminal region; Domain of unknown function (DUF1970) | K14201 |

| | | | |
|---|---|---|---|
| CBQ Anger Frustration K09882 | + | KO: cobS; cobaltochelatase CobS [EC:6.6.1.2] :: PFAMS: AAA domain (dynein-related subfamily); AAA domain; Cobaltochelatase CobS subunit N terminal; ATPase family associated with various cellular activities (AAA); CbbQ/NirQ/NorQ C-terminal | |
| CBQ Inhibitory Control K10974 | - | KO: codB; cytosine permease :: PFAMS: Permease for cytosine/purines; uracil; thiamine; allantoin; Family of unknown function (DUF5353); Endoplasmic reticulum-based factor for assembly of V-ATPase; Exopolysaccharide production repressor; DMRTA motif | K10820 K10975 K02526 K07243 K14708 K14701 |
| LEC Family Illness/Injury K16630 | + | KO: COL22A; collagen, type XXII, alpha :: PFAMS: Collagen triple helix repeat (20 copies); von Willebrand factor type A domain; Concanavalin A-like lectin/glucanases superfamily; Laminin G domain; Clostridium neurotoxin; N-terminal receptor binding | |
| LEC Poverty Related Events K16844 | + | KO: comC; (2R)-3-sulfolactate dehydrogenase (NADP+) [EC:1.1.1.338] :: PFAMS: Malate/L-lactate dehydrogenase; Domain of unknown function (DUF1731); Family of unknown function (DUF5462); Thiamine biosynthesis protein (ThiI); Domain of unknown function (DUF4774) | |
| LEC Poverty Related Events K13023 | - | KO: CPN2; carboxypeptidase N regulatory subunit :: PFAMS: Leucine rich repeat; Leucine rich repeats (6 copies); Leucine Rich repeats (2 copies); Leucine-rich repeat; Leucine Rich Repeat | |
| C Unsafe/Violent Neighborhood K10914 | - | KO: crp; CRP/FNR family transcriptional regulator, cyclic AMP receptor protein :: PFAMS: Cyclic nucleotide-binding domain; Bacterial regulatory proteins; crp family; Crp-like helix-turn-helix domain; MarR family; Helix-turn-helix domain | K16326 |
| LEC Family Illness/Injury K09879 | + | KO: crtU; isorenieratene synthase :: PFAMS: Flavin containing amine oxidoreductase; Pyridine nucleotide-disulphide oxidoreductase; FAD dependent oxidoreductase; NAD(P)-binding Rossmann-like domain; FAD binding domain | |
| CBCL Depressive Problems K04333 | + | KO: csgD; LuxR family transcriptional regulator, csgAB operon transcriptional regulatory protein :: PFAMS: Bacterial regulatory proteins; luxR family; Sigma-70; region 4; Winged helix-turn-helix DNA-binding; Homeodomain-like domain; Helix-turn-helix domain | |
| CBQ Inhibitory Control K14407 | + | KO: CSTF2, RNA15; cleavage stimulation factor subunit 2 :: PFAMS: Hinge domain of cleavage stimulation factor subunit 2; RNA recognition motif. (a.k.a. RRM; RBD; or RNP domain); Transcription termination and cleavage factor C-terminal; RNA recognition motif; Occluded RNA-recognition motif | |
| CBCL Depressive Problems K12952 | + | KO: ctpE; cation-transporting P-type ATPase E [EC:3.6.3.-] :: PFAMS: E1-E2 ATPase; haloacid dehalogenase-like hydrolase; Cation transporting ATPase; C-terminus; Cation transporter/ATPase; N-terminus; Protein of unknown function (DUF454) | K12954 K01560 K05967 K05315 |
| CBCL Depressive Problems K12954 | + | KO: ctpG; cation-transporting P-type ATPase G [EC:3.6.3.-] :: PFAMS: haloacid dehalogenase-like hydrolase; E1-E2 ATPase; Haloacid dehalogenase-like hydrolase; Uncharacterised conserved protein (DUF2156) | K12952 K01560 K05967 K05315 |
| CBQ Inhibitory Control K15726 | + | KO: czcA; cobalt-zinc-cadmium resistance protein CzcA :: PFAMS: AcrB/AcrD/AcrF family; MMPL family; Outer membrane efflux protein; Protein export membrane protein; Dimerisation domain of Zinc Transporter | K16846 K14716 K02074 K15727 |
| CBQ Inhibitory Control K15727 | + | KO: czcB; membrane fusion protein, cobalt-zinc-cadmium efflux system :: PFAMS: Barrel-sandwich domain of CusB or HlyD membrane-fusion; HlyD family secretion protein; Biotin-lipoyl like; Biotin-requiring enzyme; HlyD membrane-fusion protein of T1SS | K16846 K14716 K02074 K15726 |
| CBCL Depressive Problems K13181 | + | KO: DDX27, DRS1; ATP-dependent RNA helicase DDX27 [EC:3.6.4.13] :: PFAMS: DEAD/DEAH box helicase; Helicase conserved C-terminal domain; Type III restriction enzyme; res subunit; AAA domain; Utp25; U3 small nucleolar RNA-associated SSU processome protein 25 | |
| LEC Poverty Related Events K05988 | - | KO: dexA; dextranase [EC:3.2.1.11] :: PFAMS: Glycosyl hydrolase family 66; LPXTG cell wall anchor motif; MG2 domain; Protein of unknown function (DUF3962) | K01224 K01220 K06113 K09963 K09955 K01187 K05991 K01223 K11931 K15531 K15924 |
| CBQ Impulsivity K13677 | + | KO: dgs, bgsA; 1,2-diacylglycerol-3-alpha-glucose alpha-1,2-glucosyltransferase [EC:2.4.1.208] :: PFAMS: Glycosyl transferases group 1; Glycosyltransferase Family 4 | |
| CBQ Fear K14333 | + | KO: DHBD; 2,3-dihydroxybenzoate decarboxylase [EC:4.1.1.46] :: PFAMS: Amidohydrolase; Xylose isomerase-like TIM barrel; Protein of unknown function (DUF1778); SAICAR synthetase; TatD related DNase | |
| CBCL Depressive Problems K11592 | - | KO: DICER1, DCR1; endoribonuclease Dicer [EC:3.1.26.-] :: PFAMS: Ribonuclease III domain; Ribonuclease-III-like; Dicer dimerisation domain; PAZ domain; Helicase conserved C-terminal domain | |
| CBCL Depressive Problems K03722 | + | KO: dinG; ATP-dependent DNA helicase DinG [EC:3.6.4.12] :: PFAMS: Helicase C-terminal domain; DEAD/DEAH box helicase; Exonuclease; Type III restriction enzyme; res subunit; DEAD_2 | K10738 |
| amily Separation/Social Services K08092 | + | KO: dlgD; 3-dehydro-L-gulonate 2-dehydrogenase [EC:1.1.1.130] :: PFAMS: Malate/L-lactate dehydrogenase; Dermonecrotoxin of the Papain-like fold; C3HC zinc finger-like; Flagellar C1a complex subunit C1a-32; vWA found in TerF C terminus | |
| CBQ Anger Frustration K07307 | + | KO: dmsB; anaerobic dimethyl sulfoxide reductase subunit B (DMSO reductase iron- sulfur subunit) :: PFAMS: 4Fe-4S dicluster domain; 4Fe-4S binding domain; 4Fe-4S double cluster binding domain; 4Fe-4S single cluster domain; Electron transfer flavoprotein-ubiquinone oxidoreductase; 4Fe-4S | |
| CBQ Inhibitory Control K03686 | - | KO: dnaJ; molecular chaperone DnaJ :: PFAMS: DnaJ C terminal domain; DnaJ domain; DnaJ central domain; Tryptophan RNA-binding attenuator protein inhibitory protein; Cytochrome c7 and related cytochrome c | |
| amily Separation/Social Services K12204 | + | KO: dotC, traI; defect in organelle trafficking protein DotC :: PFAMS: Type IV secretory system; conjugal DNA-protein transfer; LysM domain; Pkip-1 protein; Domain of unknown function (DUF4734); Sterol carrier protein domain | |
| LEC Total K11790 | + | KO: DTL, CDT2, DCAF2; denticleless :: PFAMS: WD domain; G-beta repeat; Anaphase-promoting complex subunit 4 WD40 domain; WD40 region of Ge1; enhancer of mRNA-decapping protein; Nucleoporin Nup120/160; Eukaryotic translation initiation factor eIF2A | |
| CBQ Inhibitory Control K06058 | + | KO: DTX; deltex [EC:2.3.2.27] :: PFAMS: WWE domain; Zinc finger; C3HC4 type (RING finger); Ring finger domain; RING-type zinc-finger; RING-H2 zinc finger domain | |
| CBQ Inhibitory Control K05540 | - | KO: dusB; tRNA-dihydrouridine synthase B [EC:1.-.-.-] :: PFAMS: Dihydrouridine synthase (Dus); Histidine biosynthesis protein; Putative N-acetylmannosamine-6-phosphate epimerase; Dihydroorotate dehydrogenase; SOR/SNZ family | |
| CBCL Anxious Depressed K09461 | + | KO: E1.14.13.40; anthraniloyl-CoA monooxygenase [EC:1.14.13.40] :: PFAMS: NADH:flavin oxidoreductase / NADH oxidase family; FAD binding domain; Pyridine nucleotide-disulphide oxidoreductase; Lycopene cyclase protein; NAD(P)-binding Rossmann-like domain | |
| CBCL Depressive Problems K00256 | + | KO: E1.3.99.16; isoquinoline 1-oxidoreductase [EC:1.3.99.16] :: PFAMS: Molybdopterin-binding domain of aldehyde dehydrogenase; [2Fe-2S] binding domain; 2Fe-2S iron-sulfur cluster binding domain; Aldehyde oxidase and xanthine dehydrogenase; a/b hammerhead domain; Cytochrome C oxidase; cbb3-type; subunit III | K02258 K07112 |
| CBQ Anger Frustration K00900 | - | KO: E2.7.1.105, PFK; 6-phosphofructo-2-kinase [EC:2.7.1.105] :: PFAMS: 6-phosphofructo-2-kinase; Histidine phosphatase superfamily (branch 1); AAA domain; Chromatin associated protein KTI12; Adenylylsulphate kinase | K16328 |
| CBQ Anger Frustration K00851 | + | KO: E2.7.1.12, gntK, idnK; gluconokinase [EC:2.7.1.12] :: PFAMS: AAA domain; FGGY family of carbohydrate kinases; N-terminal domain; FGGY family of carbohydrate kinases; C-terminal domain; Shikimate kinase; Adenylylsulphate kinase | |
| CBQ Inhibitory Control K05991 | - | KO: E3.2.1.123; endoglycosylceramidase [EC:3.2.1.123] :: PFAMS: Cellulase (glycosyl hydrolase family 5); Beta-galactosidase; Glycosyl hydrolases family 35; Defensin propeptide | K11903 K11906 K11910 K01224 K01220 K01058 |
| amily Separation/Social Services K01223 | + | KO: E3.2.1.86B, bglA; 6-phospho-beta-glucosidase [EC:3.2.1.86] :: PFAMS: Glycosyl hydrolase family 1; Cellulase (glycosyl hydrolase family 5); Glycosyl hydrolase family 10; Protein of unknown function (DUF4038); Albumin I chain a | K01224 K01220 K06113 K09963 K09955 K05988 K05991 K01187 K11931 K15531 K15924 |
| CBCL Depressive Problems K01426 | + | KO: E3.5.1.4, amiE; amidase [EC:3.5.1.4] :: PFAMS: Amidase; Carbon-nitrogen hydrolase; Acetamidase/Formamidase family; Ribosomal protein S7p/S5e; NUDIX; or N-terminal NPxY motif-rich; region of KRIT | K01455 |
| CBCL Depressive Problems K01455 | + | KO: E3.5.1.49; formamidase [EC:3.5.1.49] :: PFAMS: Acetamidase/Formamidase family; Carbon-nitrogen hydrolase; Amidohydrolase; zinc-ribbon domain; Putative PD-(D/E)XK phosphodiesterase (DUF2161) | K01426 |
| CBQ Impulsivity K01505 | - | KO: E3.5.99.7; 1-aminocyclopropane-1-carboxylate deaminase [EC:3.5.99.7] :: PFAMS: Pyridoxal-phosphate dependent enzyme; Sin-like protein conserved region; LytB protein; Variant SH3 domain; Transglutaminase-like superfamily | |
| CBQ Anger Frustration K01560 | + | KO: E3.8.1.2; 2-haloacid dehalogenase [EC:3.8.1.2] :: PFAMS: Haloacid dehalogenase-like hydrolase; haloacid dehalogenase-like hydrolase; HAD-hyrolase-like; Putative Phosphatase; Mitochondrial PGP phosphatase | K05967 K12952 K12954 |
| CBQ Inhibitory Control K09459 | + | KO: E4.1.1.82; phosphonopyruvate decarboxylase [EC:4.1.1.82] :: PFAMS: Thiamine pyrophosphate enzyme; N-terminal TPP binding domain; Thiamine pyrophosphate enzyme; C-terminal TPP binding domain; DinB superfamily; Fructose-1;6-bisphosphatase; AAA domain | |
| CBQ Anger Frustration K01802 | - | KO: E5.2.1.8; peptidylprolyl isomerase [EC:5.2.1.8] :: PFAMS: Cyclophilin type peptidyl-prolyl cis-trans isomerase/CLD; FKBP-type peptidyl-prolyl cis-trans isomerase; Phosphotyrosyl phosphate activator (PTPA) protein; PPIC-type PPIASE domain; Domain amino terminal to FKBP-type peptidyl-prolyl isomerase | |
| CBCL Depressive Problems K11249 | + | KO: eamB; cysteine/O-acetylserine efflux protein :: PFAMS: LysE type translocator; Cation transporter/ATPase; N-terminus; Integral membrane protein TerC family; TrbC/VIRB2 family; ATP synthase subunit H | K11939 |
| CBCL Emotionally Reactive K14087 | + | KO: echB; ech hydrogenase subunit B :: PFAMS: NADH dehydrogenase; Protein of unknown function (DUF2721); YtpI-like protein; PRA1 family protein; NADH-ubiquinone oxidoreductase MWFE subunit | K14091 |
| CBQ Fear K07393 | + | KO: ECM4, yqjG; glutathionyl-hydroquinone reductase [EC:1.8.5.7] :: PFAMS: Glutathione S-transferase; C-terminal domain; Glutathione S-transferase; N-terminal domain; Glutaredoxin 2; C terminal domain; Testis-expressed 12; SANT/Myb-like domain of DAMP1 | |
| CBQ Impulsivity K16314 | - | KO: ELM1; serine/threonine-protein kinase ELM1 [EC:2.7.11.1] :: PFAMS: Protein kinase domain; Protein tyrosine kinase; Protein of unknown function (DUF1241); Kinase-like; Haspin like kinase domain | K15504 |
| CBCL Anxiety Problems K14211 | + | KO: ELN; elastin :: PFAMS: Nucleoporin homology of Germinal-centre associated nuclear protein | |
| CBQ Inhibitory Control K16692 | + | KO: etk-wzc; tyrosine-protein kinase Etk/Wzc [EC:2.7.10.-] :: PFAMS: G-rich domain on putative tyrosine kinase; AAA domain; Chain length determinant protein; CobQ/CobB/MinD/ParA nucleotide binding domain; NUBPL iron-transfer P-loop NTPase | |
| amily Separation/Social Services K04027 | + | KO: eutM; ethanolamine utilization protein EutM :: PFAMS: BMC domain; YopH; N-terminal; Pyruvate:ferredoxin oxidoreductase core domain II | |
| CBQ Inhibitory Control K03561 | + | KO: exbB; biopolymer transport protein ExbB :: PFAMS: MotA/TolQ/ExbB proton channel family; Domain of unknown function (DUF2341); Concanavalin A-like lectin/glucanases superfamily; Lipopolysaccharide assembly protein A domain; Predicted membrane protein (DUF2207) | K03559 |

| | | | |
|---|---|---|---|
| CBQ Inhibitory Control K03559 | + | KO: exbD; biopolymer transport protein ExbD :: PFAMS: Biopolymer transport protein ExbD/TolR; Uncharacterized conserved protein (DUF2149); NETI protein; Photosystem II 10 kDa phosphoprotein; Oxaloacetate decarboxylase; gamma chain | K03561 |
| CBCL Anxious Depressed K10255 | + | KO: FAD6, desA; acyl-lipid omega-6 desaturase (Delta-12 desaturase) [EC:1.14.19.23 1.14.19.45] :: PFAMS: Fatty acid desaturase; Phosphatidylinositol N-acetylglucosaminyltransferase; Putative Actinobacterial Holin-X; holin superfamily III; Citrate transporter; Predicted membrane protein (DUF2207) | K00665 K10680 K09940 |
| CBQ Inhibitory Control K16091 | + | KO: fecA; Fe(3+) dicitrate transport protein :: PFAMS: TonB dependent receptor; TonB-dependent Receptor Plug Domain; Outer membrane protein beta-barrel family; Secretin and TonB N terminus short domain; CarboxypepD_reg-like domain | |
| CBQ Inhibitory Control K03839 | + | KO: fldA, nifF, isiB; flavodoxin I :: PFAMS: Flavodoxin; Flavodoxin domain; NADPH-dependent FMN reductase; NrdI Flavodoxin like; Holliday junction resolvase | |
| LEC Family Illness/Injury K02411 | - | KO: fliH; flagellar assembly protein FliH :: PFAMS: Flagellar assembly protein FliH; Essential protein Yae1; N terminal; Nodulation protein NolV; Vacuolar (H+)-ATPase G subunit; ATP synthase B/B' CF(0) | |
| Income to Needs K04437 | + | KO: FLNA; filamin :: PFAMS: Filamin/ABP280 repeat; Calponin homology (CH) domain; Y_Y_Y domain; Domain of unknown function (DUF5060); Bacterial Ig-like domain (group 3) | |
| CBCL Depressive Problems K07243 | + | KO: FTR, FTH1, efeU; high-affinity iron transporter :: PFAMS: Iron permease FTR1 family; Cytochrome C oxidase; cbb3-type; subunit III; Cytochrome c; Major Facilitator Superfamily; Alpha/beta-hydrolase family N-terminus | K10820 K10975 K10974 K02526 K14708 K14701 |
| CBQ Inhibitory Control K09669 | + | KO: FUT10; alpha-1,3-fucosyltransferase 10 [EC:2.4.1.-] :: PFAMS: Glycosyltransferase family 10 (fucosyltransferase) C-term; Fucosyltransferase; N-terminal; Sigma-70; region 4; Mid domain of argonaute; Tap; RNA-binding | |
| CBQ Impulsivity K16515 | + | KO: galB; 4-oxalomesaconate hydratase [EC:4.2.1.83] :: PFAMS: GlcNAc-PI de-N-acetylase; Glycosyl transferase 4-like; Putative sugar-binding domain; CBL proto-oncogene N-terminal domain 1; Helix-turn-helix domain | |
| CBQ Inhibitory Control K04618 | + | KO: GAOA; galactose oxidase [EC:1.1.3.9] :: PFAMS: Kelch motif; Domain of unknown function (DUF1929); Ricin-type beta-trefoil lectin domain-like; Galactose oxidase; central domain; Ricin-type beta-trefoil lectin domain | K01785 |
| CBCL Anxiety Problems K02436 | + | KO: gatR; DeoR family transcriptional regulator, galactitol utilization operon repressor :: PFAMS: DeoR C terminal sensor domain; DeoR-like helix-turn-helix domain; Bacterial regulatory proteins; gntR family; HTH domain; Coenzyme A transferase | |
| :C Unsafe/Violent Neighborhood K13211 | + | KO: GCFC; GC-rich sequence DNA-binding factor :: PFAMS: GC-rich sequence DNA-binding factor-like protein; Nineteen complex-related protein 2; Lipoprotein amino terminal region; Growth-arrest specific micro-tubule binding; Protein of unknown function (DUF1687) | |
| CBCL Depressive Problems K03566 | + | KO: gcvA; LysR family transcriptional regulator, glycine cleavage system transcriptional activator :: PFAMS: LysR substrate binding domain; Bacterial regulatory helix-turn-helix protein; lysR family; PucR C-terminal helix-turn-helix domain; MarR family; Helix-turn-helix domain | |
| CBQ Inhibitory Control K06296 | - | KO: gerKB; spore germination protein KB :: PFAMS: Spore germination protein; Sodium:alanine symporter family; Malarial early transcribed membrane protein (ETRAMP); Spore germination B3/ GerAC like; C-terminal; Protein of unknown function (DUF2530) | K06294 |
| CBQ Anger Frustration K12972 | + | KO: ghrA; glyoxylate/hydroxypyruvatereductase [EC:1.1.1.79 1.1.1.81] :: PFAMS: D-isomer specific 2-hydroxyacid dehydrogenase; NAD binding domain; NAD binding domain of 6-phosphogluconate dehydrogenase; NAD-dependent glycerol-3-phosphate dehydrogenase N-terminus; 3-hydroxyacyl-CoA dehydrogenase; NAD binding domain; D-isomer specific 2-hydroxyacid dehydrogenase; catalytic domain | |
| CBQ Inhibitory Control K03501 | - | KO: gidB, rsmG; 16S rRNA (guanine527-N7)-methyltransferase [EC:2.1.1.170] :: PFAMS: rRNA small subunit methyltransferase G; Methyltransferase domain; Methyltransferase small domain; Lysine methyltransferase; Putative methyltransferase | |
| CBQ Impulsivity K00111 | + | KO: glpA, glpD; glycerol-3-phosphate dehydrogenase [EC:1.1.5.3] :: PFAMS: FAD dependent oxidoreductase; C-terminal domain of alpha-glycerophosphate oxidase; FAD binding domain; Pyridine nucleotide-disulphide oxidoreductase; NAD(P)-binding Rossmann-like domain | |
| CBQ Anger Frustration K10670 | + | KO: grdA; glycine/sarcosine/betaine reductase complex component A [EC:1.21.4.2 1.21.4.3 1.21.4.4] :: PFAMS: Glycine/sarcosine/betaine reductase selenoprotein B (GRDB); Glycine/sarcosine/betaine reductase component B subunits; Fatty acid synthesis protein; Epidermal patterning factor proteins; Tex protein YqgF-like domain | K10793 K10794 K10795 |
| CBCL Depressive Problems K01706 | + | KO: gudD; glucarate dehydratase [EC:4.2.1.40] :: PFAMS: Enolase C-terminal domain-like; Mandelate racemase / muconate lactonizing enzyme; N-terminal domain; Radical SAM superfamily; Acyl-CoA dehydrogenase; C-terminal domain; Quinolinate phosphoribosyl transferase; C-terminal domain | |
| CBCL Anxious Depressed K00015 | + | KO: gyaR, GOR1; glyoxylate reductase [EC:1.1.1.26] :: PFAMS: D-isomer specific 2-hydroxyacid dehydrogenase; NAD binding domain; D-isomer specific 2-hydroxyacid dehydrogenase; catalytic domain; NAD binding domain of 6-phosphogluconate dehydrogenase; NADP oxidoreductase coenzyme F420-dependent; Acetohydroxy acid isomeroreductase; NADPH-binding domain | |
| Income to Needs K16589 | - | KO: HAUS6; HAUS augmin-like complex subunit 6 :: PFAMS: HAUS augmin-like complex subunit 6 N-terminus; Pilus assembly protein; PilO; Bacterial flagellin N-terminal helical region; Protein SOGA; Cell division protein ZapB | K14744 K03746 |
| CBQ Fear K03580 | + | KO: hepA; ATP-dependent helicase HepA [EC:3.6.4.-] :: PFAMS: RNA polymerase recycling family C-terminal; SNF2 family N-terminal domain; Helicase conserved C-terminal domain; Type III restriction enzyme; res subunit; DEAD/DEAH box helicase | |
| CBCL Depressive Problems K07225 | + | KO: hmuS; putative hemin transport protein :: PFAMS: Haemin-degrading HemS.ChuX domain; Haem utilisation ChuX/HutX; Protein of unknown function (DUF3920); PcrB family; Family of unknown function (DUF5473) | |
| CBQ Inhibitory Control K03705 | - | KO: hrcA; heat-inducible transcriptional repressor :: PFAMS: HrcA protein C terminal domain; Winged helix-turn-helix transcription repressor; HrcA DNA-binding; DeoR-like helix-turn-helix domain; HTH domain; LexA DNA binding domain | |
| CBQ Impulsivity K08784 | + | KO: HTRA1, PRSS11; HtrA serine peptidase 1 [EC:3.4.21.-] :: PFAMS: Trypsin-like peptidase domain; Trypsin; Kazal-type serine protease inhibitor domain; PDZ domain (Also known as DHR or GLGF); PDZ domain | |
| CBCL Depressive Problems K06281 | + | KO: hyaB, hybC; hydrogenase large subunit [EC:1.12.99.6] :: PFAMS: Nickel-dependent hydrogenase; Glycosyl hydrolase 108; Berberine and berberine like; Foot-and-mouth virus L-proteinase; Serine-threonine protein kinase 19 | |
| Income to Needs K12136 | + | KO: hyfA; hydrogenase-4 component A [EC:1.-.-.-] :: PFAMS: 4Fe-4S dicluster domain; 4Fe-4S binding domain; 4Fe-4S double cluster binding domain; 4Fe-4S single cluster domain; 4Fe-4S single cluster domain of Ferredoxin I | K12137 K12142 |
| CBQ Anger Frustration K12137 | + | KO: hyfB; hydrogenase-4 component B [EC:1.-.-.-] :: PFAMS: Proton-conducting membrane transporter; NADH-Ubiquinone oxidoreductase (complex I); chain 5 N-terminus; Protein of unknown function (DUF4225); 7TM diverse intracellular signalling; YesK-like protein | K12136 K12142 |
| Income to Needs K12142 | + | KO: hyfG; hydrogenase-4 component G [EC:1.-.-.-] :: PFAMS: Respiratory-chain NADH dehydrogenase; 49 Kd subunit; Respiratory-chain NADH dehydrogenase; 30 Kd subunit; Nickel-dependent hydrogenase | K12136 K12137 |
| CBCL Depressive Problems K04651 | + | KO: hypA, hybF; hydrogenase nickel incorporation protein HypA/HybF :: PFAMS: Hydrogenase/urease nickel incorporation; metallochaperone; hypA; Zinc ribbon domain; zinc-ribbon; zinc-ribbon domain; Double zinc ribbon | |
| CBQ Anger Frustration K01473 | + | KO: hyuA; N-methylhydantoinase A [EC:3.5.2.14] :: PFAMS: Hydantoinase/oxoprolinase; Hydantoinase/oxoprolinase N-terminal region; MutL protein; BadF/BadG/BcrA/BcrD ATPase family; Cell division protein FtsA | |
| amily Separation/Social Services K05596 | + | KO: iciA; LysR family transcriptional regulator, chromosome initiation inhibitor :: PFAMS: Bacterial regulatory helix-turn-helix protein; lysR family; LysR substrate binding domain; PucR C-terminal helix-turn-helix domain; MarR family; Helix-turn-helix domain | |
| CBCL Depressive Problems K13641 | + | KO: iclR; IclR family transcriptional regulator, acetate operon repressor :: PFAMS: Bacterial transcriptional regulator; IclR helix-turn-helix domain; MarR family; Winged helix-turn-helix DNA-binding; Transcriptional regulator | |
| amily Separation/Social Services K12209 | + | KO: icmE, dotG; intracellular multiplication protein IcmE :: PFAMS: Pentapeptide repeats (9 copies); Bacterial conjugation TrbI-like protein; Bacterial toxin 33; Ribosomal protein L13e; Domain of unknown function (DUF1127) | K12217 K12218 |
| amily Separation/Social Services K12217 | + | KO: icmO, trbC, dotL; intracellular multiplication protein IcmO :: PFAMS: Type IV secretion-system coupling protein DNA-binding domain; TraM recognition site of TraD and TraG; Domain of unknown function DUF87; FtsK/SpoIIIE family; Type IV secretory system Conjugative DNA transfer | K12209 K12218 |
| amily Separation/Social Services K12218 | + | KO: icmP, trbA; intracellular multiplication protein IcmP :: PFAMS: Uncharacterized protein conserved in bacteria (DUF2330); Family of unknown function; Protein of unknown function (DUF1235); Phosphatidylglycerophosphatase A; Protein of unknown function (DUF4079) | K12209 K12217 |
| LEC Poverty Related Events K09826 | + | KO: irr; Fur family transcriptional regulator, iron response regulator :: PFAMS: Ferric uptake regulator family; Transposase; Helix-turn-helix domain; Sugar-specific transcriptional regulator TrmB; Tetratricopeptide repeat | |
| Income to Needs K01063 | + | KO: JHE; juvenile-hormone esterase [EC:3.1.1.59] :: PFAMS: Carboxylesterase family; alpha/beta hydrolase fold; Prolyl oligopeptidase family; Putative esterase; Sif protein | |
| LEC Family Illness/Injury K00183 | + | KO: K00183; prokaryotic molybdopterin-containing oxidoreductase family, molybdopterin binding subunit :: PFAMS: Molybdopterin oxidoreductase; Molydopterin dinucleotide binding domain; Molybdopterin oxidoreductase Fe4S4 domain; TAT (twin-arginine translocation) pathway signal sequence; Domain of unknown function (DUF4440) | K08357 K00397 |
| CBCL Depressive Problems K05967 | - | KO: K05967; uncharacterized protein :: PFAMS: 5' nucleotidase; deoxy (Pyrimidine); cytosolic type C protein (NT5C); HAD superfamily; subfamily IIIB (Acid phosphatase); Haloacid dehalogenase-like hydrolase; 5'-nucleotidase; haloacid dehalogenase-like hydrolase | K12954 K01560 K12952 |
| CBQ Anger Frustration K06910 | + | KO: K06910; uncharacterized protein :: PFAMS: Phosphatidylethanolamine-binding protein; Polyketide cyclase / dehydrase and lipid transport; Clustered mitochondria; Domain of unknown function (DUF4292); Protein of unknown function (DUF998) | |
| CBCL Depressive Problems K06922 | - | KO: K06922; uncharacterized protein :: PFAMS: Protein of unknown function (DUF499); Protein of unknown function (DUF3619); Minor tail protein T; Domain of unknown function (DUF4154); TATA-box binding | |
| CBQ Inhibitory Control K07033 | - | KO: K07033; uncharacterized protein :: PFAMS: Uncharacterized protein family (UPF0051); Uncharacterised protein family (UPF0181); Geminivirus rep protein central domain; T4 recombination endonuclease VII; dimerisation; Beta/Gamma crystallin | |
| CBQ Anger Frustration K07038 | + | KO: K07038; inner membrane protein :: PFAMS: LexA-binding; inner membrane-associated putative hydrolase; Uncharacterized metal-binding protein (DUF2227); Zinc dependent phospholipase C; Domain of unknown function (DUF4184); Protein of unknown function (DUF3043) | |
| LEC Family Illness/Injury K07059 | - | KO: K07059; uncharacterized protein :: PFAMS: Rhomboid family; AN1-like Zinc finger; Eukaryotic integral membrane protein (DUF1751); Ubiquitin family; Tc5 transposase C-terminal domain | |
| CBQ Inhibitory Control K07098 | + | KO: K07098; uncharacterized protein :: PFAMS: Calcineurin-like phosphoesterase; Calcineurin-like phosphoesterase superfamily domain; Metallophosphoesterase; calcineurin superfamily; TAT (twin-arginine translocation) pathway signal sequence; Cytochrome B6-F complex Fe-S subunit | |

| | | | |
|---|---|---|---|
| CBQ Anger Frustration K07109 | + | KO: K07109; uncharacterized protein :: PFAMS: Bacterial protein of unknown function (YtfJ_HI0045); ATP10 protein; Domain of unknown function (DUF4174); Pyridoxamine 5'-phosphate oxidase; Protein of unknown function (DUF1007) | |
| CBQ Anger Frustration K07112 | + | KO: K07112; uncharacterized protein :: PFAMS: Sulphur transport; Domain of unknown function (DUF4073); Sulfurtransferase TusA; Molybdate transporter of MFS superfamily; Cytochrome C oxidase subunit II; transmembrane | K02258 K00256 |
| CBCL Depressive Problems K07177 | + | KO: K07177; Lon-like protease :: PFAMS: PDZ domain; Lon protease (S16) C-terminal proteolytic domain; PDZ domain (Also known as DHR or GLGF); Tricorn protease PDZ domain; Birnavirus VP4 protein | K04770 |
| CBQ Inhibitory Control K07231 | + | KO: K07231; putative iron-regulated protein :: PFAMS: Imelysin; Domain of unknown function (DUF4349); Prokaryotic lipoprotein-attachment site; Domain of unknown function (DUF5102); Potyvirus coat protein | |
| Income to Needs K07234 | + | KO: K07234; uncharacterized protein involved in response to NO :: PFAMS: NnrS protein; Tetratricopeptide repeat; Family of unknown function (DUF5316); Bacterial membrane protein YfhO; Conserved TM helix | |
| LEC Poverty Related Events K07492 | + | KO: K07492; putative transposase :: PFAMS: Putative transposase of IS4/5 family (DUF4096); Transposase DDE domain; DDE superfamily endonuclease; Protein of unknown function (DUF1048); ATP phosphoribosyltransferase | |
| CBCL Depressive Problems K07498 | + | KO: K07498; putative transposase :: PFAMS: DDE domain; Integrase core domain; Transposase IS66 family; Homeodomain-like domain; DDE superfamily endonuclease | |
| CBQ Anger Frustration K07721 | + | KO: K07721; ArsR family transcriptional regulator :: PFAMS: Helix-turn-helix domain; Bacterial regulatory protein; arsR family; MarR family; Sugar-specific transcriptional regulator TrmB; Winged helix-turn-helix DNA-binding | K10973 |
| CBQ Anger Frustration K09137 | + | KO: K09137; uncharacterized protein :: PFAMS: Uncharacterized ACR; COG1993; Small subunit of acetolactate synthase; Phosphoenolpyruvate hydrolase-like; CBS domain; Region in Clathrin and VPS | |
| CBQ Anger Frustration K09142 | + | KO: K09142; uncharacterized protein :: PFAMS: Putative RNA methyltransferase; SpoU rRNA Methylase family; Translation initiation factor 1A / IF-1; Amyotrophic lateral sclerosis 2 chromosomal region candidate gene 8; AF2226-like SPOUT RNA Methylase fused to THUMP | |
| CBQ Impulsivity K09703 | + | KO: K09703; uncharacterized protein :: PFAMS: Protein of unknown function (DUF917); Protein of unknown function (DUF1296); TrfA protein; Uncharacterised 5xTM membrane BCR; YitT family COG1284; Domain of unknown function (DUF4774) | |
| CBCL Emotionally Reactive K09764 | + | KO: K09764; uncharacterized protein :: PFAMS: Protein of unknown function (DUF503); Type II secretion system pilotin lipoprotein (PulS_OutS); Protein of unknown function (DUF1351); Family of unknown function (DUF5468); Diguanylate cyclase; GGDEF domain | K02454 K02663 K06218 |
| CBQ Fear K09896 | + | KO: K09896; uncharacterized protein :: PFAMS: Protein of unknown function (DUF1040); Orthopoxvirus F14 protein; NADPH-dependent FMN reductase; P53 tetramerisation motif; Vacuolar sorting protein 39 domain 1 | |
| LEC Family Illness/Injury K09931 | + | KO: K09931; uncharacterized protein :: PFAMS: Uncharacterized protein conserved in bacteria (DUF2064); Guanylyl transferase CofC like; MobA-like NTP transferase domain; Glycosyl transferase family 2; Glycosyltransferase like family 2 | |
| CBQ Inhibitory Control K09940 | + | KO: K09940; uncharacterized protein :: PFAMS: Domain of unknown function (DUF4870); Predicted membrane protein (DUF2207); Fatty acid desaturase; Tetraspanin family; ABC-2 family transporter protein | K10255 K10680 K00665 |
| :C Unsafe/Violent Neighborhood K09964 | - | KO: K09964; uncharacterized protein :: PFAMS: ACT domain; Acetyltransferase (GNAT) domain; Acetyltransferase (GNAT) family; Putative prokaryotic signal transducing protein; Peptidase S24-like | K01993 K05570 |
| CBQ Impulsivity K09979 | + | KO: K09979; uncharacterized protein :: PFAMS: Pyridoxamine 5'-phosphate oxidase; 1-deoxy-D-xylulose-5-phosphate synthase; Pyridoxamine 5'-phosphate oxidase like; Natural killer cell receptor 2B4; F420H(2)-dependent quinone reductase | |
| CBCL Aggressive Behavior K11442 | + | KO: K11442; putative uridylyltransferase [EC:2.7.7.-] :: PFAMS: UTP--glucose-1-phosphate uridylyltransferase; MobA-like NTP transferase domain; Domain of unknown function (DUF4301); PPAR gamma N-terminal region; Nucleotidyl transferase | |
| Income to Needs K15640 | + | KO: K15640, phoE; uncharacterized phosphatase :: PFAMS: Histidine phosphatase superfamily (branch 1); Serine hydrolase; GDSL-like Lipase/Acylhydrolase family; Terminase RNaseH-like domain; Lipase (class 3) | |
| LEC Family Illness/Injury K11211 | - | KO: kdkA; 3-deoxy-D-manno-octulosonic acid kinase [EC:2.7.1.166] :: PFAMS: Lipopolysaccharide kinase (Kdo/WaaP) family; RIO1 family; Phosphotransferase enzyme family; Protein kinase domain; Choline/ethanolamine kinase | |
| CBQ Inhibitory Control K05636 | + | KO: LAMB1; laminin, beta 1 :: PFAMS: Laminin EGF domain; Laminin N-terminal (Domain VI); F5/8 type C domain; CorA-like Mg2+ transporter protein; Domain of unknown function (DUF2935) | |
| CBQ Anger Frustration K16120 | + | KO: licB; lichenysin synthetase B :: PFAMS: Condensation domain; AMP-binding enzyme; Phosphopantetheine attachment site; AMP-binding enzyme C-terminal domain; Transferase family | K15665 K15658 K15660 K16130 K12239 K05914 K16129 K16416 K16124 |
| CBQ Anger Frustration K03820 | + | KO: lnt; apolipoprotein N-acyltransferase [EC:2.3.1.-] :: PFAMS: Carbon-nitrogen hydrolase; tRNA Pseudouridine synthase II; C terminal; Glycosyl transferase family 2; Vacuolar protein sorting 55; Domain of unknown function (DUF4131) | |
| CBQ Anger Frustration K04770 | + | KO: lonH; Lon-like ATP-dependent protease [EC:3.4.21.-] :: PFAMS: AAA domain; Lon protease (S16) C-terminal proteolytic domain; Family of unknown function (DUF5346); Mad3/BUB1 homology region 1; SurA N-terminal domain | K07177 |
| CBCL Depressive Problems K00880 | + | KO: lyxK; L-xylulokinase [EC:2.7.1.53] :: PFAMS: FGGY family of carbohydrate kinases; N-terminal domain; FGGY family of carbohydrate kinases; C-terminal domain; BadF/BadG/BcrA/BcrD ATPase family; Acyl carrier protein phosphodiesterase; Hydantoinase/oxoprolinase N-terminal region | |
| CBCL Anxious Depressed K06061 | + | KO: MAML; mastermind :: PFAMS: MamL-1 domain; KRI1-like family; CDC45-like protein; Otopetrin; Ryanodine Receptor TM 4-6 | |
| CBCL Anxiety Problems K02538 | + | KO: manR; activator of the mannose operon, transcriptional antiterminator :: PFAMS: PRD domain; HTH domain; Phosphoenolpyruvate-dependent sugar phosphotransferase system; EIIA 2; Mga helix-turn-helix domain; DeoR-like helix-turn-helix domain | |
| LEC Family Illness/Injury K10661 | + | KO: MARCH6, DOA10; E3 ubiquitin-protein ligase MARCH6 [EC:2.3.2.27] :: PFAMS: RING-variant domain; Ring finger domain; Anaphase-promoting complex subunit 11 RING-H2 finger; FANCL C-terminal domain; Zinc finger; C3HC4 type (RING finger) | K15712 K10639 K15710 |
| amily Separation/Social Services K04765 | + | KO: mazG; nucleoside triphosphate diphosphatase [EC:3.6.1.9] :: PFAMS: MazG nucleotide pyrophosphohydrolase domain; Phosphoribosyl-ATP pyrophosphohydrolase; MazG-like family; Oxoglutarate and iron-dependent oxygenase degradation C-term; Protein of unknown function (DUF3168) | |
| :C Unsafe/Violent Neighborhood K10738 | + | KO: MCM9; DNA helicase MCM9 [EC:3.6.4.12] :: PFAMS: MCM2/3/5 family; MCM OB domain; Magnesium chelatase; subunit ChlI; AAA domain (dynein-related subfamily); ATPase family associated with various cellular activities (AAA) | K03722 |
| CBCL Aggressive Behavior K16129 | + | KO: mcyE; microcystin synthetase protein McyE :: PFAMS: Condensation domain; AMP-binding enzyme; Beta-ketoacyl synthase; N-terminal domain; Methyltransferase domain; Acyl transferase domain | K15665 K15658 K15660 K16130 K12239 K16120 K05914 K16416 K16124 |
| LEC Poverty Related Events K13929 | + | KO: mdcA; malonate decarboxylase alpha subunit [EC:2.3.1.187] :: PFAMS: Malonate decarboxylase; alpha subunit; transporter; Acetyl-CoA hydrolase/transferase C-terminal domain; Coenzyme A transferase; NocA-like zinc-finger protein 1; Transcriptional regulator; AbiEi antitoxin | K05607 K13932 K00249 K01027 |
| Income to Needs K13932 | - | KO: mdcD; malonate decarboxylase beta subunit [EC:4.1.1.87] :: PFAMS: Carboxyl transferase domain; Malonate decarboxylase gamma subunit (MdcE); Malonate decarboxylase delta subunit (MdcD); Fructose-bisphosphate aldolase class-II; Fructose-1;6-bisphosphatase | K05607 K13929 K00249 K01027 |
| CBQ Anger Frustration K06990 | + | KO: MEMO1; MEMO1 family protein :: PFAMS: Memo-like protein; AMMECR1; Catalytic LigB subunit of aromatic ring-opening dioxygenase; 3-oxo-5-alpha-steroid 4-dehydrogenase; LeuA allosteric (dimerisation) domain | |
| CBQ Impulsivity K08364 | - | KO: merP; periplasmic mercuric ion binding protein :: PFAMS: Heavy-metal-associated domain; Sulfurtransferase TusA; Domain of unknown function (DUF1930); RNA binding motif; AcrB/AcrD/AcrF family | |
| CBCL Depressive Problems K00599 | - | KO: METTL6; methyltransferase-like protein 6 [EC:2.1.1.-] :: PFAMS: Methyltransferase domain; O-methyltransferase; tRNA (Uracil-5-)-methyltransferase; SpoU rRNA Methylase family; Conserved hypothetical protein 95 | K15471 K07442 |
| CBQ Impulsivity K09186 | + | KO: MLL1; histone-lysine N-methyltransferase MLL1 [EC:2.1.1.43] :: PFAMS: SET domain; F/Y rich C-terminus; PHD-finger; F/Y-rich N-terminus; PHD-like zinc-binding domain | K11420 K11428 |
| CBCL Depressive Problems K08166 | + | KO: mmr; MFS transporter, DHA2 family, methylenomycin A resistance protein :: PFAMS: Major Facilitator Superfamily; Transmembrane secretion effector; Sugar (and other) transporter; Protein of unknown function (DUF4227); Mannosyltransferase (PIG-M) | K08369 K08172 K08144 |
| CBQ Anger Frustration K05568 | + | KO: mnhD, mrpD; multicomponent Na+:H+ antiporter subunit D :: PFAMS: Proton-conducting membrane transporter; NADH-Ubiquinone oxidoreductase (complex I); chain 5 N-terminus; NADH dehydrogenase I; subunit N related protein; Protein of unknown function (DUF1490); GET complex subunit GET2 | K05565 K05570 |
| CBQ Anger Frustration K05570 | + | KO: mnhF, mrpF; multicomponent Na+:H+ antiporter subunit F :: PFAMS: Multiple resistance and pH regulation protein F (MrpF / PhaF); Deltamethrin resistance; Family of unknown function (DUF5392); ABC-2 family transporter protein; Sulfite exporter TauE/SafE | K05568 K05570 |
| CBCL Depressive Problems K02558 | + | KO: mpl; UDP-N-acetylmuramate: L-alanyl-gamma-D-glutamyl-meso-diaminopimelate ligase [EC:6.3.2.45] :: PFAMS: Mur ligase middle domain; Mur ligase family; catalytic domain; Mur ligase family; glutamate ligase domain; CobQ/CobB/MinD/ParA nucleotide binding domain; Pyruvate ferredoxin/flavodoxin oxidoreductase | |
| CBQ Inhibitory Control K03438 | - | KO: mraW, rsmH; 16S rRNA (cytosine1402-N4)-methyltransferase [EC:2.1.1.199] :: PFAMS: MraW methylase family; Methyltransferase domain; Methyltransferase small domain; Putative rRNA methylase; FtsJ-like methyltransferase | |
| CBQ Anger Frustration K15507 | + | KO: MRM1, PET56; 21S rRNA (GM2251-2'-O)-methyltransferase [EC:2.1.1.-] :: PFAMS: SpoU rRNA Methylase family; RNA 2'-O ribose methyltransferase substrate binding; SAM-dependent RNA methyltransferase; Protein of unknown function (DUF4065); Glutamine rich N terminal domain of histone deacetylase 4 | K04768 |
| CBQ Anger Frustration K13163 | + | KO: MSL1; male-specific lethal 1 :: PFAMS: PEHE domain; Dimerisation domain of Male-specific-Lethal 1; Septum formation initiator; tRNA synthetases class I (E and Q); catalytic domain; Herpesvirus UL6 like | |
| CBQ Anger Frustration K16416 | + | KO: mxaA; myxalamid-type nonribosomal peptide synthetase MxaA :: PFAMS: AMP-binding enzyme; Condensation domain; Male sterility protein; NAD dependent epimerase/dehydratase family; Phosphopantetheine attachment site | K15665 K15658 K15660 K16130 K12239 K16120 K16129 K05914 K16124 |
| CBQ Impulsivity K03885 | - | KO: ndh; NADH dehydrogenase [EC:1.6.99.3] :: PFAMS: Pyridine nucleotide-disulphide oxidoreductase; FAD dependent oxidoreductase; FAD-NAD(P)-binding; NAD(P)-binding Rossmann-like domain; FAD binding domain | |

| | | | |
|---|---|---|---|
| CBQ Anger Frustration K07047 | + | KO: nfdA; N-substituted formamide deformylase [EC:3.5.1.91] :: PFAMS: Amidohydrolase family; TAT (twin-arginine translocation) pathway signal sequence; TatD related DNase; Cytochrome P450; Ubiquitinol-cytochrome C reductase Fe-S subunit TAT signal | |
| CBCL Depressive Problems K11671 | + | KO: NFRKB, INO80G; nuclear factor related to kappa-B-binding protein :: PFAMS: NFRKB Winged Helix-like; Asx homology domain; PSRT; Golgi-body localisation protein domain; Iron Transport-associated domain | |
| CBQ Anger Frustration K13988 | + | KO: NUDT9; ADP-ribose pyrophosphatase [EC:3.6.1.13] :: PFAMS: NUDIX domain; RNA recognition motif. (a.k.a. RRM; RBD; or RNP domain); L27 domain; Nup53/35/40-type RNA recognition motif; Protein of unknown function (DUF1033) | |
| Income to Needs K07268 | + | KO: oapA; opacity associated protein :: PFAMS: Opacity-associated protein A LysM-like domain; Opacity-associated protein A N-terminal motif; Occlusion-derived virus envelope protein ODV-E18; tRNA methyltransferase complex GCD14 subunit N-term; Eukaryotic translation initiation factor 3 subunit 8 N-terminus | |
| CBQ Inhibitory Control K03660 | - | KO: OGG1; N-glycosylase/DNA lyase [EC:3.2.2.- 4.2.99.18] :: PFAMS: 8-oxoguanine DNA glycosylase; N-terminal domain; HhH-GPD superfamily base excision DNA repair protein; Helix-hairpin-helix motif; Helix-hairpin-helix domain; RuvA; C-terminal domain | K03631 K03515 K10839 K01249 |
| Income to Needs K12982 | + | KO: opsX; heptosyltransferase I [EC:2.4.-.-] :: PFAMS: Glycosyltransferase family 9 (heptosyltransferase); Polysaccharide pyruvyl transferase; Glycosyl transferases group 1; Diacylglycerol kinase catalytic domain; Electron transfer flavoprotein domain | |
| CBCL Aggressive Behavior K00697 | + | KO: otsA; trehalose 6-phosphate synthase [EC:2.4.1.15 2.4.1.347] :: PFAMS: Glycosyltransferase family 20; Glycosyl transferases group 1; Starch synthase catalytic domain; Major ampullate spidroin 1; spider silk protein 1; N-term; N-formylglutamate amidohydrolase | |
| CBCL Internalizing Behavior K01664 | + | KO: pabA; para-aminobenzoate synthetase component II [EC:2.6.1.85] :: PFAMS: Glutamine amidotransferase class-I; Peptidase C26; SNO glutamine amidotransferase family; CobB/CobQ-like glutamine amidotransferase domain; DJ-1/PfpI family | |
| CBCL Depressive Problems K16795 | + | KO: PAFAH1B2_3; platelet-activating factor acetylhydrolase IB subunit beta/gamma [EC:3.1.1.47] :: PFAMS: GDSL-like Lipase/Acylhydrolase family; GDSL-like Lipase/Acylhydrolase; Gelsolin repeat; Metallophosphoesterase; calcineurin superfamily; Enolase; N-terminal domain | |
| amily Separation/Social Services K12518 | + | KO: papC; outer membrane usher protein PapC :: PFAMS: Outer membrane usher protein; PapC N-terminal domain; PapC C-terminal domain; Immunity protein 10 | |
| CBQ Impulsivity K00448 | + | KO: pcaG; protocatechuate 3,4-dioxygenase, alpha subunit [EC:1.13.11.3] :: PFAMS: Dioxygenase; Carboxypeptidase regulatory-like domain; Listeria-Bacteroides repeat domain (List_Bact_rpt); Salmonella repeat of unknown function (DUF824); Protocatechuate 3;4-dioxygenase beta subunit N terminal | |
| CBQ Sadness K16499 | + | KO: PCDHD2; protocadherin delta 2 :: PFAMS: Cadherin domain; Cadherin-like; Cadherin cytoplasmic C-terminal; Peptidase propeptide and YPEB domain; TMEM154 protein family | |
| LEC Family Illness/Injury K05995 | + | KO: pepE; dipeptidase E [EC:3.4.13.21] :: PFAMS: Peptidase family S51; CobB/CobQ-like glutamine amidotransferase domain; Biotin-protein ligase; N terminal; SNO glutamine amidotransferase family; Phospholipase/Carboxylesterase | |
| CBCL Depressive Problems K11931 | + | KO: pgaB; poly-beta-1,6-N-acetyl-D-glucosamine N-deacetylase [EC:3.5.1.-] :: PFAMS: Hypothetical glycosyl hydrolase family 13; Polysaccharide deacetylase; Glycosyl hydrolase-like 10; Sigma-54 factor; Activator interacting domain (AID); Uncharacterized protein conserved in bacteria (DUF2334) | K01224 K01220 K06113 K09963 K09955 K05988 K05991 K01223 K01187 K15531 K15924 |
| LEC Turmoil K00914 | + | KO: PIK3C3, VPS34; phosphatidylinositol 3-kinase [EC:2.7.1.137] :: PFAMS: Phosphoinositide 3-kinase family; accessory domain (PIK domain); Phosphatidylinositol 3- and 4-kinase; Phosphoinositide 3-kinase C2; Domain of unknown function (DUF4135); HEAT repeats | |
| CBCL Depressive Problems K02655 | + | KO: pilE; type IV pilus assembly protein PilE :: PFAMS: Type IV minor pilin ComP; DNA uptake sequence receptor; Prokaryotic N-terminal methylation motif; Minor type IV pilin; PilX; Type IV pilin-like G and H; putative; Pilin (bacterial filament) | K02663 K02666 |
| C Unsafe/Violent Neighborhood K02663 | + | KO: pilN; type IV pilus assembly protein PilN :: PFAMS: Fimbrial assembly protein (PilN); Septum formation initiator; HemX; putative uroporphyrinogen-III C-methyltransferase; Type II secretion system (T2SS); protein M; Protein of unknown function (DUF2681) | K02655 K02666 |
| CBCL Anxious Depressed K02666 | + | KO: pilQ; type IV pilus assembly protein PilQ :: PFAMS: Bacterial type II and III secretion system protein; Bacterial type II/III secretion system short domain; Secretin and TonB N terminus short domain; AMIN domain; Domain of unknown function (DUF4974) | K02655 K02663 |
| CBCL Depressive Problems K01058 | + | KO: pldA; phospholipase A1/A2 [EC:3.1.1.32 3.1.1.4] :: PFAMS: Phospholipase A1; Type VI secretion system VasI; EvfG; VC_A0118; Tetratricopeptide repeat; MIT (microtubule interacting and transport) domain; Aida N-terminus | K01220 K11903 |
| CBQ Inhibitory Control K01440 | - | KO: PNC1; nicotinamidase [EC:3.5.1.19] :: PFAMS: Isochorismatase family; Pyridine nucleotide-disulphide oxidoreductase; Putative stress-responsive nuclear envelope protein; Mitochondrial 28S ribosomal protein S34; Domain of unknown function (DUF4362) | |
| CBQ Impulsivity K15529 | + | KO: POA1; ADP-ribose 1''-phosphate phosphatase [EC:3.1.3.84] :: PFAMS: Macro domain; Cellulose biosynthesis protein BcsO; Domain of unknown function (DUF1768); RNA-binding PUA-like domain of methyltransferase RsmF; Ribosome biogenesis regulatory protein (RRS1) | |
| CBQ Anger Frustration K03767 | + | KO: PPIA; peptidyl-prolyl cis-trans isomerase A (cyclophilin A) [EC:5.2.1.8] :: PFAMS: Cyclophilin type peptidyl-prolyl cis-trans isomerase/CLD; Protein of unknown function (DUF732); FKBP-type peptidyl-prolyl cis-trans isomerase; Protein of unknown function (DUF3830); Nepovirus subgroup A polyprotein | K10598 |
| C Unsafe/Violent Neighborhood K10598 | - | KO: PPIL2, CYC4, CHP60; peptidyl-prolyl cis-trans isomerase-like 2 [EC:5.2.1.8] :: PFAMS: Cyclophilin type peptidyl-prolyl cis-trans isomerase/CLD; Rtf2 RING-finger; Zinc-finger of nitric oxide synthase-interacting protein; U-box domain; Zinc-finger of the MIZ type in Nse subunit | K03767 |
| CBCL Depressive Problems K09773 | + | KO: ppsR; [pyruvate, water dikinase]-phosphate phosphotransferase / [pyruvate, water dikinase] kinase [EC:2.7.4.28 2.7.11.33] :: PFAMS: Kinase/pyrophosphorylase; Biogenesis of lysosome-related organelles complex-1 subunit 2; Domain of unknown function (DUF4276); Chromatin remodelling complex Rsc7/Swp82 subunit; Protein of unknown function (DUF2951) | |
| CBQ Inhibitory Control K02835 | - | KO: prfA, MTRF1, MRF1; peptide chain release factor 1 :: PFAMS: PCRF domain; RF-1 domain; ABC transporter C-terminal domain; MIR domain; Hr1 repeat | |
| CBCL Depressive Problems K16329 | + | KO: psuG; pseudouridylate synthase [EC:4.2.1.70] :: PFAMS: Indigoidine synthase A like protein; Precorrin-8X methylmutase; Thiamine pyrophosphate enzyme; N-terminal TPP binding domain; 3' exoribonuclease family; domain 2; Xylose isomerase-like TIM barrel | K13540 K16328 |
| CBCL Depressive Problems K16328 | + | KO: psuK; pseudouridine kinase [EC:2.7.1.83] :: PFAMS: pfkB family carbohydrate kinase; MarR family; Winged helix-turn-helix DNA-binding; Phosphomethylpyrimidine kinase; HTH domain | K13540 K16329 |
| CBCL Depressive Problems K13252 | + | KO: ptcA; putrescine carbamoyltransferase [EC:2.1.3.6] :: PFAMS: Aspartate/ornithine carbamoyltransferase; Asp/Orn binding domain; Aspartate/ornithine carbamoyltransferase; carbamoyl-P binding domain; PD-(D/E)XK nuclease superfamily; Family 4 glycosyl hydrolase; Root cap | K09251 |
| CBQ Sadness K04458 | - | KO: PTPRR; receptor-type tyrosine-protein phosphatase R [EC:3.1.3.48] :: PFAMS: Protein-tyrosine phosphatase; Dual specificity phosphatase; catalytic domain; Tyrosine phosphatase family; Inositol hexakisphosphate; Protein of unknown function (DUF1999) | |
| CBQ Anger Frustration K02784 | + | KO: PTS-HPR.PTSH, ptsH; phosphocarrier protein HPr :: PFAMS: PTS HPr component phosphorylation site; Domain of unknown function (DUF1906); Glucodextranase; domain B; Phosphoribosylglycinamide synthetase; C domain; Domain of unknown function (DUF4417) | |
| CBQ Inhibitory Control K09684 | - | KO: pucR; purine catabolism regulatory protein :: PFAMS: Purine catabolism regulatory protein-like family; PucR C-terminal helix-turn-helix domain; Bacterial regulatory protein; Fis family; Bacterial regulatory helix-turn-helix protein; lysR family; Protein of unknown function (DUF2089) | K00087 |
| CBCL Internalizing Behavior K08018 | - | KO: RAPGEF2, PDZGEF1; Rap guanine nucleotide exchange factor 2 :: PFAMS: RasGEF domain; RasGEF N-terminal motif; PDZ domain (Also known as DHR or GLGF); Ras association (RalGDS/AF-6) domain; Cyclic nucleotide-binding domain | |
| CBQ Inhibitory Control K06218 | - | KO: relE, stbE; mRNA interferase RelE/StbE :: PFAMS: ParE toxin of type II toxin-antitoxin system; parDE; ParE-like toxin of type II bacterial toxin-antitoxin system; Phage derived protein Gp49-like (DUF891); YoeB-like toxin of bacterial type II toxin-antitoxin system; Cell division protein SepF | K09764 K02663 K02454 |
| CBQ Impulsivity K15531 | - | KO: rexA; oligosaccharide reducing-end xylanase [EC:3.2.1.156] :: PFAMS: Glycosyl hydrolases family 8; Carbohydrate family 9 binding domain-like; Polysaccharide deacetylase; Dockerin type I repeat; AAA domain | K01224 K01220 K06113 K09963 K09955 K05988 K05991 K01223 K11931 K01187 K15924 |
| CBCL Anxiety Problems K12454 | - | KO: rfbE; CDP-paratose 2-epimerase [EC:5.1.3.10] :: PFAMS: NAD dependent epimerase/dehydratase family; GDP-mannose 4;6 dehydratase; 3-beta hydroxysteroid dehydrogenase/isomerase family; Male sterility protein; RmlD substrate binding domain | |
| CBQ Anger Frustration K11939 | + | KO: rhtA; inner membrane transporter RhtA :: PFAMS: EamA-like transporter family; Nucleotide-sugar transporter; Triose-phosphate Transporter family; Bacteriophage holin family HP1; Small Multidrug Resistance protein | K03327 K11742 K03577 |
| CBCL Depressive Problems K12700 | + | KO: rihC; non-specific riboncleoside hydrolase [EC:3.2.-.-] :: PFAMS: Inosine-uridine preferring nucleoside hydrolase; Type-F conjugative transfer system pilin assembly protein; Uncharacterized protein conserved in bacteria (DUF2218); Glycosyl transferase family 41; TcdA/TcdB pore forming domain | |
| CBCL Depressive Problems K03471 | + | KO: rnhC; ribonuclease HIII [EC:3.1.26.4] :: PFAMS: Ribonuclease HII; Domain of unknown function (DUF3378); Transcription factor TFIID (or TATA-binding protein; TBP); Protein of unknown function (DUF4497); Domain of unknown function (DUF4830) | |
| CBQ Fear K02553 | + | KO: rraA, menG; regulator of ribonuclease activity A :: PFAMS: Aldolase/RraA; MOSC N-terminal beta barrel domain; Metallo-beta-lactamase superfamily; PA domain; Pre-toxin domain with VENN motif | |
| amily Separation/Social Services K14744 | + | KO: rzpD; prophage endopeptidase [EC:3.4.-.-] :: PFAMS: Bacteriophage Rz lysis protein; Protein of unknown function (DUF2681); Phage minor structural protein GP20; Phage shock protein B; HAUS augmin-like complex subunit 3 | K16589 K03746 |
| CBQ Anger Frustration K04525 | + | KO: SERPINA; serpin peptidase inhibitor, clade A :: PFAMS: Serpin (serine protease inhibitor); Protein of unknown function (DUF531); MukF winged-helix domain; 3' exoribonuclease family; domain 1; Bacteriophage protein GP30.3 | |

| | | | |
|---|---|---|---|
| CBCL Internalizing Behavior K11428 | - | KO: SETD8; histone-lysine N-methyltransferase SETD8 [EC:2.1.1.43] :: PFAMS: SET domain; Endoplasmic reticulum protein ERp29, C-terminal domain; YL1 nuclear protein; SAF domain; Raftlin | K09186 K11420 |
| CBCL Internalizing Behavior K08172 | + | KO: shiA; MFS transporter, MHS family, shikimate and dehydroshikimate transport protein :: PFAMS: Major Facilitator Superfamily; Sugar (and other) transporter; MFS/sugar transport protein; ATP synthase j chain; Domain of unknown function DUF21 | K08369 K08166 K08144 |
| Income to Needs K15710 | - | KO: SHPRH; E3 ubiquitin-protein ligase SHPRH [EC:3.6.4.- 2.3.2.27] :: PFAMS: SNF2 family N-terminal domain; Zinc finger, C3HC4 type (RING finger); Helicase conserved C-terminal domain; Ring finger domain; RING-type zinc-finger | K15712 K10639 K10661 |
| Income to Needs K14708 | + | KO: SLC26A11; solute carrier family 26 (sodium-independent sulfate anion transporter), member 11 :: PFAMS: Sulfate permease family; STAS domain; Molybdate transporter of MFS superfamily; Stannin transmembrane; Major Facilitator Superfamily | K10820 K10975 K10974 K07243 K02526 K14701 |
| Income to Needs K14701 | + | KO: SLC26A2, DTD; solute carrier family 26 (sulfate anion transporter), member 2 :: PFAMS: Sulfate permease family; STAS domain; Putative heavy-metal chelation; Domain of unknown function (DUF3488); Protein of unknown function (DUF2956) | K10820 K10975 K10974 K07243 K14708 K02526 |
| CBCL Depressive Problems K08144 | + | KO: SLC2A6, GLUT6; MFS transporter, SP family, solute carrier family 2 (facilitated glucose transporter), member 6 :: PFAMS: Sugar (and other) transporter; Major Facilitator Superfamily; Uncharacterised MFS-type transporter YbfB; MFS_1 like family; Fungal trichothecene efflux pump (TRI12) | K08369 K08166 K08172 |
| CBCL Depressive Problems K14716 | + | KO: SLC39A10, ZIP10; solute carrier family 39 (zinc transporter), member 10 :: PFAMS: ZIP Zinc transporter; Putative manganese efflux pump; Kita-kyushu lung cancer antigen 1; Transmembrane region of lysyl-tRNA synthetase; Domain of unknown function (DUF4203) | K16846 K02074 K15726 K15727 |
| CBQ Impulsivity K12486 | + | KO: SMAP; stromal membrane-associated protein :: PFAMS: Putative GTPase activating protein for Arf; C2 domain; Photosystem P840 reaction centre protein PscD; Nuclear RNA-splicing-associated protein; Integral membrane protein DUF106 | |
| CBCL Depressive Problems K08293 | + | KO: SMK1; sporulation-specific mitogen-activated protein kinase SMK1 [EC:2.7.11.24] :: PFAMS: Protein kinase domain; Protein tyrosine kinase; Haspin like kinase domain; Kinase-like; Lipopolysaccharide kinase (Kdo/WaaP) family |  |
| CBCL Depressive Problems K13639 | + | KO: soxR; MerR family transcriptional regulator, redox-sensitive transcriptional activator SoxR :: PFAMS: MerR; DNA binding; MerR HTH family regulatory protein; MerR family regulatory protein; Helix-turn-helix domain; Helix-turn-helix | |
| CBQ Inhibitory Control K06390 | - | KO: spoIIIAA; stage III sporulation protein AA :: PFAMS: AAA domain; NTPase; ATPase family associated with various cellular activities (AAA); Type II/IV secretion system protein; Bacterial TniB protein | K06393 K06387 |
| CBQ Inhibitory Control K06393 | - | KO: spoIIIAD; stage III sporulation protein AD :: PFAMS: Stage III sporulation protein AC/AD protein family; CAAD domains of cyanobacterial aminoacyl-tRNA synthetase; LysE type translocator; Nuclear pore complex component; Bacteriophage T holin | K06390 K06387 |
| CBQ Inhibitory Control K06387 | - | KO: spoIIR; stage II sporulation protein R :: PFAMS: Stage II sporulation protein R (spore_II_R); Stage III sporulation protein AF (Spore_III_AF); Domain of unknown function (DUF966); Domain of unknown function (DUF4382); Presenilin | K06390 K06393 |
| CBQ Impulsivity K00299 | + | KO: ssuE; FMN reductase [EC:1.5.1.38] :: PFAMS: NADPH-dependent FMN reductase; Flavodoxin-like fold; Flavodoxin domain; Flavodoxin; Protein-tyrosine phosphatase receptor IA-2 | |
| CBCL Depressive Problems K03316 | + | KO: TC.CPA1; monovalent cation:H+ antiporter, CPA1 family :: PFAMS: Sodium/hydrogen exchanger family; Cyclic nucleotide-binding domain; Alkali metal cation/H+ antiporter Nha1 C terminus; Zn-finger in ubiquitin-hydrolases and other protein; Phage late-transcription coactivator | |
| CBQ Inhibitory Control K02014 | + | KO: TC.FEV.OM; iron complex outermembrane recepter protein :: PFAMS: TonB dependent receptor; TonB-dependent Receptor Plug Domain; Outer membrane protein beta-barrel family; CarboxypepD_reg-like domain; Carboxypeptidase regulatory-like domain | |
| CBQ Anger Frustration K03327 | - | KO: TC.MATE, SLC47A, norM, mdtK, dinF; multidrug resistance protein, MATE family :: PFAMS: MatE; Polysaccharide biosynthesis C-terminal domain; Tyrosyl-DNA phosphodiesterase; MviN-like protein; Family of unknown function (DUF5389) | K11742 K11939 K03577 |
| amily Separation/Social Services K16051 | + | KO: tesI; 3-oxo-5alpha-steroid 4-dehydrogenase [EC:1.3.99.5] :: PFAMS: FAD binding domain; FAD dependent oxidoreductase; Pyridine nucleotide-disulphide oxidoreductase; HI0933-like protein; Thi4 family | |
| CBQ Impulsivity K04075 | + | KO: tilS, mesJ; tRNA(Ile)-lysidine synthase [EC:6.3.4.19] :: PFAMS: PP-loop family; TilS substrate C-terminal domain; TilS substrate binding domain; Phosphoadenosine phosphosulfate reductase family; Queuosine biosynthesis protein QueC | |
| CBQ Impulsivity K03699 | + | KO: tlyC; putative hemolysin :: PFAMS: Domain of unknown function DUF21; Transporter associated domain; CBS domain; Colicin V production protein; Uncharacterized protein family UPF0016 | |
| CBQ Anger Frustration K07442 | + | KO: TRM61, GCD14; tRNA (adenine57-N1/adenine58-N1)-methyltransferase catalytic subunit [EC:2.1.1.219 2.1.1.220] :: PFAMS: tRNA methyltransferase complex GCD14 subunit; Methyltransferase domain; tRNA methyltransferase complex GCD14 subunit N-term; Protein-L-isoaspartate(D-aspartate) O-methyltransferase (PCMT); Ribosomal RNA adenine dimethylase | K00599 K15471 |
| CBQ Inhibitory Control K06173 | - | KO: truA, PUS1; tRNA pseudouridine38-40 synthase [EC:5.4.99.12] :: PFAMS: tRNA pseudouridine synthase; Uncharacterized protein conserved in bacteria (DUF2344); Acid Phosphatase; Hydroxyacylglutathione hydrolase C-terminus; WVELL protein | |
| CBQ Anger Frustration K16124 | + | KO: tycC; tyrocidine synthetase III :: PFAMS: Condensation domain; AMP-binding enzyme; Phosphopantetheine attachment site; AMP-binding enzyme C-terminal domain; Thioesterase domain | K15665 K15658 K15660 K16130 K12239 K16120 K16129 K16416 K05914 |
| CBQ Fear K12836 | - | KO: U2AF1; splicing factor U2AF 35 kDa subunit :: PFAMS: Zinc finger C-x8-C-x5-C-x3-H type (and similar); RNA recognition motif. (a.k.a. RRM, RBD, or RNP domain); Torus domain; Zinc-finger containing family; Nup53/35/40-type RNA recognition motif | |
| CBCL Depressive Problems K09273 | - | KO: UBTF; upstream-binding transcription factor :: PFAMS: HMG (high mobility group) box; HMG-box domain; HMG (high mobility group) box 5; CHDNT (NUC034) domain; Domain of unknown function (DUF4473) | |
| CBQ Inhibitory Control K03502 | - | KO: umuC; DNA polymerase V :: PFAMS: impB/mucB/samB family; impB/mucB/samB family C-terminal domain; Domain of unknown function (DUF4113); IMS family HHH motif; Fingers domain of DNA polymerase lambda | |
| CBQ Anger Frustration K03648 | - | KO: UNG, UDG; uracil-DNA glycosylase [EC:3.2.2.27] :: PFAMS: Uracil DNA glycosylase superfamily; 3-alpha domain; Domain of unknown function (DUF1738); Toxin with a H; D/N and C signature; Proteinaceous host-selective toxin ToxA |  |
| CBCL Aggressive Behavior K03702 | + | KO: uvrB; excinuclease ABC subunit B :: PFAMS: Ultra-violet resistance protein B; Helicase conserved C-terminal domain; AAA domain; Type III restriction enzyme, res subunit; UvrB/uvrC motif | |
| CBQ Impulsivity K04613 | - | KO: V2R; vomeronasal 2 receptor :: PFAMS: 7 transmembrane sweet-taste receptor of 3 GCPR; Receptor family ligand binding region; Nine Cysteines Domain of family 3 GPCR; Periplasmic binding protein; Putative ephrin-receptor like | |
| CBCL Depressive Problems K16711 | + | KO: wcaM; colanic acid biosynthesis protein WcaM :: PFAMS: TAT (twin-arginine translocation) pathway signal sequence; Pectate lyase superfamily protein; Protein of unknown function (DUF861); Ribosomal protein L20; Cupin | |
| EC Unsafe/Violent Neighborhood K03815 | + | KO: xapA; xanthosine phosphorylase [EC:2.4.2.-] :: PFAMS: Phosphorylase superfamily; Ribulose bisphosphate carboxylase large chain, N-terminal domain; Uncharacterised protein family (UPF0227); Fungal protease inhibitor; K cyclin, C terminal | |
| CBCL Depressive Problems K06909 | + | KO: xtmB; phage terminase large subunit :: PFAMS: Phage terminase large subunit; Terminase RNAseH like domain; Terminase-like family; Terminase RNaseH-like domain; Helicase | |
| CBQ Impulsivity K15924 | - | KO: xynC; glucuronoarabinoxylan endo-1,4-beta-xylanase [EC:3.2.1.136] :: PFAMS: Glycosyl hydrolase family 30 beta sandwich domain; Glycosyl hydrolase family 30 TIM-barrel domain; Carbohydrate binding module (family 6); O-Glycosyl hydrolase family 30; Glycosyl hydrolase family 59 | K01224 K01220 K06113 K09963 K09955 K05988 K05991 K01223 K11931 K15531 K01187 |
| CBCL Anxious Depressed K06677 | - | KO: YCS4, CNAP1, CAPD2; condensin complex subunit 1 :: PFAMS: non-SMC mitotic condensation complex subunit 1; non-SMC mitotic condensation complex subunit 1, N-term; HEAT repeat; HEAT repeats; Nuclear condensing complex subunits, C-term domain | |
| CBCL Aggressive Behavior K16066 | + | KO: ydfG; 3-hydroxy acid dehydrogenase / malonic semialdehyde reductase [EC:1.1.1.381 1.1.1.-] :: PFAMS: short chain dehydrogenase; Enoyl-(Acyl carrier protein) reductase; KR domain; NAD dependent epimerase/dehydratase family; NAD(P)H-binding | |
| CBCL Depressive Problems K03492 | + | KO: ydhQ; GntR family transcriptional regulator :: PFAMS: UTRA domain; Bacterial regulatory proteins, gntR family; Helix-turn-helix domain; MarR family; HTH domain | |
| CBCL Depressive Problems K08369 | + | KO: ydjE; MFS transporter, putative metabolite:H+ symporter :: PFAMS: Major Facilitator Superfamily; Sugar (and other) transporter; Uncharacterised MFS-type transporter YbfB; MFS_1 like family; Organic Anion Transporter Polypeptide (OATP) family | K08166 K08172 K08144 |
| CBCL Depressive Problems K16326 | + | KO: yeiL; CRP/FNR family transcriptional regulator, putaive post-exponential-phase nitrogen-starvation regulator :: PFAMS: Cyclic nucleotide-binding domain; Crp-like helix-turn-helix domain; Winged helix-turn helix; Winged helix DNA-binding domain; Cysteine-rich CWC | K10914 |
| CBQ Fear K03214 | + | KO: yfiF, trmG; RNA methyltransferase, TrmH family [EC:2.1.1.-] :: PFAMS: SpoU rRNA Methylase family; RNA 2'-O ribose methyltransferase substrate binding; GTP cyclohydrolase I feedback regulatory protein (GFRP); Glyoxalase/Bleomycin resistance protein/Dioxygenase superfamily; Glycine/sarcosine/betaine reductase component B subunits | K05810 |
| LEC Family Illness/Injury K05810 | - | KO: yfiH; polyphenol oxidase [EC:1.10.3.-] :: PFAMS: Multi-copper polyphenol oxidoreductase laccase; Septum formation inhibitor MinC, C-terminal domain; Lipase (class 3); CheC-like family; Translation initiation factor IF-3, N-terminal domain | K03214 |
| CBQ Inhibitory Control K09299 | + | KO: ZEB1_2; zinc finger homeobox protein 1/2 :: PFAMS: Zinc-finger double domain; Zinc finger, C2H2 type; C2H2-type zinc finger; Homeobox domain; Zinc-finger double-stranded RNA-binding | |
| CBCL Depressive Problems K07803 | + | KO: zraP; zinc resistance-associated protein :: PFAMS: Heavy-metal resistance; LTXXQ motif family protein; Protein of unknown function (DUF3584); Biogenesis of lysosome-related organelles complex-1 subunit 2; Hyaluronan mediated motility receptor C-terminal | |
| CBCL Depressive Problems K03274 | + | MODS: ADP-L-glycero-D-manno-heptose biosynthesis | |
| CBQ Fear K01879 | + | MODS: Aminoacyl-tRNA biosynthesis, prokaryotes | K01889 |
| CBQ Impulsivity K01889 | + | MODS: Aminoacyl-tRNA biosynthesis, prokaryotes; Aminoacyl-tRNA biosynthesis, eukaryotes | K01879 |
| CBCL Depressive Problems K03079 | + | MODS: Ascorbate degradation, ascorbate => D-xylulose-5P | K00963 |
| CBCL Internalizing Behavior K13571 | + | MODS: Bacterial proteasome | K03064 |
| amily Separation/Social Services K10865 | + | MODS: BRCA1-associated genome surveillance complex (BASC); MRN complex; MRX complex | |

| | | | |
|---|---|---|---|
| CBQ Anger Frustration K04771 | + | MODS: Cationic antimicrobial peptide (CAMP) resistance, envelope protein folding and degrading factors DegP and DsbA | |
| LEC Family Illness/Injury K06632 | + | MODS: Cell cycle - G2/M transition; Cell cycle - G2/M transition | K06662 |
| CBCL Externalizing Behavior K01135 | - | MODS: Chondroitin sulfate degradation; Dermatan sulfate degradation | |
| CBQ Inhibitory Control K00024 | + | MODS: Citrate cycle, second carbon oxidation, 2-oxoglutarate => oxaloacetate; Glyoxylate cycle; Dicarboxylate-hydroxybutyrate cycle; Reductive citrate cycle (Arnon-Buchanan cycle); Incomplete reductive citrate cycle, acetyl-CoA => oxoglutarate; CAM (Crassulacean acid metabolism), dark; Formaldehyde assimilation, serine pathway; Citrate cycle (TCA cycle, Krebs cycle); Methylaspartate cycle | K01595 K00174 |
| Income to Needs K00174 | - | MODS: Citrate cycle, second carbon oxidation, 2-oxoglutarate => oxaloacetate; Reductive citrate cycle (Arnon-Buchanan cycle); Incomplete reductive citrate cycle, acetyl-CoA => oxoglutarate; Citrate cycle (TCA cycle, Krebs cycle) | K00024 |
| CBCL Depressive Problems K01922 | + | MODS: Coenzyme A biosynthesis, pantothenate => CoA | |
| CBQ Anger Frustration K01697 | + | MODS: Cysteine biosynthesis, homocysteine + serine => cysteine; Methionine degradation | K08965 |
| CBCL Depressive Problems K02258 | + | MODS: Cytochrome c oxidase | K00256 K07112 |
| amily Separation/Social Services K01631 | + | MODS: D-galactonate degradation, De Ley-Doudoroff pathway, D-galactonate => glycerate-3P | K03738 |
| amily Separation/Social Services K08323 | + | MODS: D-Glucuronate degradation, D-glucuronate => pyruvate + D-glyceraldehyde 3P | K00161 |
| Income to Needs K04561 | + | MODS: Denitrification, nitrate => nitrogen | K02567 |
| LEC Family Illness/Injury K16202 | + | MODS: Dipeptide transport system, Firmicutes | |
| CBCL Depressive Problems K00362 | + | MODS: Dissimilatory nitrate reduction, nitrate => ammonia | K02567 |
| CBQ Inhibitory Control K02322 | - | MODS: DNA polymerase II complex, archaea | |
| LEC Family Illness/Injury K03227 | + | MODS: EHEC/EPEC pathogenicity signature, T3SS and effectors; Type III secretion system; Xanthomonas spp. pathogenicity signature, T3SS and effectors | K03220 |
| CBQ Anger Frustration K02132 | + | MODS: F-type ATPase, eukaryotes | |
| CBCL Depressive Problems K11212 | + | MODS: F420 biosynthesis | |
| CBQ Inhibitory Control K01785 | + | MODS: Galactose degradation, Leloir pathway, galactose => alpha-D-glucose-1P | K04618 |
| C Unsafe/Violent Neighborhood K11131 | + | MODS: H/ACA ribonucleoprotein complex | |
| CBQ Impulsivity K06225 | - | MODS: Hedgehog signaling | |
| CBQ Fear K00228 | + | MODS: Heme biosynthesis, glutamate => protoheme/siroheme | K05612 K00467 K10005 |
| CBQ Fear K02841 | + | MODS: Lipopolysaccharide biosynthesis, inner core => outer core => O-antigen | |
| CBQ Anger Frustration K00164 | + | MODS: Lysine degradation, lysine => saccharopine => acetoacetyl-CoA; Citrate cycle, second carbon oxidation, 2-oxoglutarate => oxaloacetate; Citrate cycle (TCA cycle, Krebs cycle) | |
| CBQ Anger Frustration K13788 | + | MODS: Methanogenesis, acetate => methane; Phosphate acetyltransferase-acetate kinase pathway, acetyl-CoA => acetate | K04480 K14083 |
| CBQ Inhibitory Control K04480 | - | MODS: Methanogenesis, methanol => methane | K13788 K14038 |
| CBQ Anger Frustration K08965 | + | MODS: Methionine salvage pathway | K01697 |
| CBCL Depressive Problems K03577 | + | MODS: Multidrug resistance, efflux pump AcrAB-TolC/SmeDEF | K03327 K11939 K11742 |
| CBQ Inhibitory Control K01001 | - | MODS: N-glycan precursor biosynthesis | |
| amily Separation/Social Services K07151 | + | MODS: N-glycosylation by oligosaccharyltransferase | |
| Income to Needs K10094 | + | MODS: Nickel transport system | |
| CBQ Anger Frustration K03738 | + | MODS: Non-phosphorylative Entner-Doudoroff pathway, gluconate/galactonate => glycerate | K01631 |
| LEC Family Illness/Injury K08678 | + | MODS: Nucleotide sugar biosynthesis, eukaryotes | K00963 K01791 |
| CBQ Anger Frustration K00963 | + | MODS: Nucleotide sugar biosynthesis, eukaryotes; Nucleotide sugar biosynthesis, prokaryotes; Ascorbate biosynthesis, animals, glucose-1P => ascorbate; Nucleotide sugar biosynthesis, glucose => UDP-glucose | K03079 K08678 K01791 |
| CBQ Inhibitory Control K01791 | + | MODS: Nucleotide sugar biosynthesis, prokaryotes | K08678 K00963 |
| CBCL Emotionally Reactive K12743 | + | MODS: Penicillin biosynthesis, aminoadipate + cycteine + valine => penicillin; Cephamycin C biosynthesis, aminoadipate + cycteine + valine => cephamycin C | |
| CBCL Depressive Problems K00981 | - | MODS: Phosphatidylethanolamine (PE) biosynthesis, PA => PS => PE | |
| LEC Family Illness/Injury K03064 | - | MODS: Proteasome, 19S regulatory particle (PA700) | K13571 |
| CBCL Depressive Problems K02788 | + | MODS: PTS system, lactose-specific II component | K10984 K02799 K02810 |
| CBQ Anger Frustration K02799 | + | MODS: PTS system, mannitol-specific II component | K02788 K10984 K02810 |
| CBQ Anger Frustration K02810 | + | MODS: PTS system, sucrose-specific II component | K02788 K02799 K10984 |
| CBCL Depressive Problems K00087 | + | MODS: Purine degradation, xanthine => urea | K09684 |
| CBCL Depressive Problems K06998 | + | MODS: Pyocyanine biosynthesis, chorismate => pyocyanine | K02548 K01658 K04092 |
| CBQ Inhibitory Control K00275 | + | MODS: Pyridoxal biosynthesis, erythrose-4P => pyridoxal-5P | |
| CBQ Anger Frustration K01494 | + | MODS: Pyrimidine deoxyribonuleotide biosynthesis, CDP/CTP => dCDP/dCTP,dTDP/dTTP | |
| CBCL Depressive Problems K00161 | + | MODS: Pyruvate oxidation, pyruvate => acetyl-CoA | K08323 |
| CBQ Anger Frustration K13409 | + | MODS: RaxAB-RaxC type I secretion system | |
| C Unsafe/Violent Neighborhood K02942 | + | MODS: Ribosome, eukaryotes | K02936 |
| CBQ Inhibitory Control K14988 | - | MODS: SalK-SalR two-component regulatory system | |
| CBCL Aggressive Behavior K15554 | + | MODS: Sulfonate transport system | |
| CBCL Depressive Problems K02062 | + | MODS: Thiamine transport system | |
| LEC Family Illness/Injury K00872 | + | MODS: Threonine biosynthesis, aspartate => homoserine => threonine | |
| LEC Family Illness/Injury K01658 | - | MODS: Tryptophan biosynthesis, chorismate => tryptophan | K16314 K04092 K01556 K01593 |
| CBQ Anger Frustration K03220 | + | MODS: Type III secretion system | K03227 |
| CBCL Anxious Depressed K04092 | + | MODS: Tyrosine biosynthesis, chorismate => tyrosine; Phenylalanine biosynthesis, chorismate => phenylalanine | K16314 K01658 K06998 K02548 K01593 |
| CBQ Impulsivity K02120 | + | MODS: V/A-type ATPase, prokaryotes | |
| CBQ Impulsivity K10243 | + | PFAMS: ABC transporter; AAA domain; TOBE domain; AAA ATPase domain; AAA domain; putative AbiEii toxin; Type IV TA system | K02526 K10975 K10974 K07243 K14708 K14701 K10820 |
| CBQ Anger Frustration K01249 | + | PFAMS: Inosine-uridine preferring nucleoside hydrolase; Uracil DNA glycosylase superfamily; DNA alkylation repair enzyme; ADP-ribosylglycohydrolase; snoRNA binding domain; fibrillarin | K03631 K03515 K03660 K10839 |
| CBCL Emotionally Reactive K05315 | - | PFAMS: Ion transport protein; Voltage-dependent L-type calcium channel; IQ-associated; Voltage gated calcium channel IQ domain; Polycystin cation channel; Voltage-gated calcium channel subunit alpha; C-term | K12954 K01560 K05967 05315 |
| CBQ Anger Frustration K00397 | + | PFAMS: Molybdopterin oxidoreductase; Phosphoadenosine phosphosulfate reductase family; Molydopterin dinucleotide binding domain; Nitrate reductase delta subunit; Molybdopterin oxidoreductase Fe4S4 domain | K08357 K00183 |
| CBQ Impulsivity K07185 | + | PFAMS: TspO/MBR family; Cytochrome c-type biogenesis protein CcmF C-terminal; Domain of unknown function (DUF368); Tripartite tricarboxylate transporter TctB family; Predicted membrane protein (DUF2070) | |
| CBCL Depressive Problems K01632 | + | PFAMS: XFP N-terminal domain; D-xylulose 5-phosphate/D-fructose 6-phosphate phosphoketolase; XFP C-terminal domain; Thiamine pyrophosphate enzyme; C-terminal TPP binding domain; Phosphotransferase enzyme family | |
